# A Unique Polyfunctional IL-33-responsive ST2⁺ NK cell State Potentiates Antitumor Immunity and Response to Immunotherapy

**DOI:** 10.1101/2023.02.14.528486

**Authors:** Anaïs Eberhardt, Elena Blanc, Valentin Picant, Aurélien Voissière, Lara Revol-Bauz, Timothée Casini, Vincent Alcazer, Dominique Poujol, Laurie Tonon, Raphaël Schneider, Emilie Picard, Yamila Rocca, Maude Ardin, Fanny Onodi, Cyril Degletagne, Céline Rodriguez, Lyvia Moudombi, Benjamin Estavoyer, Emily Charrier, Xi Wang, Ana Stojanovic, Tilman Rau, Olivier Tredan, Isabelle Treilleux, Giorgio Trinchieri, Eric Vivier, Jenny Valladeau-Guilemond, Antoine Marçais, Thierry Walzer, Philippe Krebs, Adelheid Cerwenka, Margaux Hubert, Christophe Caux, Marie-Cécile Michallet, Nathalie Bendriss-Vermare

## Abstract

Despite increasing recognition of NK cell diversity, their functional heterogeneity and its relevance to cancer remain incompletely understood. Here, we show that IL-12 promotes the emergence of a distinct ST2/IL33R+ NK cell state poised for robust activation by IL-33. Human ST2⁺ NK cells exhibit a unique transcriptional program, an intermediate differentiation state between CD56^bright^ and CD56^dim^ NK cells, and enhanced polyfunctionality combining proliferation, cytokine production, and cytotoxicity. ST2⁺ NK cells were identified across human and murine tumor datasets. In murine breast cancer models, IL-33 and IL-12 cooperated to elicit potent antitumor responses dependent on IFN-γ and ST2⁺ NK cells. In cancer patients, IL-12 unleashed tumor-infiltrating ST2⁺ NK cells’ potential to produce IFN-γ in response to IL-33, an *IL33^hi^-NK*^hi^ score was associated with prolonged survival, and ST2⁺ NK cells correlated with improved immunotherapy response. Collectively, these findings identify ST2⁺ NK cells as a therapeutically actionable NK cell state promoting antitumor immunity.

**One sentence summary:** The IL-33/IL-33R(ST2)/NK cell axis is a key determinant of cancer immunity and response to immunotherapy.

**SIGNIFICANCE:** NK cell functional diversity in cancer remains incompletely understood. Integrated transcriptional and functional analyses identify a unique IL-33-responsive ST2+ NK cell state representing an intermediate differentiation stage with strong polyfunctional anti-tumor activity and favorable prognostic value in solid tumors. IL-33 therefore represents a novel therapeutic axis for harnessing NK cells against cancer.

## INTRODUCTION

Natural Killer (NK) cells are major innate effector cells involved in host defense against viral infections and tumors (1), by directly killing target cells but also by supporting effective innate and adaptive immune responses (2). Accordingly, high intratumoral infiltration by NK cells is associated with improved prognosis of cancer patients (3,4). However, chronic exposure to activating signals, presence of inhibitory ligands and immunosuppressive molecules in tumor microenvironment favor NK cells dysfunction, leading to tumor escape (1). In this context, NK cells are emerging as promising targets for cancer therapy with a growing number of therapeutic agents aiming to unleash their full antitumor potential.

NK cells have evolved into a spectrum of effector subsets diverging in terms of phenotype, function, location, and responsiveness to activating stimuli to ensure complementary roles in immune responses (5). Indeed, in humans, CD56^bright^ NK cells are predominant in lymphoid tissues, secrete high levels of cytokines and chemokines including Interferon-gamma (IFN)-γ, Tumor Necrosis Factor-alpha (TNF-α), CCL3/4 (Macrophage Inflammatory Protein 1 α/β), and CCL5 (RANTES) and undergo robust proliferation after cytokine stimulation. Conversely, CD56^dim^ NK cells are the largest circulating population, show a greater cytotoxic ability but a lower cytokine production and proliferation, and express the activating Fc receptor CD16 endowing them with antibody-dependent cellular cytotoxicity (ADCC) function, a central mechanism in certain antibody-based antitumor therapies.

The landscape of NK cell subsets and developmental relationships between them was partially solved, with CD56^dim^ NK cells originating from the differentiation of CD56^bright^ NK cells (6,7) and evidences of phenotypically and functionally distinct intermediate subsets along this developmental trajectory including terminally differentiated CD56^dim^ CD57^+^ (8,9), memory-like/adaptive (10) and hypo-responsive exhausted-like NK cells (11). More recently, single cell transcriptomic (scRNAseq) analyses allowed to precisely dissect the complexity and heterogeneity of NK cell subsets, in physiological (12,13) and tumoral contexts (13–15). NK cell subsets have distinct transcriptional programs that are governed by profound epigenetic modifications and chromatin re-arrangements (16,17). Yet, functional heterogeneity in tumoral context and in response to cytokine stimuli from the tumor microenvironment remains to be fully uncovered. Recent evidence show that a subset of cytotoxic NK cells limits the efficacy of immune checkpoint blockade, by interfering with an efficient cytotoxic T cell response (18,19). Therefore, NK cell transcriptional and functional diversity extends beyond current paradigms, with distinct states differentially shaping antitumor immunity and clinical outcomes, providing new opportunities for therapeutic intervention.

The activation of NK cells depends on the signaling balance between activating and inhibitory receptors. However, numerous cytokines such as Interleukin(IL)-2, IL-12, IL-15, IL-18, and type I IFN (20) also regulate NK cell maturation, activation, and survival by acting alone or in cooperation. The IL-1-familly cytokine IL-33 is constitutively expressed in the nucleus of non-immune cells at epithelial barriers (21). In response to tissue injury, cellular stress, and inflammation, IL-33 is released and functions as an alarmin by activating immune cells expressing its heterodimeric receptor, composed of ST2 (encoded by *IL1RL1* gene) and ILRAcP accessory chain (21). The IL-33/ST2 axis was originally described to support type 2 immunity (22), as well as a subset of regulatory T cells (Tregs) constitutively expressing ST2 and involved in tissue repair (23) and immunomodulation (24). In contrast, IL-33 has later emerged as a regulator of type 1 immune responses during viral infection and chronic immune pathologies in mice, enhancing IFN-γ production in CD4^+^ cells and CD8^+^ T cells that acquire ST2 expression under inflammatory conditions (25–27). In cancer context, IL-33 has revealed opposing effects depending on the target cells and the complex nature of the tumor microenvironment (28). IL-33 can contribute to tumor progression and metastasis through both immune cell-dependent (*i.e*., recruitment of immune suppressive cells, activation of type 2-mediated inflammation) and immune cell-independent (*i.e.,* angiogenesis, cancer cell stemness or proliferation) mechanisms. In contrast, IL-33 can inhibit tumor growth and metastasis by enhancing both adaptive and innate immune responses, supporting promising IL-33-based therapeutic strategies in cancer (28).

Yet, the role of IL-33/ST2 axis in human NK cell subsets biology, especially in the context of cancer, remains scarcely addressed. Here, we evaluated the role of the IL-33/ST2 axis in regulating NK cell biology in physiological and tumor context in human and mouse models. We uncovered an IL-33-responsive human NK cell state defined by IL-12/STAT4-dependent acquisition of ST2 expression and characterized by a distinct functional and transcriptional profile, with high polyfunctional activity and proliferative ability. In cancer context, IL-12 unleashed IL-33-dependent ST2^+^ NK cell activation resulting in a potent antitumor effect relying on IFN-γ secretion in mice. In patients, ST2^+^ NK cells were detected in multiple tumor types, an *NK*^hi^ - *IL33^hi^* score correlates with improved prognosis in several solid cancers, and ST2^+^ NK cells associate with improved response to immune checkpoint blockade. Our results reveal ST2^+^ NK cells as a hitherto unappreciated polyfunctional cell state involved in tumor immune surveillance and response to immunotherapy, and identify IL-33 as a promising NK cell modulator for cancer immunotherapy.

## RESULTS

### IL-12 and IL-33 cooperate to activate polyfunctional human NK cells

As previously reported for mouse NK cells in infectious contexts (27), IL-33 was a strong activator of human NK cells solely when combined with IL-12, as evidenced by a potent increase in IFN-γ production (Fig. 1A; >500-fold, 4-fold increase and comparable levels as compared to IL-12 alone, IL-1α/IL-1β and IL-18 respectively), CD25 and CD69 activation marker expression of (Supplementary Fig. S1A) and cytotoxicity towards K562 target cells (Fig. 1B). IL-33 also synergized with IL-2 (60-fold increase), IL-15 (7-fold increase) for IFN-γ production but not with anti-NKaR agonistic antibodies or K562 target cells (Supplementary Fig. S1B-D). IFN-γ was detected within a few hours upon IL-12 and IL-33 combination, and peaked at 72 h to reach levels comparable to IL-12 and IL-18 combination (Supplementary Fig. 1E). A broader analysis of NK cell secretory profile revealed that the combination of IL-12 and IL-33 induced a wide range of cytokines (IFN-γ, CCL5, TNF-α, CCL3, IL-8, CCL4, IL-6), predominantly produced by CD56^dim^ NK cells. Notably, IL-33 elicited a more inflammatory secretory profile than IL-18, with higher levels of TNF-α, CCL3, and CCL4 (Fig. 1C). Coherently with CD25 expression (Supplementary Fig. S1A), IL-12 and IL-33 combination potentiates NK cell proliferation in response to IL-2 of CD56^dim^ NK cells, which are known as poorly proliferative. Of note, IL-33 was more potent that IL-18 in inducing NK cell proliferation (Fig. 1D). Together, these results indicate that IL-33, in the presence of IL-12, is a potent activator of a subset of NK cells by inducing their polyfunctional activation and strong proliferation.

**Figure 1.**
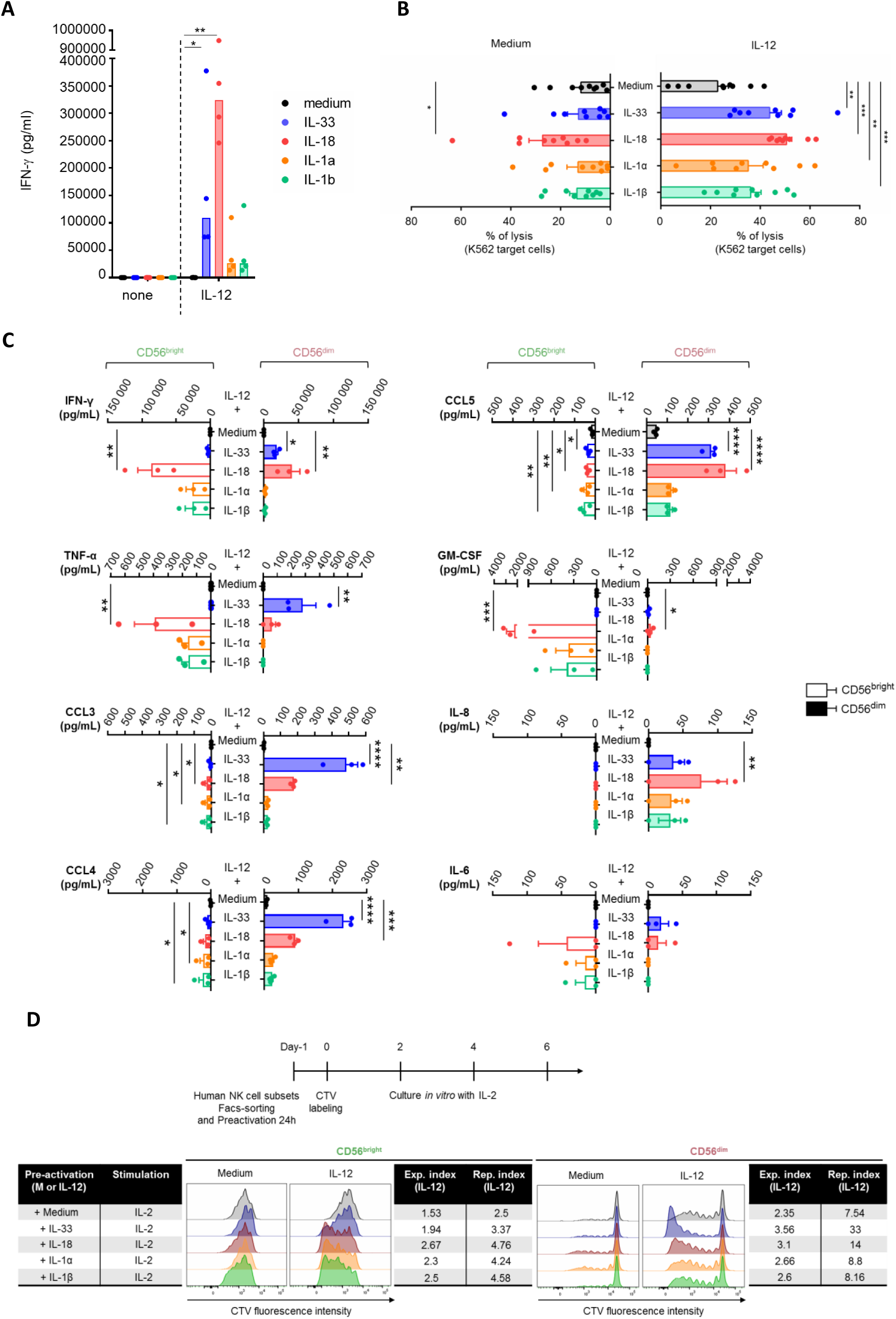
IL-12 and IL-33 cooperate to activate polyfunctional human NK cells. **A,** Quantification of IFN-γ secretion by healthy donors’ blood NK cells upon stimulation with IL-33, IL-18, IL-1α or IL-1β alone (10 ng/mL) or in combination with IL-12 (10 ng/mL) for 24 h. Histogram bars represent the median (n = 4 individual experiments). Friedman test with Dunn’s multiple comparisons test was performed. **B,** NK cells were activated for 24 h with IL-33, IL-18, IL-1α or IL-1β alone (10 ng/mL) or in combination with IL-12 (10 ng/mL) and then co-cultured with calcein-loaded K562 target cells for 4 h at a 5:1 effector to target ratio. Calcein release in the supernatant was quantified by fluorometry to calculate lysis percentage. Results are expressed as mean + SEM (n = 9 individual experiments). One-way repeated measures ANOVA with Dunnett’s multiple comparisons test against levels in medium condition was performed. **C,** Supernatants from FACS-sorted CD56^bright^ (empty histogram bars) and CD56^dim^ (filled histogram bars) NK cells were collected to quantify cytokine and chemokine release by Luminex assay. Of note, IL-2, IL-3, IL-4, IL-5, IL-10, IL-13, IL-17A, IL-22, and G-CSF were not detected (not shown). Results are expressed as mean + SEM (n = 3 individual donors). One-way repeated measures ANOVA with Tukey’s multiple comparisons test was performed. **D,** FACS-sorted CD56^bright^ and CD56^dim^ healthy donors’ blood NK cells were pre-activated as indicated, labeled with CTV, and cultured in low dose IL-2 (100 UI/mL). *In vitro* NK cell proliferation was analyzed at day 3 for CD56^bright^ and day 6 for CD56^dim^ NK cells. Expansion Index (Exp. Index) and Replication Index (Rep. Index) determine the fold-expansion of the overall culture and of the responding cells only, respectively. Data shown are representative of three individual experiments.

### IL-12 licenses a ST2⁺ IL-33-responsive intermediate NK cell state via STAT4

Blocking experiments using specific antagonists of IL-1 family cytokines or receptors demonstrated that ST2 is required for IL33-dependent activation of NK cells (Supplementary Fig. S2A and S2B). Analysis of *IL1RL1* gene (encoding for ST2) mRNA expression showed very low levels in resting human NK cells, but its expression increased upon stimulation with IL-12 alone (Supplementary Fig. S2C) or combined with IL-33 (Supplementary Fig. S2D). In contrast, its co-receptor *IL1RAP* was constitutively expressed in NK cells and moderately upregulated in presence of IL-12 (Supplementary Fig. S2C). Accordingly, ST2 protein was not detected on resting NK cells, but ∼20% of NK cells acquired ST2 surface expression upon IL-12 stimulation (Fig. 2A). IL-12 mainly activates STAT4-dependent transcription programs (29). The treatment of IL-12-activated NK cells with lisofylline, a specific inhibitor of STAT4 phosphorylation (Supplementary Fig. 2E), prevented IL-12 dependent ST2 expression on NK cells (Fig. 2A). *In silico* analysis of the *ST2* promoter identified several potential STAT4 binding sites (Supplementary Fig. S2F), and the binding of STAT4 to the *ST2* promoter in IL-12-stimulated human NK cells was confirmed by chromatin immunoprecipitation (Supplementary Fig. S2G). Next, we aimed to characterize the NK cell subset acquiring ST2 expression upon IL-12 stimulation, hence endowed with IL-33-responsiveness. For this, we first used NF-κB (p65) (30), MAPkinases (p38) (31), and mTOR (S6) (32) signaling pathway activation as surrogates for ST2/IL-33 signaling. In line with ST2 expression, phosphorylation of p65 (Fig. 2B), p38, and S6 (Supplementary Fig. S3A-B) in response to IL-12 and IL-33 co-stimulation was significantly higher in CD56^dim^ compared to CD56^bright^ NK cells. A much greater proportion of CD56^dim^ NK cells as compared to CD56^bright^ NK cells also produced IFN-γ and co-expressed CD25 in response to IL-12 and IL-33 combination (Supplementary Fig. S3C). In agreement, ST2 expression (Fig. 2C) and IFN-γ production (Fig. 2D) were significantly enriched in an immature CD56^dim^ CD57^-^ subset upon IL-12 and IL-33 combination. Collectively, these results demonstrate that IL-12/STAT4 signaling preferentially induces an intermediate ST2^+^ NK cell state enriched in the non-terminally differentiated CD56^dim^ CD57^-^ NK population, allowing for their potent activation in response to IL-33.

**Figure 2.**
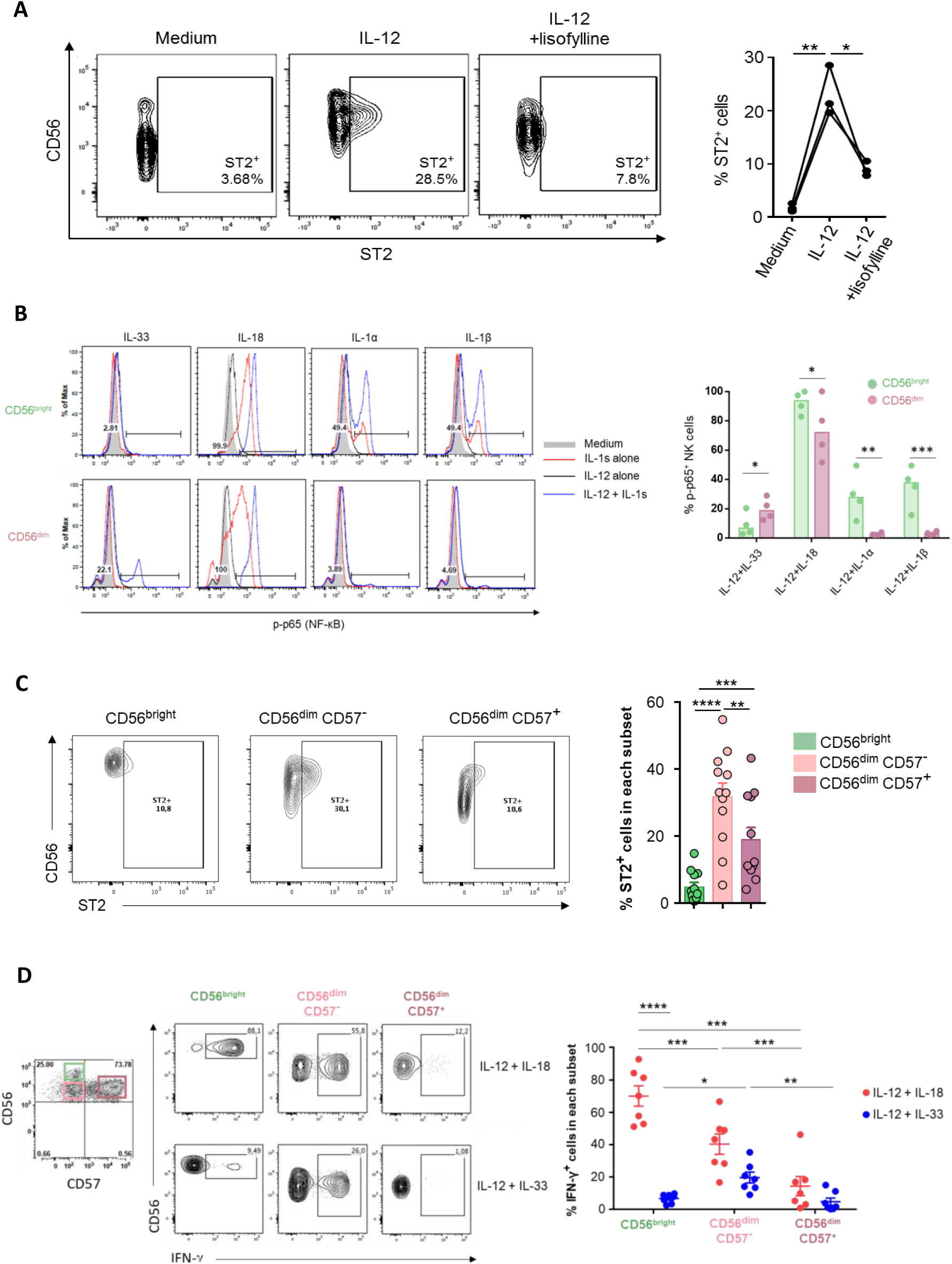
IL-12/STAT4 signaling induces a highly IL-33-responsive ST2^+^ intermediate NK cell state. **A,** NK cells were activated with IL-12 (10 ng/mL) in the presence or not of lisofylline (500 μM) for 24 h prior to ST2 surface expression analysis by flow cytometry. Representative dot plots (left) and quantification (%) (right) of ST2^+^ NK cells after cytokine stimulation. One-way repeated measures ANOVA with Tukey’s multiple comparisons test was performed; n = 3. **B,** FACS-sorted CD56^bright^ and CD56^dim^ healthy donors’ blood NK cells were activated with medium or IL-12 alone for 24 h prior to the addition of IL-33, IL-18, IL-1α or IL-1β, supplemented or not with IL-12. Each cytokine was used at 10ng/mL. p65 (NF-κB) phosphorylation was analyzed 5 min after the addition of IL-1 family cytokines. Representative histogram plots (left) and quantification (%) (right) of p-S6^+^ NK cells after cytokine activation. Histogram bars indicate the median. Two-way ANOVA with Bonferroni’s multiple comparisons test was performed; n = 4 to 5 experiments. **C,** Healthy donors’ blood NK cells were activated with IL-12 (10 ng/mL) for 24 h prior to flow cytometry analysis of ST2 expression on NK cell subsets based on CD16, CD56, and CD57 surface expression. Representative dot plots (left) and quantification (%) (right) of ST2^+^ NK cells in each subset. Results are expressed as mean + SEM (n = 12 experiments). One-way repeated measures ANOVA with Tukey’s multiple comparisons test was performed. **D,** Healthy donors’ blood NK cells were activated with combinations of IL-12 and IL-33 or IL-12 and IL-18 (10 ng/mL of each cytokine) for 24 h prior to flow cytometry analysis of intracellular IFN-γ expression in NK cell subsets based on CD56 and CD57 surface expression. Representative dot plots (left) and quantification (%) (right) of IFN-γ^+^ NK cells in each subset. Results are expressed as mean +/- SEM (n = 7 individual experiments). Two-way ANOVA with Bonferroni’s multiple comparisons test was performed.

### ST2^+^ NK cells display a unique gene regulatory program delineating high activation and polyfunctionality

To characterize the transcriptomic profile of human ST2^+^ NK cells, we performed RNA-seq on eight resting or IL-12 activated NK cell subsets, FACS-sorted based on surface expression of CD56, CD57, and ST2 (Supplementary Fig. S4A-B). Principal Component Analysis (PCA) revealed that ST2^+^ NK cells clustered between ST2^-^ CD56^bright^ and ST2^-^ CD56^dim^ canonical NK cell subsets (Fig. 3A). Focusing on IL-12-activated conditions, 2,399 genes displayed differential expression across all NK cell subsets (Supplementary Fig. S4C). Analysis of genes specifically upregulated in each subset relative to the other two, after pooling CD57^+^ and CD57^-^ cells, allowed us to define transcriptional signatures specific for ST2^-^ CD56^bright^ (418 genes), ST2^-^ CD56^dim^ (396 genes) and a ST2^+^ NK cells signature containing 233 genes encoding for surface proteins (e.g., *IL23R*, *IL1RL1*, *ITGA3)*, secreted proteins (*VEGFA*, *GZMA, IFNG*), and transcription factors (TFs) (*DMRTA2*, *E2F2*, *E2F1*, *EZH2*) (Fig. 3B, Supplementary Fig. S4D and Supplementary tables 1-3). Of note, ST2^+^ NK cells displayed an intermediate enrichment score for both ST2^-^ CD56^bright^ and ST2^-^ CD56^dim^ NK cell signatures (Supplementary Fig. S4E). Importantly, genes encoding canonical TFs, surface markers or soluble mediators of CD56^bright^ or CD56^dim^ NK cells were consistently expressed at intermediate levels in ST2^+^ NK cells (Fig. 3C and Supplementary Fig. S4F). Gene set enrichment analyses using Gene Ontology (GO), Hallmarks, and KEGG databases, as well as NK cell subset-specific cytokine and homing signatures (Supplementary tables 4-7) derived from the literature (33) and our data revealed that ST2^+^ NK cells were enriched in pathways related to cell proliferation and metabolism, and combined features of both CD56^bright^ and CD56^dim^ NK cells, consistent with their enhanced activation and polyfunctionality (Fig. 3D). Of note, the GO term associated with NK cell differentiation was progressively enriched from ST2^-^ CD56^bright^ to ST2^+^ and to ST2^-^ CD56^dim^ NK cells (Fig. 3D). We next performed single-cell RNA-seq (scRNA-seq) to precisely uncover human NK cell diversity and ontogeny upon IL-12 activation (Supplementary Fig. S5A). UMAP visualization revealed four distinct NK cell clusters (Fig. 3E), consistently represented across independent donors (Supplementary Fig. S5B). Notably, cluster 1 represented CD56^bright^ NK cells, while clusters 3 and 4 corresponded to CD56^dim^ and terminally-differentiated NK cells (Fig. 3E, Supplementary Fig. S5C-D). Strikingly, cluster 2 segregated cells overexpressing ST2^+^ NK cells gene signature, including *IL1RL1* gene among the top 10 expressed genes (Fig. 3E-F and Supplementary Fig. S5E). To dissect the ontogeny of ST2^+^ NK cells, we performed trajectory analysis using Slingshot and identified a unique developmental trajectory, with ST2^+^ NK cells emerging along CD56^bright^ differentiation and standing as an intermediate and transitional population between canonical CD56^bright^ and CD56^dim^ NK cells (Fig. 3G). Accordingly, ST2^+^ NK cells signature is progressively acquired across pseudotime upon transition from CD56^bright^ to ST2^+^ NK cells, followed by a gradual loss of ST2-associated gene program upon CD56^dim^ NK cell terminal differentiation (Fig. 3H). Finally, the identification of the top 10 regulons specifically enriched in each cluster using DoRothEA (34) evidenced a profound and progressive remodeling of transcription factor activity upon acquisition of ST2^+^ NK cell program, with an enrichment of a unique set of transcription factors (*ATF6, STAT6, ELK4, NFKB1, RELA, CEBPA, NFATC2, CREB1*) as compared to CD56^bright^ and CD56^dim^ NK cells (Fig. 3I). Altogether, in-depth transcriptomic analyses identify ST2^+^ NK cells as a unique intermediate NK cell state, emerging across NK cell differentiation and endowed with high metabolic and proliferative abilities as well as polyfunctional effector functions.

**Figure 3.**
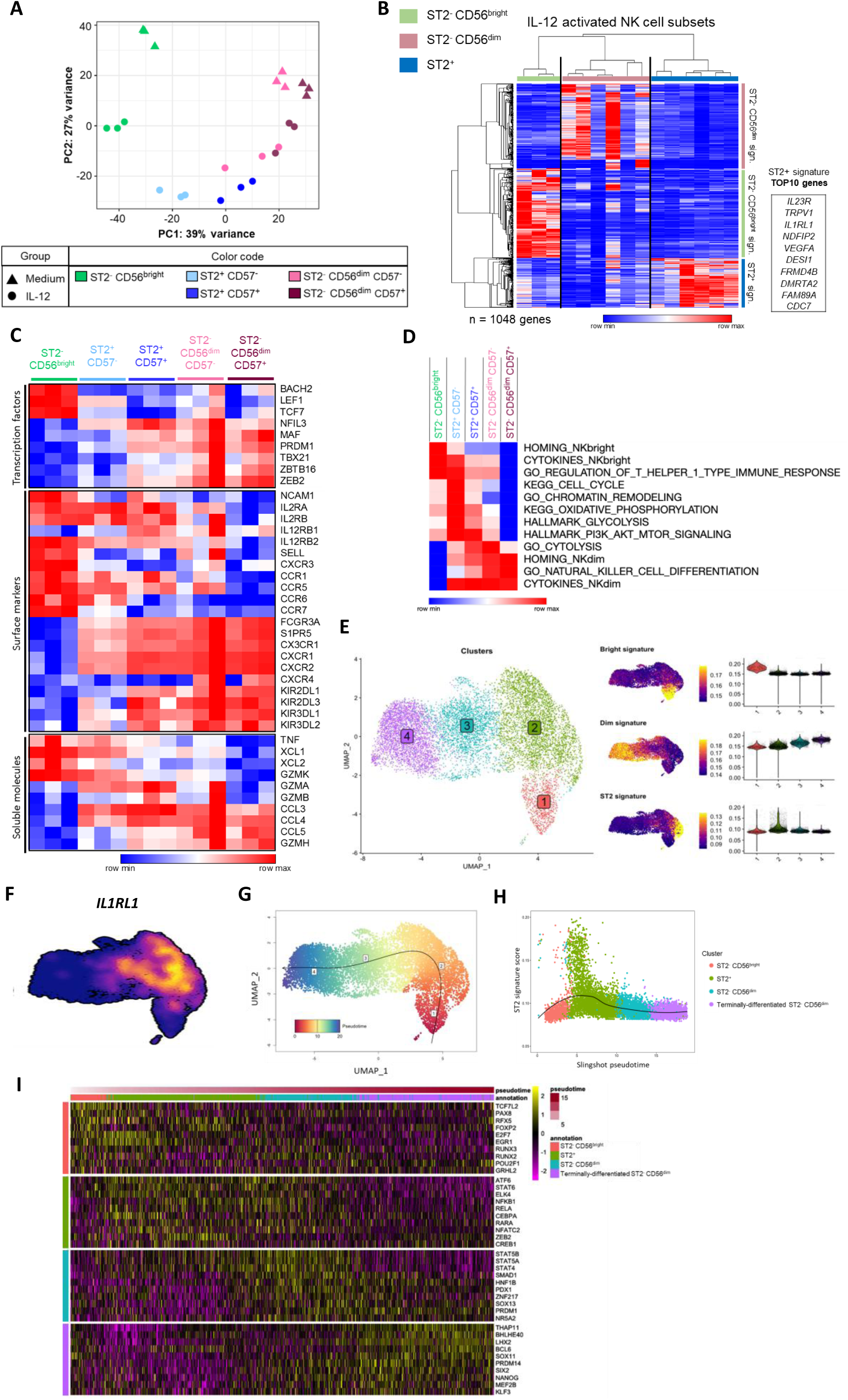
ST2^+^ NK cells represent a unique intermediate state emerging during NK cell differentiation. **A,** Principal component analysis for eight FACS-sorted NK cell subsets based on expression of the top 500 most variant genes obtained from 3 individual healthy donors. **B,** Heatmap representing upregulated genes in each sorted NK cell subset compared to the two other NK cell subsets following IL-12 activation for 24 h (n = 3 individual healthy donors). **C,** Heatmap showing normalized expression for selected genes related to NK cell phenotype and functions. **D,** ssGSEA analysis of selected biological pathways in each sorted NK cell subset following IL-12 activation for 24 h (n = 3 individual healthy donors). **E,** UMAP representation of blood NK cells isolated from 2 independent healthy donors with distinct clusters identified by unsupervised hierarchical clustering based on scRNA-seq analysis following activation with IL-12 for 48 h. Gene signatures specifically enriched in CD56^dim^, CD56^bright^ and ST2^+^ NK cells identified from our bulk RNAseq analysis are projected (middle panels). Their relative expression in each cluster is represented using violin plots (right panels). **F,** UMAP projections of IL1RL1 expression. **G,** Pseudotime projection and trajectory inference on UMAP representation of blood NK cells clusters identified by scRNAseq by using Slingshot. UMAP representation of trajectory analysis on the distinct clusters identified by scRNAseq. **H,** Enrichment analysis of our ST2^+^ NK signature according to pseudotime in blood NK cells analyzed by scRNAseq. NK cell clusters identified in panel F are colored along the pseudotime. **I,** Heatmap showing the top-10 regulons predicted by DoRothEA(34) from scRNAseq data and sorted per NK cell cluster, according to pseudotime. The color scale is based on the z-score of the regulon activity. The z-score distribution ranges from −2 (purple) to 2 (yellow).

### Combination of IL-33 with IL-12 promotes ST2-dependent NK cell antitumor activity *in vivo*

As observed in human NK cells, IL-12 induced ST2 expression on a subset of murine splenic NK cells (Supplementary Fig. S6A), which subsequently produced IFN-γ in response to IL-33 (Supplementary Fig. S6B). To assess the antitumor potential of this pathway *in vivo*, wild-type mice bearing orthotopic E0771 triple negative breast tumors (35) were treated by intratumoral injection of IL-33 and IL-12, alone or in combination (Fig. 4A-C). Combined IL-12+IL-33 treatment markedly reduced primary tumor growth (Fig. 4A) and significantly extended mice survival (Fig. 4B) compared with control, IL-12, or IL-33 monotherapy. These antitumor effects were linked to increased systemic IFN-γ levels (Fig. 4C) and were partially reversed by IFN-γ neutralization in T/B cell-deficient *Rag2*-ko mice, indicating a contribution of IFN-γ to treatment efficacy (Supplementary Fig. S7A-B). Notably, NK cell depletion completely abolished the antitumor effects of IL-12/IL-33 co-treatment in both wild-type (Fig. 4D-F) and *Rag2*-KO mice (Fig. 4G-I), establishing NK cells as the primary effectors of IL-12/IL-33-induced tumor control. We next investigated whether ST2 signaling was required intrinsically in NK cells by generating *Ncr1*^iCre^ *Il1rl1*^flox^ mice, in which *Il1rl1* is selectively deleted in NK cells (Supplementary Fig. S8A). Efficient NK cell-specific *Il1rl1* ablation was confirmed by the failure of NK cells to upregulate ST2 or produce IFN-γ following IL-12+IL-33 stimulation (Supplementary Fig. S8B-C), while ST2 remained expressed on other immune cells in *Ncr1*^iCre^ *Il1rl1*^flox^ mice (Supplementary Fig. S8D). Consistent with a cell-intrinsic requirement for ST2 signaling, the antitumor activity of IL-12 and IL-33 combination was completely lost in *Ncr1*^iCre/+^ *Il1rl1*^fl/fl^ mice compared with *Ncr1*^iCre/+^ *Il1rl1*^+/+^ controls (Fig. 4J-K). Collectively, these results demonstrate that IL-12/IL-33-induced antitumor function is mediated by ST2-expressing NK cells and is partially dependent on IFN-γ.

**Figure 4.**
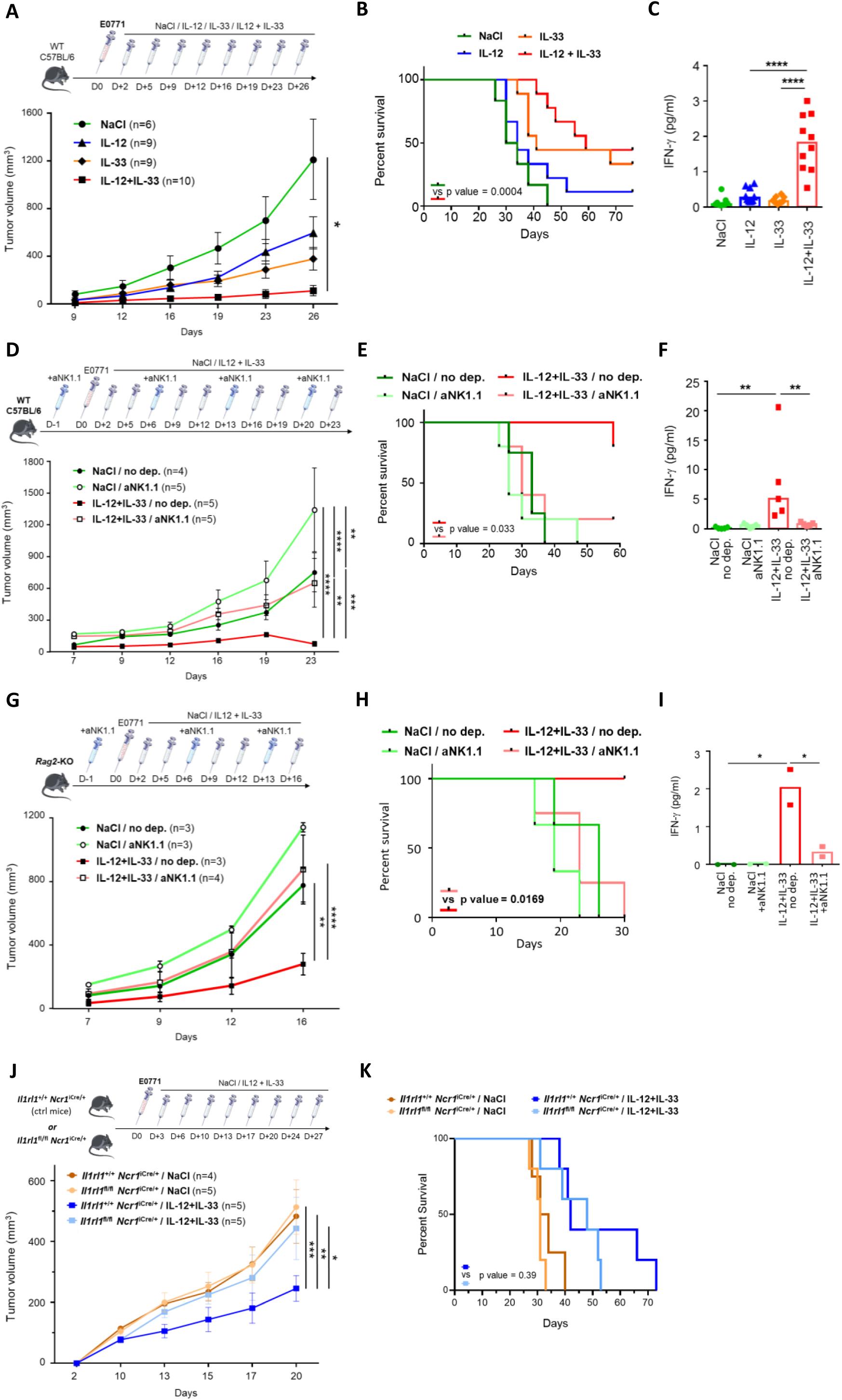
The combination of IL-33 and IL-12 promotes antitumor functions in an NK-cell-dependent manner. **(A-F)** WT, **(G-I)** *Rag2*-KO, and **(J-K)** genetically modified mice were injected orthotopically with 2.5 × 10^5^ E0771 mammary cells on day 0. Cytokines were administered in the tumor area twice weekly. Mice were euthanized when tumors reached 17 mm in diameter. Serum IFN-γ was quantified by ELISA 4 h after treatment on day 5. Data are presented as mean ± SEM (**A**, **D**, **G**, **J**) or median (**C**, **F**, **I**). Two-way ANOVA with Tukey’s multiple-comparison test (**A**, **D**, **G**, **J**), log-rank test (**B**, **E**, **H**, **K**), and Kruskal-Wallis test with Dunn’s multiple-comparison correction (**C**, **F**, **I**) were performed. Group sizes are indicated in the graphs **A**, **D**, **G**, **J**. **A,** Tumor growth in WT mice treated with NaCl, rmIL-12 (10 ng), rmIL-33 (100 ng), or both cytokines twice weekly from day 2 to day 26. **B,** Kaplan-Meier survival curves of mice shown in **A**. **C,** Serum IFN-γ levels measured 4 h after treatment on day 5 in mice shown in **A**. **D,** Tumor growth in WT mice treated with NaCl or rmIL-12 (10 ng) + rmIL-33 (100 ng) twice a week from day 2 to day 23 in the presence or absence of NK cell depletion (anti-NK1.1 antibody). **E,** Kaplan-Meier survival curves of mice shown in **D**. **F,** Serum IFN-γ levels measured 4 h after treatment on day 5 in mice shown in **D**. **G,** Tumor growth in *Rag2-*KO mice treated with NaCl or rmIL-12(10 ng) + rmIL-33 (100 ng) twice a week from day 2 to day 23 in the presence or absence of NK cell depletion (anti-NK1.1 antibody). **H,** Kaplan-Meier survival curves of mice shown in **G**. **I,** Serum IFN-γ levels measured 4 h after treatment on day 5 in mice shown in **G.** **J,** Tumor growth in control (*Il1rl1^+/+^ Ncr1^iCre/+^)* and NK cell-specific ST2-deficient (*Il1rl1^fl/fl^ Ncr1^iCre/+^*) mice treated with NaCl or rmIL-12(10 ng) + rmIL-33 (100 ng) twice a week from day 3 to day 20. **K,** Kaplan-Meier survival curves of mice shown in **J**.

### ST2^+^ NK cells efficiently infiltrate tumors and retain responsiveness to IL-33 and IL-12 stimulation

We next investigated the relevance of ST2^+^ NK cells in tumor settings. Analysis of a human pan-cancer dataset comprising 20,999 tumor-infiltrating NK cells (tiNK) from three independent studies (15,36,37) identified a distinct NK cell cluster highly enriched for ST2^+^ NK cell signature (cluster #6) among eight tiNK clusters (Supplementary Fig. S9A and B). Importantly, the same population was readily detected in a breast cancer sub-atlas containing 2,012 tiNK cells (Fig. 5A). This population exhibited transcriptional programs related to chromatin remodeling, metabolism, differentiation and effector functions shared by both CD56^bright^ (homing, cytokine) and CD56^dim^ (cytokine, differentiation) NK cell subsets (Fig. 5B), closely resembling IL-12-induced blood ST2^+^ NK cells. Consistent with these human data, two NK cell populations (clusters #4 and #7) identified in a murine lymphoma scRNA-seq dataset (38) were also enriched for the ST2^+^ NK cell signature (Supplementary Fig. S9C), indicating that this NK cell state is conserved across species and tumor types. Flow cytometry analysis of breast cancer patient samples identified that ST2^+^ NK cells accounted for ≍5 to 10% of tiNK cells within tumors, while <2% of ST2^+^ NK cells were detected in patients’ blood (Supplementary Fig. S10 for gating strategy and Fig. 5C). As a surrogate for IL-33 responsiveness, we monitored p65 phosphorylation in *ex vivo*-stimulated circulating NK and tiNK cells. While IL-33 had no effect alone, its association with IL-12 induced p65 phosphorylation in a subset of circulating NK and tiNK cells (Fig. 5D), highlighting the potential of NK cells to respond to IL-33 in cancer patients. In agreement, IL-12 and IL-33 co-stimulation enhanced IFN-γ production by circulating NK and tiNK cells from patients, as compared to IL-12 alone (Fig. 5E). Collectively, these results indicate that ST2^+^ NK cells can be detected in tumors and blood from cancer patients, and respond to IL-12 and IL-33 combination by producing IFN-γ, suggesting a potential ST2^+^ NK cell-dependent antitumor function of IL-33.

**Figure 5.**
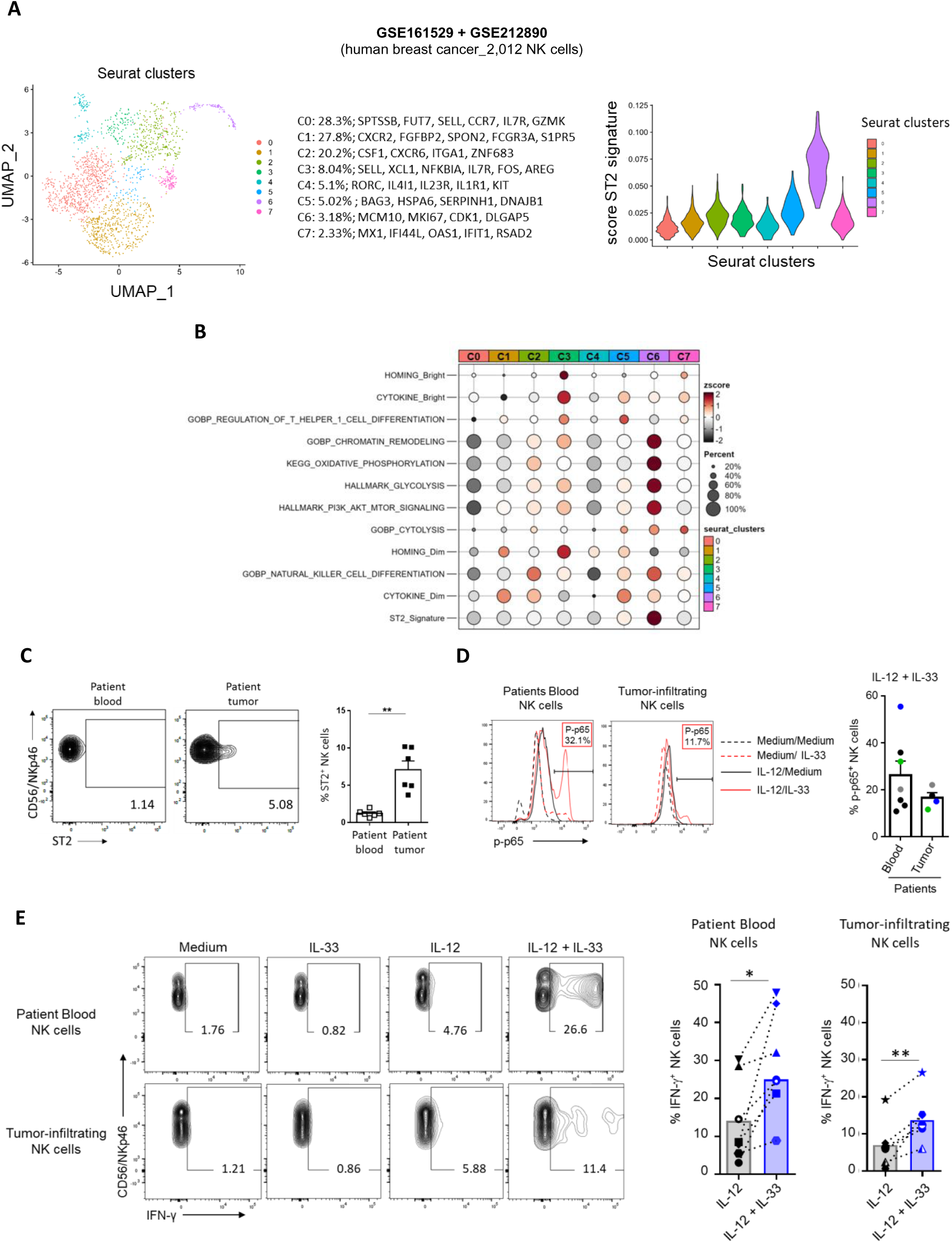
ST2⁺ NK cells infiltrate tumors efficiently and maintain responsiveness to IL-33 and IL-12. **A,** UMAP visualization of tiNK subsets and violin plots showing the distribution of the ST2^+^ NK cell 233-gene signature (see Table 1) across tiNK clusters identified from scRNA-seq analysis of NK cells infiltrating breast cancer (GSE161529 (37) and GSE212890 (15) datasets). **B,** Bubble plot representing the expression of relevant biological pathways shown in Fig. 3D in tiNK cell clusters from breast cancer datasets (GSE161529 (37) and GSE212890 (15)). Dot size encodes the percentage of positive cells and dot color the average expression. **C,** Flow cytometry analysis of ST2 protein expression on the surface of CD56/NKp46^+^ NK cells among PBMCs and tumor cell suspensions from breast cancer patients (n = 6). Representative dot plots (left) and quantification (%) (right) of ST2^+^ NK cells. Results are expressed as mean + SEM. Paired two-tailed Student *t-*test was performed. **D,** PBMCs (n = 7) and tumor cell suspensions (n = 4) from breast cancer patients were activated with medium or IL-12 for 24 h prior to the addition of medium or IL-33. p65 phosphorylation was analyzed 5 min after the addition of IL-33 by flow cytometry in CD56^+^ CD7^+^ NK cells. Representative histogram plots (left) and quantification (%) (right) of p-p65^+^ NK cells after IL-12 and IL-33 combination. Results are expressed as mean + SEM. Unpaired two-tailed Student *t-*test was performed with no statistically significant difference. **E,** PBMCs and tumor cell suspensions from breast cancer patients (n = 6) were activated as indicated (10 ng/mL of each cytokine) for 24 h prior to flow cytometry analysis of intracellular IFN-γ. Representative dot plots (left) and quantification (%) (right) of IFN-γ^+^ NK cells. Symbols represent individual breast cancer patients and histogram bars the median. Wilcoxon matched-pairs signed rank test was performed.

### *An NK*^hi^-*IL33*^hi^ score predicts favorable clinical outcome across cancers

To assess the clinical relevance of the IL-33/ST2 axis, we analyzed The Cancer Genome Atlas (TCGA) pan-cancer datasets and found that both *IL33* and *IL1RL1* were frequently downregulated in tumors relative to adjacent normal tissues, including invasive breast carcinoma (IBC) (Supplementary Fig. S11A), suggesting that loss of IL-33/ST2 signaling may represent a mechanism of immune escape. Consistently, transcriptomic analysis of human breast tissues (39,40) showed a progressive decline in *IL33* expression during tumor progression, while expression largely confined to the stromal compartment (Supplementary Fig. S11B). These observations were supported at the protein level, where IL-33 expression was higher in normal mammary epithelial cells than in malignant cells and was also detected in endothelial cells within the stroma (Supplementary Fig. S11C-D). Although reduced during tumor progression, soluble IL-33 remained detectable in most breast tumors and was present at significantly higher levels in luminal than in HER2⁺ or triple-negative tumors (Fig. 6A). We therefore investigated whether preservation of IL-33/ST2 signaling could influence clinical outcome. Notably, high expression of either *IL33* or *IL1RL1* was frequently associated with improved Progression-Free Survival (PFS) across multiple cancer types (Supplementary Fig. S12A-C). In breast cancer, *IL33*-based patient stratification remained independently prognostic when adjusted for age, molecular subtype, and tumor stage (Supplementary Fig. S12D). Using a validated NK cell gene signature (41) (Supplementary table 8), we observed higher *IL33* expression in *NK*^hi^ BRCA tumors, suggesting a positive association between these two parameters (Fig. 6B). Notably, combined patient stratification identified an *NK*^high^-*IL33*^high^ subgroup with significantly prolonged PFS across six cancer types (Fig. 6C and Supplementary Fig. S13A,B), including breast cancer in which a trend toward improved overall survival (OS) was also observed (Fig. 6D). Importantly, the *NK*^high^-*IL33*^high^ signature remained independently associated with favorable PFS in breast cancer in multivariate analysis adjusted for age, molecular subtype, and disease stage (Fig. 6E). Collectively, these findings identify the IL-33/ST2 axis as a clinically relevant determinant of patient outcome and suggest that IL-33 is particularly beneficial in the context of high NK cell infiltration.

**Figure 6.**
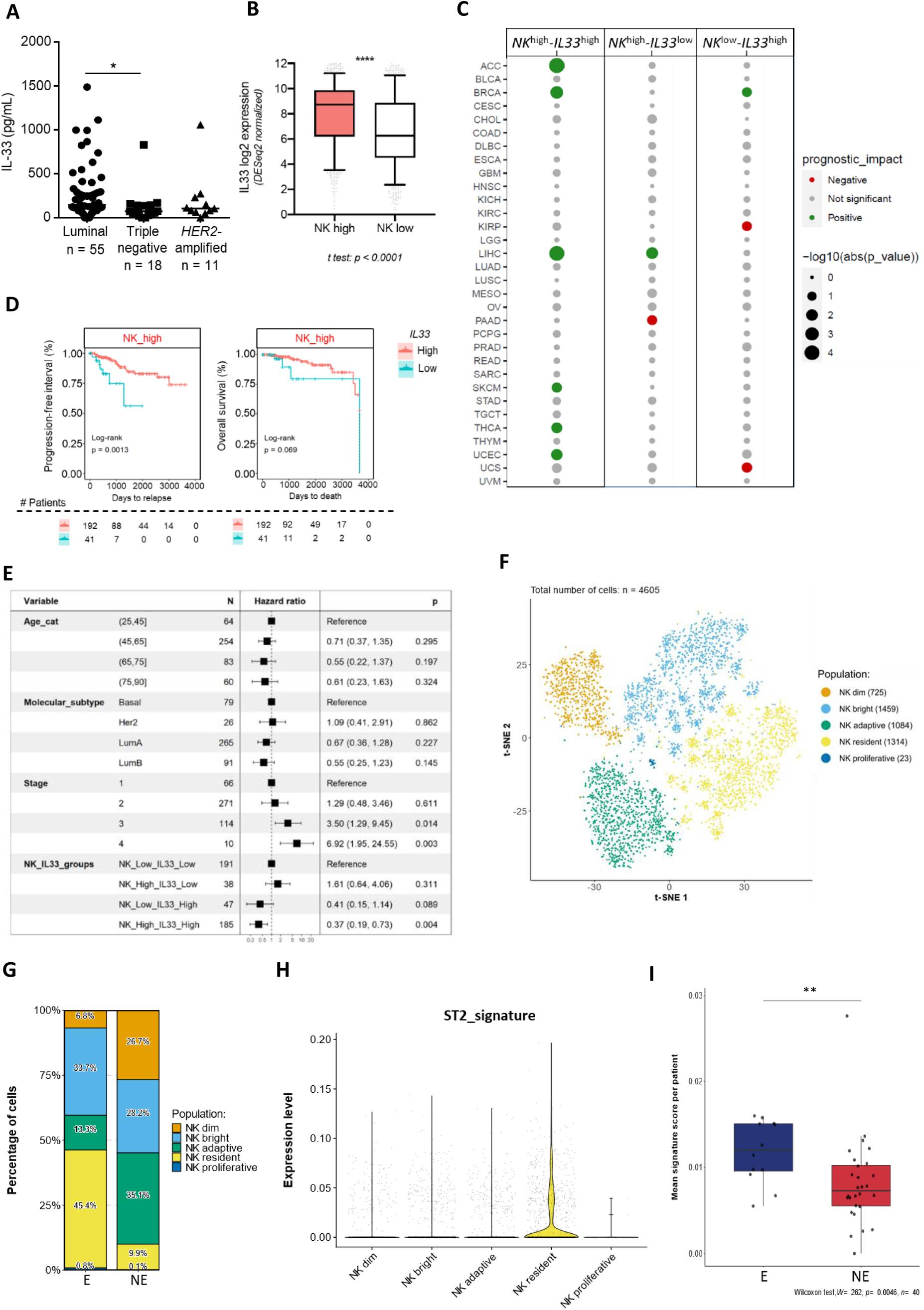
Prognostic value of an *NK*^hi^/*IL33*^hi^ signature and association of ST2^+^ NK cells with response to immune checkpoint blockade. **A,** IL-33 protein was quantified by Luminex assay in breast tumor-derived supernatants (n = 89). Kruskal-Wallis test with Dunn multiple comparisons test was performed. **B,** Boxplots display *IL33* gene expression in NK^hi^ vs NK^low^ breast cancer patients from the TCGA database. An unpaired Student *t*-test was performed. NK signature was defined in (41). **C,** Summary of p-values associated with the log rank test performed to evaluate prognostic value of *NK*^high^/*IL33*^high^, *NK*^high^/*IL33*^low^ and *NK*^low^/*IL33*^high^ scores as compared to *NK*^low^/*IL33*^low^ in 32 human cancers from TCGA database (legend is the same as in panel A, MESO: Mesothelioma; UVM: Uveal Melanoma were added). Results are represented as a bubble map showing positive (green) or negative (red) impact on progression-free survival. Dots size represents p-values obtained by log-rank test. An unpaired Student *t-*test was performed. NK signature was defined in (41). **D,** Kaplan-Maier curves for progression-free survival (left panel) and overall survival (right panel) for *NK*^high^ patients stratified according to *IL33* expression in breast cancer dataset from TCGA. p-values were obtained with log-rank test. NK signature was defined in (41). **E,** Multivariate Cox analysis of the impact of *NK*^high^/*IL33*^high^ score on prognosis in breast cancer patients from TCGA dataset regarding the age, molecular subtype, and stage of the tumors. p-values were obtained with log-rank test. NK signature was defined in (41). **F,** t-SNE plots visualizing the total ti-NK cell clusters from scRNAseq dataset EGAD00001006608 (42). Each dot represents a single cell and is colored according to NK-cell subset identity. **G,** Stacked bar plots showing the relative proportions of tiNK subsets according to ICB response status, inferred from T-cell clonal expansion profiles as Expanded (E) versus Non-Expanded (NE) from scRNAseq dataset EGAD00001006608 (42). **H,** Violin plots showing the distribution of the ST2^+^ NK cell top 10-gene signature (see Table 1) enrichment scores at single cell level, calculated using the Mann–Whitney U statistic, across tiNK subsets from scRNAseq dataset EGAD00001006608 (42). **I,** Pseudobulk analysis of mean top 10-gene ST2+ NK signature scores per patient according to clonal expansion status from scRNAseq dataset EGAD00001006608 (42). Each dot represents the aggregated single-cell expression profile of individual patient scored for the ST2 NK signature. Statistical significance was assessed using a two-sided Wilcoxon rank-sum test (\**p* < 0.05, \*\**p* < 0.01, \*\*\**p* < 0.001).

### ST2^+^ NK cells are associated with improved response to immune checkpoint blockade

Given their potent polyfunctional anti-tumor activity, we investigated whether ST2^+^ NK cells were associated with response to immune checkpoint blockade. Analysis of a public scRNA-seq dataset from anti-PD1-treated breast cancer patients, including both pre- and on-treatment samples (42), identified five tiNK clusters, including conventional CD56^bright^ and CD56^dim^ NK cells, adaptive, resident, and proliferating NK cells (Fig. 6F). Consistent with recent findings (18), CD56^dim^ NK cells were enriched in T cell non-expanded (NE) patients considered as non-responders, whereas resident NK cells were enriched in T cell expanded (E) patients considered as responders (Fig. 6G and Supplementary Fig. S14A). Notably, ST2^+^ NK cells identified using the ST2^+^ NK cell signature derived from our RNAseq analysis, mapped predominantly to the resident NK cell compartment and were significantly enriched in E/responding patients (Fig. 6H-I and Supplementary Fig. S14B). This association is significant in patients treated with first line immunotherapy (cohort A, n = 29) and shows a non-significant trend in patients receiving second line immunotherapy following neoadjuvant chemotherapy (cohort B, n = 11), likely due to the limited sample size (Supplementary Fig. S14C-S14E). Together, these results link IL-33-responsive ST2^+^ NK cells to favorable response to PD-1 blockade in breast cancer.

## DISCUSSION

Previous studies have established a central role for the IL-33/ST2 pathway in type 2 immunity, yet its contribution to type 1 immune responses and antitumor immunity has remained poorly defined. Here, we identify a previously unrecognized NK cell state, termed ST2⁺ NK cells, that integrates IL-12 and IL-33 signals to drive potent antitumor immunity. Positioned between canonical CD56^bright^ and CD56^dim^ NK cells, these metabolically active, highly proliferative, and polyfunctional effectors are present in human tumors and contribute to tumor control in mice through an NK cell-, IFN-γ-, and ST2-dependent, yet T cell-independent mechanism. An *NK*hi-*IL33*hi score predicts favorable survival across cancers, while ST2^+^ NK cell abundance correlates with improved responses to immune checkpoint blockade. Together, these findings establish ST2⁺ NK cells as key mediators of IL-33-driven antitumor responses and identify the IL-33/ST2/NK cell axis as promising target for cancer immunotherapy.

Mechanistically, we demonstrate that IL-12 is a key upstream regulator of ST2 expression in both human and mouse NK cells. IL-12 signaling directly induces ST2 through STAT4 binding to the *Il1rl1* promoter, thereby licensing NK cells to respond to IL-33. Consistent with this mechanism, a recent study showed that the T-bet and STAT4 transcription factors orchestrate the activation of *Il1rl1* promoter in NK cells and other type-1 lymphocytes during viral infection (43). Our findings therefore position IL-12 as a molecular switch that redirects IL-33 activity toward type 1 immunity. Similar to well-established IL-12/IL-18 axis, which promotes IFN-γ production through STAT4/c-Jun complexes and p38-MAPK activation in human NK cells (44,45), IL-33 similarly activates c-jun (46) and p38-MAPK signaling (our data and (31)), providing a mechanistic basis for its synergy with IL-12. However, IL-33 extends beyond IFN-γ induction by promoting broad array inflammatory cytokines and chemokines production by ST2+ CD56^dim^ NK cells, while markedly enhancing their IL-2-driven proliferation, despite the intrinsically limited proliferative capacity of conventional CD56^dim^ NK cells. These observations reveal a distinctive activation program that couple effector function and proliferation, which is of particular interest to fine-tune the harnessing of NK cell effector functions.

Single-cell transcriptomic and functional analyses position ST2^+^ NK cells as an IL-12-induced state within the NK cell differentiation continuum, bridging CD56^bright^ and CD56^dim^ NK cells. This model aligns with previous descriptions of intermediate NK populations defined by distinct phenotypic (GZMK, XCL1, CD94/NKG2A, CD62L, CX3CR1, KIRs, or CD57 (6,8,9,47)), transcriptional, and epigenetic programs (12,14,16,48–50). Functionally, besides broad cytokine-producing capacity, ST2^+^ NK cells combine the proliferative and metabolic fitness of CD56^bright^ NK cells with the effector functions of CD56^dim^ NK cells, endowing them with an unusual degree of polyfunctionality. These features identify ST2^+^ NK cells as a distinct functional state with considerable therapeutic potential for NK-cell-based immunotherapy.

The presence of ST2^+^ NK cells in both human and mouse tumors strongly suggests local IL-12-driven conditioning within the tumor microenvironment. We previously reported significant intratumoral levels of IL-12p40 and IL-12p70 in advanced breast cancer patients (51), likely produced by activated DCs (52). This is consistent with studies demonstrating that DC-derived IL-12 is required for optimal NK cell-mediated tumor control (53), while NK cell-derived IFN-γ sustains IL-12 production of by cDC1 (54), supporting a positive feedback loop that amplifies ST2^+^ NK cell responses. Importantly, dysfunctional NK cells isolated from patients with advanced solid tumors can be reactivated *ex vivo* by IL-12 and IL-33, resulting in robust IFN-γ production. These findings suggest that IL-33 exerts antitumor activity when integrated into an appropriate IL-12-driven inflammatory context, whether released locally following cell stress and death or delivered therapeutically. Consistently, our *in vivo* data demonstrate that NK cell-intrinsic ST2 signaling is required for the anti-tumor activity of the IL-12 and IL-33 combination, promoting ti-NK cell activation and systemic IFN-γ production. While IL-33 alone can promote antitumor immunity and enhance response to immunotherapy (our data and (55–57)), we demonstrate that its activity is substantially amplified by IL-12-driven ST2 expression. This mechanism complements previously described effects of IL-33 on tumor-resident DCs (58), eosinophils (59), and ILC2 (60) that collectively support antitumor T cell immunity. Thus, IL-33 appears to orchestrate a broad antitumor network in which NK cells represent a major effector component.

Although *IL33* expression was associated with favorable prognosis across multiple cancer types (our data and (61,62)), we show that the addition of NK cell infiltration markedly enhanced its predictive value. Patients with *NK*^hi^-*IL33*^hi^ tumors consistently exhibited superior survival, indicating that NK cells are key effectors of IL-33-mediated tumor control. Moreover, ST2^+^ NK cells were enriched in responders to anti-PD-1 therapy and associated with stronger T-cell responses, further supporting their role in promoting effective antitumor immunity. These clinical observations position the IL-33/ST2/NK-cell axis not only as a determinant of patient outcome but also as a potential lever to enhance the efficacy of immune checkpoint blockade. Collectively, these observations supports the emerging concept of IL-33 as a clinically relevant immunotherapeutic agent, capable of enhancing checkpoint blockade efficacy (58,59,63,64), reprogramming the immunosuppressive tumor microenvironment (65) and mobilizing multiple antitumor immune populations (55–57,66), including the ST2⁺ NK-cell program described here. Recent advances, such as CAR T cells engineered to express IL-2 and IL-33, further emphasize the translational potential of exploiting this pathway (67,68).

In conclusion, we identify in humans and mice a previously unrecognized ST2^+^ NK cell state that emerges during NK cell differentiation under IL-12 instruction. These metabolically active, highly proliferative, and polyfunctional effectors integrate IL-12 and IL-33 signals to drive potent antitumor immunity. Our findings establish IL-33/ST2/NK cell axis as a clinically relevant determinant of cancer outcome and response to immune checkpoint blockade, and provide a mechanistic framework for the therapeutic strategies aimed at harnessing ST2⁺ NK cells to enhance antitumor immunity.

## METHODS

### Human breast tumors and blood

Fresh tumors and blood samples (collected in EDTA anticoagulant-containing tubes) from breast cancer patients were provided by the tissue bank of CLB (BB-0033-00050, CRB-CLB, Lyon, France, French agreement number: AC-2019-3426), after approval from the institutional review board and ethics committee (L-06-36) and patient written informed consent, in accordance with the Declaration of Helsinki. Healthy donor (HD) blood samples were obtained from the ‘Etablissement Français du Sang’ (Lyon). Next-generation tissue microarray (ngTMA®) (https://www.ngtma.com) generated from human breast tumors (69) (n = 128 patients) was provided by the Tissue Bank Bern and approved by the Cantonal Ethics Committee of Bern (200/2014 and 2018-01502).

### Human peripheral blood NK cell isolation and culture

Peripheral Blood Mononuclear Cells (PBMCs) were isolated from whole blood of healthy donors by density gradient centrifugation on Lymphocyte Separation Medium (Eurobio). Total NK cells were purified from PBMCs by negative immune-selection using the Human NK cell isolation kit (Miltenyi) following the manufacturer’s instructions and purity always exceeded 95%. In some experiments, NK cell subsets were FACS-sorted as CD56^dim^ CD16^+^ and CD56^bright^ CD16^-^ cells. Cells were cultured in RPMI GlutaMAX/10% FBS/1% penicillin-streptomycin (complete RPMI, cRPMI). Human PB NK cells were cultured in cRPMI supplemented, as indicated, with IL-12 (Miltenyi), IL-15 (Peprotech), IL-2 (Chiron), IFN-α2b (Schering-Plough), IL-1α (Peprotech), IL-1β (Peprotech), IL-18 (MBL) or IL-33 (Miltenyi) at concentrations specified in the figure legends. For receptor stimulation assays, anti-NKp46 and anti-NKp30 agonistic antibodies (R&D Systems) were coated onto NUNC 96-well plates (Thermofisher) at 1 µg/mL overnight at 4°C. For blocking experiments, cells were incubated with anti-ST2 (10 µg/mL, R&D), anti-IL-18 (1 µg/mL, MBL), mIgG1 (1 or 10 µg/mL, R&D) antibodies or IL-1RA (100 ng/mL, Peprotech).

### Ex vivo stimulation of mouse splenic NK cells

Single-cell suspensions were prepared from spleens from C57BL/6J mice (CharlesRiver) after red blood cell lysis with Pharm Lyse^TM^ lysing buffer (BD Biosciences). Splenocytes were activated in cRPMI supplemented with IL-12 (Miltenyi), IL-1α (Miltenyi), IL-1β (Miltenyi), IL-18 (MBL) or IL-33 (Miltenyi) for 24 hr at concentrations specified in the figure legends.

### Preparation of single cell suspensions from human breast tumor tissue

Sections of the resected tumors selected by the pathologists were placed in RPMI supplemented with 100 IU/mL penicillin (Invitrogen) and 100 μg/mL streptomycin (Invitrogen). Tissues were mechanically disrupted, supernatants were collected and frozen for subsequent cytokine and chemokine quantification by Luminex assay following the manufacturer’s instructions. Tumor pieces were then digested for 45 min at 37°C in RPMI supplemented with 100 IU/mL penicillin (Invitrogen) and 100 μg/mL streptomycin (Invitrogen) using 1 mg/mL of collagenase IV (Sigma Aldrich) and 20 μg/mL of DNase I (Sigma Aldrich). Digested samples were then filtered on a 70 μm cell strainer and re-suspended in cRPMI for activation and flow cytometry analysis.

### Tumor cell lines

The mouse E0771 (derived from a spontaneous triple negative mammary tumor that arose in a C57BL/6 mouse) and the human K562 (originally generated from a female leukemia patient) cell lines were cultured in RPMI-1640 (Sigma) supplemented with 10% FCS (GIBCO), 2mML-glutamine, and 100 U/ml penicillin/streptomycin (ThermoFisher).

### Animals and in vivo tumor models

C57BL/6J mice were purchased from Charles River Laboratories. *Ncr1*^iCre/+^ mice (70) were obtained from Pr Eric Vivier and *Il1rl1*^fl/fl^ mice were obtained from Dr Giorgio Trinchieri. Details about the generation of the *Il1rl1*^f/f^ mice have not been published. The information about this line is available upon request to G.T. Mouse lines were interbred in our P-PAC animal facility of the Cancer Research Center of Lyon to obtain the final strains described in the text. Mice were maintained at our animal facility, under specific pathogen-free conditions in accordance with standards for animal care. Experiments were conducted with female mice aged between 8-12 weeks. All animal experiments were approved by the local and the French Ethical Committee for Animal Experimentation (approval numbers #11869 and #41163). Mice were orthotopically injected into the 4^th^ mammary fat pad with 2.5x10^5^ E0771 tumor cells in 200 µL of sterile NaCl at day 0. For NK cell depletion experiments, mice were injected intra-peritoneally with 200 µg/mouse of anti-NK1.1 antibody (BioXcell) or an equivalent volume of NaCl solution one day prior to tumor cell injection and then once a week. To evaluate NK cell depletion efficiency, mice were treated with tetracaine analgesic eye drops and maintained under anesthesia using isoflurane to perform retro-orbital blood collection. Blood NK cell frequency was measured by flow cytometry. For IFN-γ blocking experiments, *Rag2*-KO were injected intra-peritoneally with 250 µg/mouse of anti-IFN-γ antibody (BioXcell) or an isotype control antibody twice a week (see supplementary table 9). For evaluation of therapeutic effect, mice were injected with 10 ng/mouse of rmIL-12 (R&D) or 100 ng/mouse of rmIL-33 (Biolegend) resuspended alone or in combination in 50 µL of NaCl and injected directly in the tumor area twice a week from day 2 until day 26, as a maximum, depending on the experiments. Control mice were injected with an equal volume of NaCl solution. Tumor sizes were monitored with a digital caliper (Mitutoyo) twice a week and expressed as volume of a sphere ((smaller length)^2^ × longer length) from day 9 to the end of experiments. Mice were sacrificed when longer length of the primary tumor reached > 17 mm^2^. All cells were tested for mycoplasma prior to injection using MycoAlert kit (Lonza). For serum collection, blood was collected as described above on day 5, 4 h after cytokine injection. Blood was then incubated at room temperature for 30 min to allow coagulation, centrifuged at 15,000rpm for 15 min and the serum was collected and aliquoted for preservation at -80°C prior IFN-γ quantification.

### Flow cytometry

Cells were stained for the expression of surface markers using antibodies listed in Supplementay table 9 for 30 min at 4°C. Cell viability was determined using Zombie dye (Biolegend) following the manufacturer’s instructions. For intracellular IFN-γ staining, GolgiPlug (BD Bioscience) was added to culture media 4 h prior to the staining. Then, cells stained for the expression of surface markers as previously described were fixed and permeabilized using FoxP3/Transcription Factor Staining Buffer Set (eBioscience) prior to intracellular staining for 30 min at 4°C. Cells were washed twice with PBS prior acquisition on a LSRII Fortessa flow cytometer (BD Biosciences). Results were analyzed using FlowJo software (Tree Star Inc.).

### Phosphorylation flow cytometry analysis

For p-STAT4 analysis, NK cells were activated or not with IL-12 (0.1 ng/mL; 1 ng/mL or 10 ng/mL) for 1 h. For p-NF-κB/p65, p-p38 and p-S6 analysis, NK cells were activated or not with IL-12 (10 ng/mL) for 24 h. When indicated, IL-1 family members were added during 5min for p-NF-κB/p65, 5min for p-p38 and 1 h for p-S6 analysis. Cells were then fixed with Lyse/Fix Buffer (BD Biosciences) for 10 min at 37°C and permeabilized with Perm Buffer III (BD Biosciences) for 30 min on ice following the manufacturer’s instructions. Cells were then stained for 45 min at room temperature, washed, and directly acquired on a Fortessa cytometer (Becton Dickinson) as previously described.

### Killing assay

PB NK cells were activated with IL-1 family members (10 ng/mL) alone or in combination with IL-12 (10 ng/mL) for 24 h and washed 3 times. K562 cells were labeled with calcein (Invitrogen) at 10 µL/mL in cRPMI for 30 min at 37°C, washed twice, and fixed with sulfinpyrazone (4 mM, Sigma-Aldrich). NK cells were incubated with K562 target cells at indicated effector:target ratios for 4 h à 37°C. Calcein release was measured using Clariostar (Labtech) and the percentage of specific lysis was calculated as followed:

% of lysis = ((Fluomeasured-Fluospontaneous release)) / ((Fluomax-Fluospontaneous release)) x 100

### Proliferation assay

For proliferation assay of total NK cells, NK cells were stained with 5 µM Cell Trace Violet (CTV) (Invitrogen) for 20 min at room temperature, washed twice, and activated with IL-1 family members (10 ng/mL) with or without IL-12 (10 ng/mL) for 24 h in RMPI supplemented with 20% human AB serum. After three washes, IL-2 (100 UI/mL, Chiron) was added, culture medium was renewed every 2 days, and CTV signal was analyzed by flow cytometry on day 3 and 6 for CD56^bright^ and CD56^dim^, respectively.

### IFN-γ ELISAs

Human IFN-γ levels were measured in culture supernatants using the human IFN-γ ELISA kit (R&D) following the manufacturer’s instructions. Mouse IFN-γ levels were measured in serum using the Extra Sensitive IFN-gamma Mouse ELISA Kit (ThermoFischer) following the manufacturer’s instructions.

### Luminex immunoassay

Culture supernatants of activated NK cells were harvested to perform IFN-γ, TNF-α, GM-CSF, IL-8, CCL3 (MIP-1α), CCL4 (MIP-1β), CCL5 (RANTES), IL-2, IL-3, IL-4, IL-5, IL-6, IL-10, IL-13, IL-17A, IL-22, and G-CSF quantification using a human Bio-plex PRO assay (Biorad) following the manufacturer’s instructions and acquired with the Luminex200 instrument (Luminex).

### ST2 promoter region analysis

The human *ST2* gene sequence was obtained from Gen-Bank and then submitted to Eukaryotic Promoter Databank (EPD) (71) to identify the promoter region. Putative binding sites for transcription factors in the *ST2*-regulating region were identified using the PROMO database (72).

### p-STAT4 Chromatin Immunoprecipitation (Ch-IP) and qPCR

PB NK cells were stimulated or not with IL-12 as previously described and Ch-IP was performed using ChIP-IT PBMCs Kit (Active Motif) according to the manufacturer’s instructions. Briefly, cells were cross-linked with 1% formaldehyde then sonicated on ice for 6 min (6 cycles of 1 min; 30 s on / 30 s off) at 4°C using a water bath-sonicator (Diagenode). Chromatin fragments were reverse cross-linked overnight at 65°C, and 500 ng of each input DNA was run on a 1% agarose gel to confirm that fragment size ranged between 200 and 1500 bp. p-STAT4 immunoprecipitation was performed overnight at 4°C using anti-p-STAT4 antibody (Cell signaling). p-STAT4 immunoprecipitation was validated by Western blot using pre- and post-immunoprecipitated samples stained with anti-p-STAT4 (Cell Signaling). Isolated DNA fragments were purified and quantitative PCR analysis was performed using 1 μg of DNA as template. Primers for *ST2* promoter region were extracted from (73) and we used *PRF1* (Cell Signaling) and Negative Primer Set 1 and 2 (Active Motif) as positive and negative controls for p-STAT4 binding (see Supplementary table 9 for sequences). The Ct Value of each sample was normalized to input DNA fraction and fold-change for p-STAT4 binding was calculated using ΔΔct method. qPCR analysis was conducted in triplicate for each sample.

### mRNA isolation and quantitative PCR

Total RNA extraction was performed using NucleoSpin RNA Kit (Macherey-Nagel) following the manufacturer’s instructions. Reverse transcription was performed with the iScript Reverse Transcription kit (BioRad) using 350 ng total RNA extract. Real Time quantitative PCR (RT-qPCR) analysis was performed using TaqMan gene expression assay with *IL1RL1* specific primers (Hs00249384_m1, Life Technologies). Normalization of *IL1RL1* expression was performed using *GADD45a* (Hs00169255_m1, Life Technologies) expression.

### RNA-seq of FACS-sorted ST2+ NK cells

MACS-purified total NK cells were cultured in RPMIc in the presence of IL-12 (10ng/mL). After 48h, NK cells were stained with antibodies for CD3, CD56, CD16, CD57 (see Supplementary table 9) as previously described, and five different populations were FACS-sorted using a FACS Aria II (BD) and collected in cRPMI. Cell viability was determined by DAPI staining (1 μg/mL, Invitrogen). 400,000 cells from each population were resuspended in 350 µL TCL lysis buffer (Qiagen) supplemented with 1 % β-mercaptoethanol. Total RNA extraction was performed using the Single Cell RNA Purification Kit (Norgen) following the manufacturer’s instructions, including an additional treatment with rDNAse (Qiagen) to avoid DNA contamination. RNA quality was addressed using 4200 TapeStation (Agilent) automated electrophoresis with RIN always exceeding 8, and total RNA was quantified using Qubit™4 Fluorometer (ThermoFischer). Reverse transcription and DNA amplification was performed with SmartSeqV4 (Takara) using 10 ng of RNA as template. DNA library was prepared with Nextera XT DNA library Prep kit (Illumina), hybridized on NovaSeqS1 Flow cell and sequenced on NovaSeq sequencing platform (Illumina) with a paired-end protocol. Raw sequencing reads were aligned on the human genome (GRCh38) with STAR (v2.7.3a) and the annotation of known genes from gencode v33. Gene expression was quantified using Salmon (v1.1.0) and the annotation of protein coding genes from gencode v33.

### RNA-seq bioinformatics analysis of FACS-sorted ST2+ NK cells

All analyses were performed in R programming language (v3.6.3). DESeq2 (v1.26.0) was used to normalize raw counts, to generate principal component analysis (PCA) plots based on the 500 most variable genes and to identify differentially-expressed genes (DEGs). To identify NK subset signatures, DEGs between one subset and the two other subsets of NK cells were filtered considering a log2 fold change (LFC) > 1 and an adjusted p-value < 0.001, resulting in 234 genes enriched in ST2^+^, 396 genes enriched in CD56^dim^ and 418 genes enriched in CD56^bright^ NK cells. These signatures were scored by ssGSEA using the GSVA package. The Morpheus tool (Broad Institute) was used to generate a heatmap representing the relative expression of these DEGs, using an unsupervised hierarchical clustering of genes and samples based on the one minus Pearson correlation metric. To evaluate the intermediate state of ST2^+^ NK cells compared with CD56^bright^ and CD56^dim^ NK cells, DEGs between one subset and the two other subsets of NK cells were filtered considering a log2 fold change (LFC) > 1 and an adjusted p-value < 0.05. The Morpheus tool (Broad Institute) was used to generate a heatmap representing their relative expression. For this heatmap, genes were classified as follows: upregulated DEGs in CD56^bright^ NK cells (n = 828 genes), downregulated DEGs in CD56^dim^ NK cells (n = 1,348 genes), upregulated DEGs in ST2^+^ NK cells (n = 547 genes), downregulated DEGs in CD56^bright^ NK cells (n = 1,009 genes) and upregulated DEGs in CD56^dim^ NK cells (n = 1,024 genes).

### scRNA-seq of ST2+ NK cells

Blood samples from 2 healthy donors were processed simultaneously to isolate NK cells and culture them in cRPMI in the presence or not of IL-12 (10ng/mL). After 48h, NK cells derived from each healthy donor were washed twice in PBS1X and independently stained and barcoded with different fluorochrome-coupled anti-CD56 antibodies and TotalSeq^TM^ B anti-human Hashtag 1 and 2 antibodies (see Supplementary table 9) diluted in staining buffer, for 15 minutes at 4°C. NK cells were then washed twice and resuspended in a cold DAPI-containing staining buffer. NK cells derived from each healthy donor were pooled together to sort total viable CD56^+^CD3^-^ using FACS ARIA III. The same number of viable NK cells was sorted for each donor (0.5M NK cells to reach a final number of 1M cells) and live cell counts were determined with Luna-FL Dual fluorescence cell counter (Logos Biosystems). Both NK pools were loaded onto a 10X G chip to obtain an expected cell recovery population of 10,000 cells per channel (≈5,000 cells per donor) and run on the Chromium iX system (10X Genomics) according to manufacturer’s instructions. Gene Expression and Cell Multiplexing single-cell RNA-seq libraries were generated with the Chromium Single Cell 3′ v.3.1 kit with feature barcoding technology for Cell Surface Protein (10X Genomics). Gene Expression and hashtags-feature barcoding libraries were sequenced on the NovaSeq 6000 platform (Illumina) to reach approximately 50,000 reads and 5,000 reads per cell respectively. Data were demultiplexed, aligned to human genome, and 10X cellular and hashtags barcodes were assigned for each cell. Genes were counted by unique molecular identifier (UMI) counts, using cellranger multi pipeline (v7.1). The filtered feature barcode matrix was? used for further data analysis.

### scRNA-seq data processing and analysis of ST2+ NK cells

Following sequencing, raw data were converted into fastq files using illumina bcl2fastq. and aligned to the human reference genome GRCh38 by running cellranger-7.1.0 from 10x Genomics. Quality control, normalization and dimension reduction were performed using Seurat version 4.2.0 (74). Some visualizations were generated using the SCP R package (75). Low quality cells with less than 200 expressed genes or with more than 20% of reads aligning to the mitochondria were filtered out. Cells expressing more than 50000 molecules were also removed from the downstream analysis. Cell cycle regression has been performed during data scaling using the S.Score and G2M.Score computed by the CellCycleScoring function from Seurat. By using harmony (v1.2) (76), an integration of both samples was performed to remove heterogeneity between the two donors. The Uniform Manifold Approximation and Projection (UMAP) embeddings and unsupervised clustering were then computed using the Leiden algorithm with the 30 first principal components and 50 k-nearest neighbour at a resolution of 0.3. Using the UCell package (v2.4) (77), signature scores were calculated on differentially expressed genes from our bulk RNAseq experiment. Pathways analyses were conducted using the R package escape (v1.12) (78) with the enrichIt function and the UCell method for different gene sets including the MSigDB Hallmark gene set collection (79,80). The activities of transcription factors were inferred using decoupleR (81) and the DoRothEA resource (34) by selecting only regulons with the confidence levels A, B and C. Pseudotime was computed using slingshot (82) by setting the starting point at the “Bright” cluster according to the literature (6), and ST2 signature scores were ploted along the calculated trajectory for each cell.

### Bioinformatic analysis of public scRNAseq datasets of human tiNK cells

GSE139249 (36), GSE161529 (37), GSE212890 (15) public scRNAseq datasets were extracted from the original publications. Filtering steps and metrics were performed as described above in the scRNAseq section. A total of 20,999 tiNK cells from 9 different tumor types from 72 patient samples were collected. Dataset integration was performed using Harmony package and the RunHarmony function was applied to remove batch effects among datasets and samples. The UMAP was run on the first 20 principal components and the neighbor graph was constructed using the FindNeighbors function, which calculates the nearest neighbors for each cell based on the first 20 principal components. The FindClusters function was applied to identify clusters with a resolution set to 0.3. Differentially expressed gene markers of each cluster were identified using the FindAllMarkers function with the following parameters: logfc.threshold = 0.58; min.pct=0.1; adjPval<0.05.

### Bioinformatic analysis of public scRNAseq dataset of mouse tiNK cells

The GSE123534 public scRNAseq dataset from mouse NK cells isolated from lymphoma tumors was used (38). To score the ST2 signature in mouse dataset, we first mapped the human signature genes to mouse genes using the ‘convert_symbols_by_species’ function in the ‘chevreul’ R package (version 0.6.0), and then calculated the ST2 signature scores with the ssGSEA method implemented in the ‘GSVA’ R package (version 1.38.2).

### Analysis of public scRNAseq dataset of patients treated with immune checkpoint blockade

To validate the association between ST2^+^ NK cells and therapeutic response at the single-cell level, we analyzed publicly available scRNA-seq datasets from (42), obtained as preprocessed count matrices from the European Genome-phenome Archive (EGA) (EGAD00001006608). The dataset included 226,635 cells from 84 samples of 42 breast cancer patients treated with anti-PD1 monotherapy (Cohort A, n=31) or combined with neoadjuvant chemotherapy (Cohort B, n=11), collected pre- and on-treatment.

Single-cell analyses were performed following a standard workflow using the Seurat (v5.3.1) pipeline (74). Cohorts were merged following the Seurat v5 layer-based framework, enabling independent normalization and feature selection per batch, followed by integration based on a consensus gene set (74). Batch effects were corrected using Harmony (v1.2.4) (76) and gene signature scores were computed using UCell (v2.10.1) (77). A summary of parameters used at each downstream analysis step is provided in Supplementary table 10.

Cell-type annotations were validated by assessing the concordance between source-dataset annotations and communities identified by unsupervised clustering following preprocessing and dimensionality reduction. Cluster identities were confirmed using UMAP visualization and systematic evaluation of annotation-cluster correspondence (42). Subsequent analyses focused on 71,623 T/NK cells. After identifying the 2,000 most variable genes using FindVariableFeatures(), data were normalized, scaled with NormalizeData() and ScaleData() and analyzed by PCA using RunPCA() (20 first PCs). Clustering (resolution 0.3) identified 12 subsets based on known T-cell and NK-cell gene signatures. Among them, 3,270 MKI67^+^ cycling T/NK cells were re-clustered to isolate cycling NK cells (Supplementary table 10). A total of 4,624 NK cells were further analyzed using 15 PCs. Reclustering (resolution 0.1) identified NK dim, NK bright, NK adaptive, NK resident, and ILCs subsets based on published gene signatures (13,14,16,83,84). After removing ILCs and integrating cycling NK cells, 4,605 NK cells were reclustered while regressing out cell cycle effects using S.Score and G2M.Score values generated with CellCycleScoring(). Final clustering (resolution 0.15) identified NK_dim (n=725), NK_bright (n=1,459), NK_adaptive (n=1,084), NK_resident (n=1,314), and NK_proliferative (n=23) subsets. Final visualization used t-SNE with RunTSNE() which better preserves local structure for NK subset analysis. Analyses of public scRNA-seq datasets were performed in RStudio (v4.4.2) using the rstatix package (v0.7.3). For signature scoring, UCell enrichment scores were computed based on the Mann–Whitney U statistic.

### Pseudobulk and patient-level proportion analyses

Single-cell signature scores were averaged at the patient level to generate mean expression scores for each gene signature, including the *ST2 NK* signature. In parallel, proportions of NK-cell subsets were calculated for each patient relative to total NK cells. Patients from both cohorts (42) were stratified into Expanded (E) and Non-Expanded (NE) groups based on metadata annotations provided in the original study, reflecting clonal expansion status inferred from paired TCR-seq analyses comparing pre-treatment and on-treatment samples.

NK-cell subset proportions were subsequently compared between E and NE patients to evaluate differential enrichment between conditions. Similarly, mean ST2 signature scores were compared between groups to assess whether total NK cells from Expanded patients exhibited enhanced functional enrichment. Comparisons of patient-level NK-cell subset proportions and pseudobulk signature scores between Expanded and Non-Expanded groups were performed using two-sided unpaired Wilcoxon rank-sum tests. Statistical significance was defined as adjusted p<0.05 using the Benjamini–Hochberg correction for multiple testing when applicable.

### Analysis of public cancer patient RNA-seq data

#### TCGA dataset

Upper-quartile normalized expression (UQN) and clinical outcome datasets from The Cancer Genome Atlas (TCGA) were downloaded from the Pan-Cancer Atlas (https://www.cancer.gov/about-nci/organization/ccg/research/structural-genomics/tcga) for 33 tumor types: adrenocortical carcinoma (ACC), bladder urothelial carcinoma (BLCA), breast invasive carcinoma (BRCA), cervical carcinoma (CESC), cholangiosarcoma (CHOL), colorectal adenocarcinoma (COAD), diffuse large B-cell lymphoma (DLBC), esophageal carcinoma (ESCA), glioblastoma multiforme (GBM), head and neck squamous cell carcinoma (HNSC), kidney chromophobe carcinoma (KICH), kidney clear renal cell carcinoma (KIRC), kidney papillary cell carcinoma (KIRP), lower grade glioma (LGG), liver hepatocellular carcinoma (LIHC), lung adenocarcinoma (LUAD), lung squamous cell carcinoma (LUSC), mesothelioma (MESO), ovarian serous cystadenocarcinoma (OV), pancreatic adenocarcinoma (PAAD), paraganglioma & pheochromocytoma (PCPG), prostate adenocarcinoma (PRAD), rectum adenocarcinoma (READ), sarcoma (SARC), skin cutaneous metastatic melanoma (SKCM), stomach adenocarcinoma (STAD), testicular germ cell cancer (TGCT), thyroid carcinoma (THCA), thymoma (THYM), uterine corpus endometrial carcinoma (UCEC), uterine carcinosarcoma (UCS) and uveal melanoma (UVM) to compare *IL33* and *IL1RL1* gene expression in tumor *versus* peritumoral healthy tissue. Progression free survival analyses according to *IL33* and *IL1RL1* gene expression, *NK_signature* expression and combined *IL33* and *NK_signature* expression and plots were performed with R software v 3.6.0, using the packages survival and survminer. For *IL33* and *IL1RL1* genes, patients were stratified into quartiles according to normalized gene expression levels in log2 (RSEM+1) to compare top (*i.e.* high) and bottom (*i.e.* low) quartiles. For *NK_signature*, patients were stratified according to the median of normalized gene expression levels in log2 (RSEM+1) to compare top (*i.e.* high) and bottom (*i.e.* low) halves. The log-rank test was used to determine statistical significance for overall survival between groups of patients. For progression-free and overall survival analyses in the breast cancer dataset, *NK_high* and *NK_low* patients were stratified according to *IL33* expression. The log-rank test was used to determine statistical significance between groups of patients. For multivariate analysis according to *IL33* or combined *IL33 and NK_signature* expression in the breast cancer dataset, a cox model including the main prognostic factors (age, molecular subtype and stage) was used. Gender was removed from the model due to the low number of males in the study.

#### Normal, DCIS, and IBC datasets

Publicly available gene expression datasets were retrieved from the Gene Expression Omnibus (GEO) database. The dataset GSE41228 (39) was used to evaluate *IL33* expression in laser-capture microdissected epithelial and stromal compartments from ductal carcinoma in situ (DCIS) and invasive breast carcinoma (IBC) lesions. The dataset GSE21422 (40) was used to compare *IL33* expression among healthy mammary tissue, DCIS, and IBC samples. Normalized expression values for IL33 were extracted from the corresponding datasets and analyzed according to tissue type and histological compartment. Comparisons of *IL33* expression among multiple groups were performed using the non-parametric Kruskal-Wallis test, followed by the Steel-Dwass-Fligner post hoc multiple comparison test for pairwise analyses. Statistical significance was defined as p < 0.05.

### Immunohistochemical assessment of IL-33 expression

IL-33 staining was performed automatically using a BOND RX autostainer (Leica Biosystems) on FFPE human breast tumor ngTMA (69). Sections were first deparaffinized and antigen was retrieved using Tris buffer (pH 9.0) for 30 min at 95°C. Sections were then stained with goat anti-human IL-33 (R&D Systems, # AF3625; dilution of 1:400 for 30 min) primary antibody. A rabbit anti-goat antibody (Agilent, # E0466) was used as secondary antibody, at a dilution of 1:400 for 15 min. Specific binding of primary antibodies was visualized using a polymer-based visualization system with horseradish peroxidase as the enzyme and 3,3-diaminobenzidine (DAB) as a brown chromogen for IL-33. Sections were counterstained with hematoxylin, dehydrated and mounted with Tissue-Tek Glas Mounting Medium (Sakura). Slides were scanned in high resolution on whole slide scanners Panoramic 250 Flash (3DHISTECH) or NanoZoomer S360 (Hamamatsu). Nuclear IL-33 expression was evaluated in vascular cells, tumor cells, stromal cells, and lymphocytes. Immunostained slides were independently examined by trained observers under light microscopy. IL-33 expression was quantified using the Allred scoring system, which combines staining intensity and the proportion of positively stained cells. Briefly, the proportion score ranged from 0 to 5 according to the percentage of positive cells, and the intensity score ranged from 0 to 3 (absent, weak, moderate, or strong staining). The final Allred score was obtained by summing the proportion and intensity scores, yielding a total score ranging from 0 to 8. For each sample, Allred scores were determined separately for tumor cells, stromal cells, lymphocytes, and vascular cells.

### Data analysis and statistics

Statistical analyses were performed using the GraphPad software and tests conducted are indicated in the Figure legends. P-values lower than 0.05 were considered to be significant, with stars corresponding to * p < 0.05; ** p < 0.01; *** p < 0.001 and **** p < 0.0001. If no stars are indicated, no statistically significant difference was found.

## Data Availability

The bulk RNA-seq data generated during this study are deposited in GEO repository with accession number GEO: GSE199134. The scRNA-seq data generated during this study are deposited in GEO repository with accession number GEO: GSE293848. The raw and processed data are publicly available as of the date of publication. This paper does not report original code. The RNA-seq data from laser-microdissected stromal versus epithelial zones from DCIS and IBC lesions (39) and from healthy mammary tissue versus DCIS and IBC (40) were downloaded from GEO with accession number GEO: GSE41228 and GSE21422 respectively. The scRNA-seq data derived from NK cells infiltrating human tumors (GSE139249 (36), GSE161529 (37), GSE212890 (15)) and mouse tumors (GSE123534 (38)) were downloaded from GEO. The scRNA-seq dataset from anti-PD1-treated breast cancer patients, including both pre- and on-treatment samples, was downloaded from EGA (EGAD00001006608) (42). The Cancer Genome Atlas data were downloaded from Firehose (https://gdac.broadinstitute.org/). Genes Signatures used in this study are listed in supplementary tables 1-8.

## List of Supplementary material

### Supplementary Figures

**Supplementary Fig. 1.** IL-33 in combination with IL-12 strongly activates NK cells through ST2 receptor

**Supplementary Fig. 2.** IL-12/STAT4 signaling induces ST2 expression in NK cells

**Supplementary Fig. 3.** IL-33 preferentially activates a subset of CD56^dim^ NK cells

**Supplementary Fig. 4.** ST2^+^ NK cells display a unique gene signature compared to CD56^bright^ and CD56^dim^ NK cells

**Supplementary Fig. 5.** scRNAseq analysis of IL-12-cultured blood NK cells reveals a distinct subset enriched for the ST2^+^ NK gene signature, characterized by unique transcriptomic features.

**Supplementary Fig. 6.** IL-12 sensitizes mouse splenic NK cells to the production of IFN-γ in response to IL-33

**Supplementary Fig. 7.** Antitumoral effect of IL-33 and IL-12 combination is partially dependent on IFN-γ

**Supplementary Fig. 8.** Validation of *Il1rl1* deletion efficiency in NK cells from the conditional targeting mouse model

**Supplementary Fig. 9.** Unsupervised clustering of public sc-RNAseq datasets of tiNK cells

**Supplementary Fig. 10.** Gating strategy to identify NK cells in human tumor cell suspensions by flow cytometry

**Supplementary Fig. 11.** IL-33 expression decreases during tumor progression and is restricted to vascular cells

**Supplementary Fig. 12.** *IL33* expression correlates with favorable progression-free survival across multiple solid cancers

**Supplementary Fig. 13.** Pan-cancer survival analysis of the NK cell–IL-33 association in TCGA datasets

**Supplementary Fig. 14.** ST2^+^ NK gene signature contribution across tiNK cell subsets stratified by treatment cohort and ICB response status

### Supplementary Tables

**Supplementary table 1.** List of ST2^+^ NK cell signature genes used in this study, Related to Fig. 3B, 3E, 3H, 3I, 5A, 5B, 6H, 6I, Supplementary Fig. S4E, S9A, S9C, S14A-C

**Supplementary table 2.** List of ST2^-^ CD56^bright^ NK cell signature genes used in this study, Related to Fig. 3B, 3E, 3H, 3I, 5A, 5B, Supplementary Fig. S4E, S9A

**Supplementary table 3.** List of ST2^-^ CD56^dim^ NK cell signature genes used in this study, Related to Fig. 3B, 3E, 3H, 3I, 5A, 5B, Supplementary Fig. S4E, S9A

**Supplementary table 4.** List of CYTOKINE_Bright signature genes used in this study, Related to Fig. 3D, 5B

**Supplementary table 5.** List of CYTOKINE_Dim signature genes used in this study, Related to Fig. 3D, 5B

**Supplementary table 6.** List of HOMING_Bright signature genes used in this study, Related to Fig. 3D, 5B

**Supplementary table 7.** List of HOMING_Dim signature genes used in this study, Related to Fig. 3D, 5B

**Supplementary table 8.** List of NK cell signature genes used in this study (from (41)), Related to Fig. 6B-E, Supplementary Fig. S13A, S13A

**Supplementary Table 9.** Research Resource Identifier

**Supplementary Table 10.** Summary of data preprocessing and clustering settings for downstream analyses of the validation scRNA-seq dataset (42)

## Author’s Contributions

**A Eberhardt:** Conceptualization, data curation, formal analysis, supervision, validation, investigation, visualization, methodology, writing–original draft. **E Blanc:** Conceptualization, data curation, formal analysis, supervision, validation, investigation, visualization, methodology, writing–original draft. **V Picant:** Data curation, formal analysis, validation, investigation, visualization, methodology. **A Voissière:** Data curation, formal analysis, visualization, methodology. **L Revol-Bauz:** Data curation, formal analysis, investigation, visualization, methodology. **T Casini:** Data curation, formal analysis, visualization, methodology. **V Alcazer:** Data curation, formal analysis, visualization, methodology. **D Poujol:** Data curation, investigation. **L Tonon:** Formal analysis, visualization, methodology. **R Schneider:** Formal analysis, visualization, methodology. **E Picard:** Data curation, formal analysis, investigation, visualization, methodology. **Y Rocca:** Data curation, formal analysis, investigation, visualization, methodology. **M Ardin:** Formal analysis, visualization, methodology. **F Onodi:** Data curation, formal analysis, investigation, visualization. **C Degletagne:** Data curation, investigation. **C Rodriguez:** Data curation, investigation. **L Moudombi:** Methodology. **B Estavoyer:** Data curation, formal analysis, investigation, visualization, methodology. **E Charrier:** Methodology. **X Wang:** Resources, Data curation, formal analysis, investigation, visualization. **A Stojanovic:** Resources, Data curation, formal analysis, investigation, visualization. **T Rau:** Resources, methodology, investigation. **O Tredan:** Resources, data curation. **I Treilleux:** Resources, data curation. **G Trinchieri:** Resources, methodology. **E Vivier:** Resources, methodology. **J Valladeau-Guilemond:** Supervision, methodology. **A Marçais:** Supervision, funding acquisition, methodology. **T Walzer:** Supervision, funding acquisition, methodology. **P Krebs:** Resources, methodology. **A Cerwenka:** Resources, methodology. **M Hubert:** Data curation, formal analysis, visualization, methodology. **C Caux:** Conceptualization, supervision, funding acquisition, methodology, writing–original draft. **M-C Michallet:** Supervision, methodology. **N Bendriss-Vermare:** Conceptualization, supervision, funding acquisition, methodology, writing–original draft.

## Acknowledgments

Bulk and single-cell RNA sequencings were performed by the Cancer genomics platform (CGP) of the CRCL supported by the SiRIC-LYriCAN program (grant INCa-DGOS-Inserm_12563). Cell sorting was performed at the Flow cytometry core facility of the CRCL. Mouse experiments were performed at the Small animal platform/animal core facility/imaging (P-PAC) platform of the CRCL. Fresh tumor samples and blood samples from breast cancer patients were provided by Biological sample management platform (PGEB) of the CLB (BB-0033-00050, CRB - CLB, Lyon, France; French agreement number: AC-2019-3426). The breast cancer immunostaining was generated by Coya Tapia, with amended data monitoring by Rupert Langer, and stained by the Translational Research Unit in Bern. Stefan Reinhard helped for digitalization of the histology pictures.

We thank S. Boyault, J. Auclair, and C. Audoynaud for performing RNA sequencing. We thank M. Pratviel, B. Vernière, M. Sanchez, and David Neves for excellent technical assistance for mouse experiments. P. Battiston-Montagne and A. Jambon provided help during cell sorting. We want to additionally thank V. Sisirak, Y. Grinberg-Bleyer, and B. Dubois for careful reading of the manuscript, helpful comments, and suggestions.

