## Supplemental data for "A Unique Polyfunctional IL-33-responsive ST2⁺ NK cell State Potentiates Antitumor Immunity and Response to Immunotherapy"

### Supplementary Material

#### Supplementary Figures

**Supplementary Fig. 1.** IL-33 in combination with IL-12 strongly activates NK cells through ST2 receptor

**Supplementary Fig. 2.** IL-12/STAT4 signaling induces ST2 expression in NK cells

**Supplementary Fig. 3.** IL-33 preferentially activates a subset of CD56$^{dim}$ NK cells

**Supplementary Fig. 4.** ST2$^{+}$ NK cells display a unique gene signature compared to CD56$^{bright}$ and CD56$^{dim}$ NK cells

**Supplementary Fig. 5.** scRNAseq analysis of IL-12-cultured blood NK cells reveals a distinct subset enriched for the ST2$^{+}$ NK gene signature, characterized by unique transcriptomic features.

**Supplementary Fig. 6.** IL-12 sensitizes mouse splenic NK cells to the production of IFN-γ in response to IL-33

**Supplementary Fig. 7.** Antitumoral effect of IL-33 and IL-12 combination is partially dependent on IFN-γ

**Supplementary Fig. 8.** Validation of *Il1rl1* deletion efficiency in NK cells from the conditional targeting mouse model

**Supplementary Fig. 9.** Unsupervised clustering of public sc-RNAseq datasets of tiNK cells

**Supplementary Fig. 10.** Gating strategy to identify NK cells in human tumor cell suspensions by flow cytometry

**Supplementary Fig. 11.** IL-33 expression decreases during tumor progression and is restricted to vascular cells

**Supplementary Fig. 12.** *IL33* expression correlates with favorable progression-free survival across multiple solid cancers

**Supplementary Fig. 13.** Pan-cancer survival analysis of the NK cell–IL-33 association in TCGA datasets

**Supplementary Fig. 14.** ST2$^{+}$ NK gene signature contribution across tiNK cell subsets stratified by treatment cohort and ICB response status

#### Supplementary Tables

**Supplementary table 1.** List of ST2$^+$ NK cell signature genes used in this study, Related to Fig. 3B, 3E, 3H, 3I, 5A, 5B, 6H, 6I, Supplementary Fig. S4E, S9A, S9C, S14A-C

**Supplementary table 2.** List of ST2$^-$ CD56$^{bright}$ NK cell signature genes used in this study, Related to Fig. 3B, 3E, 3H, 3I, 5A, 5B, Supplementary Fig. S4E, S9A

**Supplementary table 3.** List of ST2$^-$ CD56$^{dim}$ NK cell signature genes used in this study, Related to Fig. 3B, 3E, 3H, 3I, 5A, 5B, Supplementary Fig. S4E, S9A

**Supplementary table 4.** List of CYTOKINE_Bright signature genes used in this study, Related to Fig. 3D, 5B

**Supplementary table 5.** List of CYTOKINE_Dim signature genes used in this study, Related to Fig. 3D, 5B

**Supplementary table 6.** List of HOMING_Bright signature genes used in this study, Related to Fig. 3D, 5B

**Supplementary table 7.** List of HOMING_Dim signature genes used in this study, Related to Fig. 3D, 5B

**Supplementary table 8.** List of NK cell signature genes used in this study (from (41)), Related to Fig. 6B-E, Supplementary Fig. S13A, S13A

**Supplementary Table 9.** Research Resource Identifier

**Supplementary Table 10.** Summary of data preprocessing and clustering settings for downstream analyses of the validation scRNA-seq dataset (42)

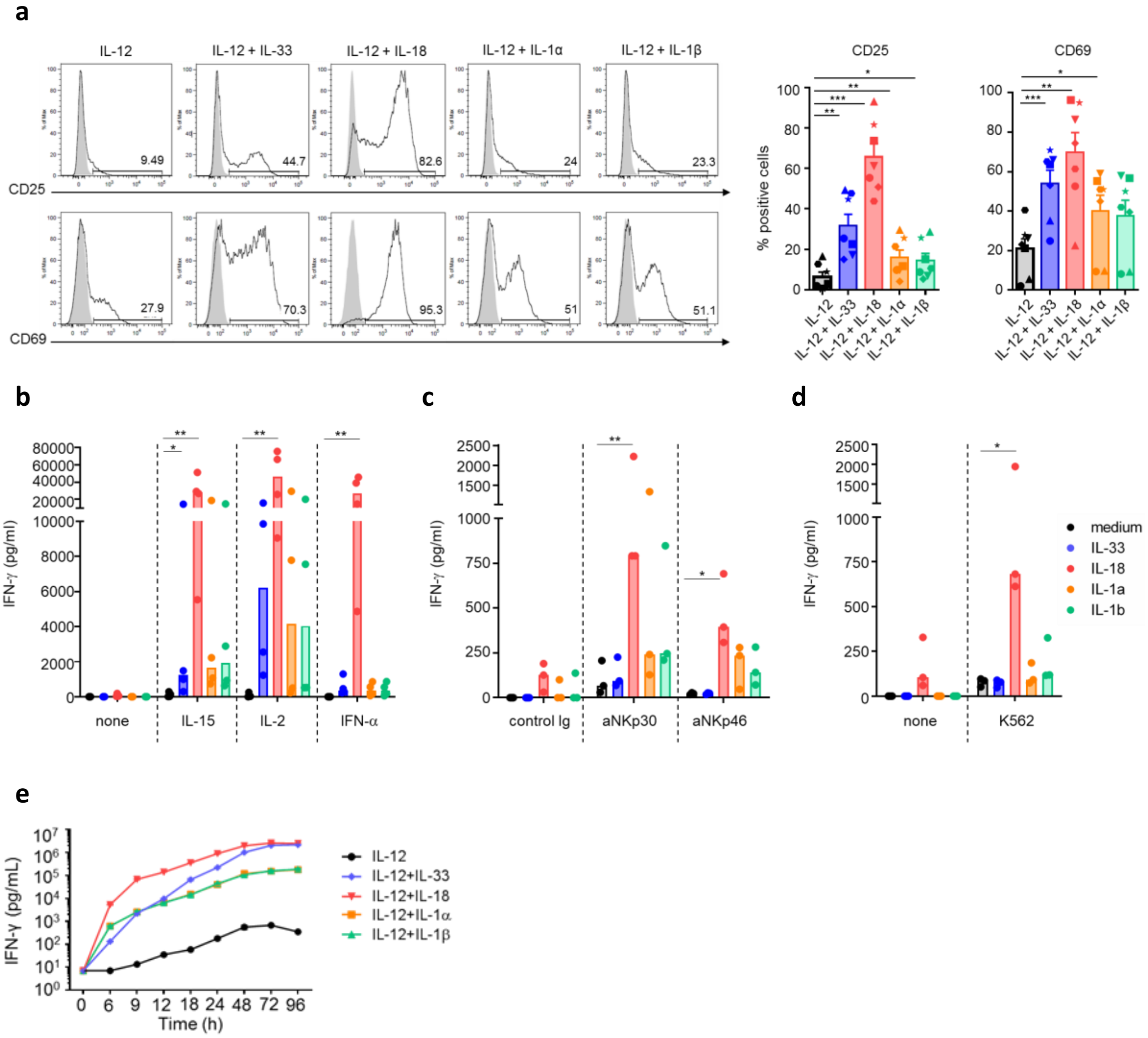


**Supplementary Fig. 1. IL-33 in combination with IL-12 strongly activates NK cells through ST2 receptor**

**a,** NK cells were activated for 24 h with IL-12 (10 ng/mL) alone or in combination with IL-33, IL-18, IL-1α or IL-1β (10 ng/mL) and then stained for CD25 and CD69 (black line) or corresponding isotypic control (grey). Representative histogram plots (left) and quantification (%) (right) of $CD25^+$ or $CD69^+$ NK cells after cytokine stimulation. Symbols represent paired individual experiments (n=7). Results are expressed as mean + SEM. One-way repeated measures ANOVA with Dunnett's multiple comparisons test against levels in IL-12 was performed.

**b-d**) Quantification of IFN-γ secretion by healthy donors' blood NK cells upon stimulation with IL-33, IL-18, IL-1α or IL-1β alone (10 ng/mL) or in combination with (**b**) IL-15 (10 ng/mL), IL-2 (500 UI/mL),

IFN-α (500 UI/mL), (**c**) anti-NKp30 (1 μg/mL), anti-NKp46 (1 μg/mL) agonistic antibodies or (**d**) K562 target cells for 24 h. Histogram bars represent the median (n = 4 individual experiments). Friedman test with Dunn's multiple comparisons test was performed.

**e)** Healthy donors' blood NK cells were activated as described above and supernatants were collected at several time points prior to IFN-γ quantification by ELISA (n = 1).

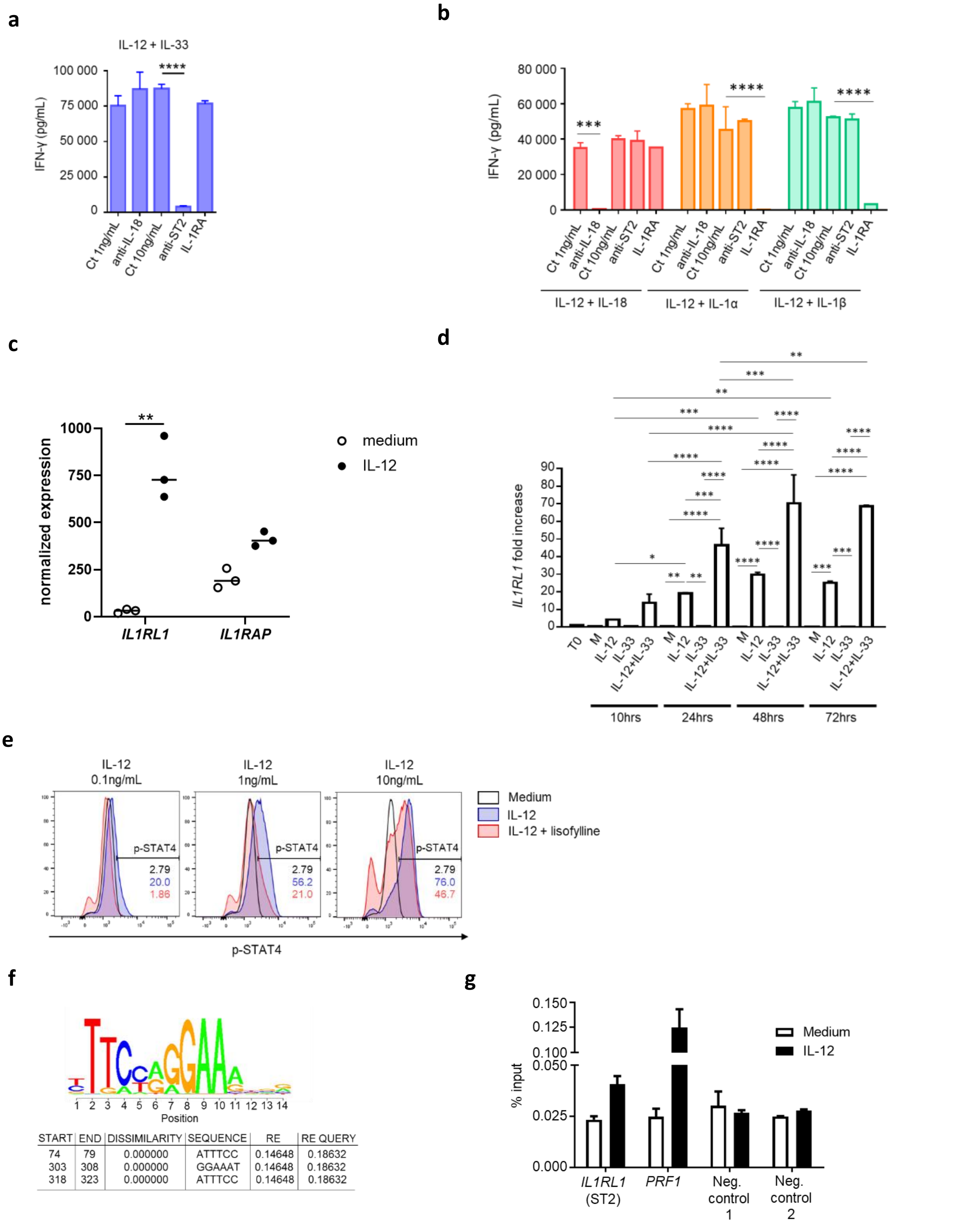


**Supplementary Fig. 2. IL-12/STAT4 signaling induces ST2 expression in NK cells**

**a)** IFN-γ secretion by healthy donors' blood NK cells upon stimulation for 24 h with IL-33 and IL-12 (10 ng/mL each) in the presence of anti-ST2 (10 µg/mL), anti-IL-18 (1 µg/mL) blocking antibodies, IL-1RA antagonist (100 ng/mL) or mIgG1 control antibody (1 or 10 µg/mL as control for anti-IL-18 or anti-ST2, respectively). Results are expressed as mean + SD and are representative of three individual experiments. One-way repeated measures ANOVA with Tukey's multiple comparisons test was performed.

**b)** Healthy donors' blood NK cells were activated by different cytokine combinations in the presence of anti-ST2 (10 µg/mL), anti-IL-18 (1 µg/mL), control mIgG1 antibodies (1 or 10 µg/mL as control for anti-IL-18 or anti-ST2, respectively) or IL-1RA antagonist (100 ng/mL). Supernatants were collected after 24 h and IFN-γ secretion was quantified by ELISA. Results are expressed as mean + SD and are representative of three individual experiments. Two-way ANOVA with Tukey's multiple comparisons test was performed.

**c)** Normalized expression of *IL1RL1* and *IL1RAP* in human NK cells cultured in medium or IL-12 for 24h from our RNAseq datas. Two-way ANOVA with Bonferroni multiple comparisons test was performed; n = 3 experiments.

**d)** *IL1RL1/ST2* relative mRNA expression was analyzed by RT-qPCR at indicated times. Results are expressed as mean + SD and are representative of two individual experiments. Two-way ANOVA with Tukey's multiple comparisons test was performed.

**e)** NK cells were treated or not with lisofylline (500 µM) for 24 h prior to activation with IL-12 (0.1 to 10 ng/mL) for 5 min and p-STAT4 intracellular levels were analyzed by flow cytometry (n = 3 individual experiments).

**f)** *In silico* analysis of putative STAT-4 binding sites in *ST2* promoter region. Dissimilarity indicates the deviation between the identified sequence and the STAT-4 consensus binding site. RE and RE query values indicate the probability to randomly finding the motif respectively considering a model with equiprobability of the four nucleotides and a model with the same nucleotide frequency as the query sequence. A value of 0.001 means that the hit is only expected to occur by chance once in 1 Mb of sequence.

**g)** p-STAT4 was immunoprecipitated in NK cells following activation or not with IL-12 (10 ng/mL) for 24 h and p-STAT4 enrichment in *ST2* promoter region was measured by RT-qPCR analysis. *PERFORIN-1*

*(PRF1)* and negative control sets 1 and 2 were used respectively as positive and negative controls for p-STAT4 binding. Results are expressed as mean + SEM of two individual experiments.

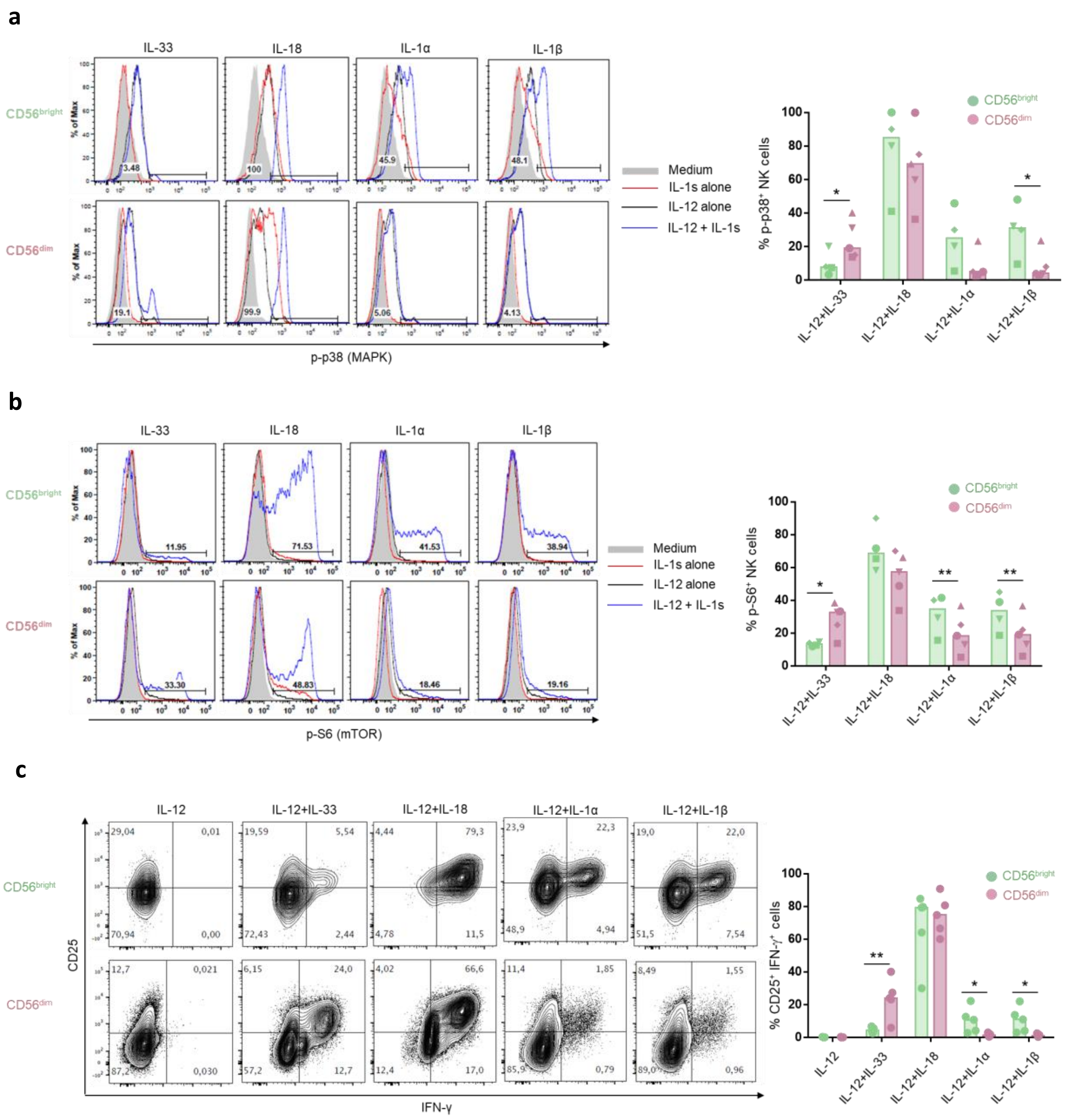


**Supplementary Fig. 3. IL-33 preferentially activates a subset of CD56$^{dim}$ NK cells**

**a-b)** FACS-sorted CD56$^{bright}$ and CD56$^{dim}$ healthy donors' blood NK cells were activated with medium or IL-12 alone for 24 h prior to the addition of IL-33, IL-18, IL-1α or IL-1β, supplemented or not with IL-12. Each cytokine was used at 10ng/mL. **a)** p38 (MAPK) and **b)** S6 (mTOR) phosphorylation was analyzed 5 min and 1 h after the addition of IL-1 family cytokines, respectively. Representative histogram plots (left) and quantification (%) (right) of p-p65$^+$ or p-p38$^+$ NK cells after cytokine activation. Histogram bars indicate the median. Two-way ANOVA with Bonferroni's multiple comparisons test was performed; n = 4 to 5 experiments.

**c)** Healthy donors' blood NK cells were activated as indicated for 24 h, each cytokine was used at 10 ng/mL. CD25 surface and IFN-γ intracellular expression was analyzed by flow cytometry. CD56 surface expression levels were used to discriminate $CD56^{bright}$ and $CD56^{dim}$ NK cells. Representative dot plots (left) and quantification (%) (right) of $CD25^{+}$ IFN-$\gamma^{+}$ NK cells after cytokine activation. Histogram bars indicate the median. Two-way ANOVA test with Bonferroni's multiple comparisons test was performed; n = 5 experiments.

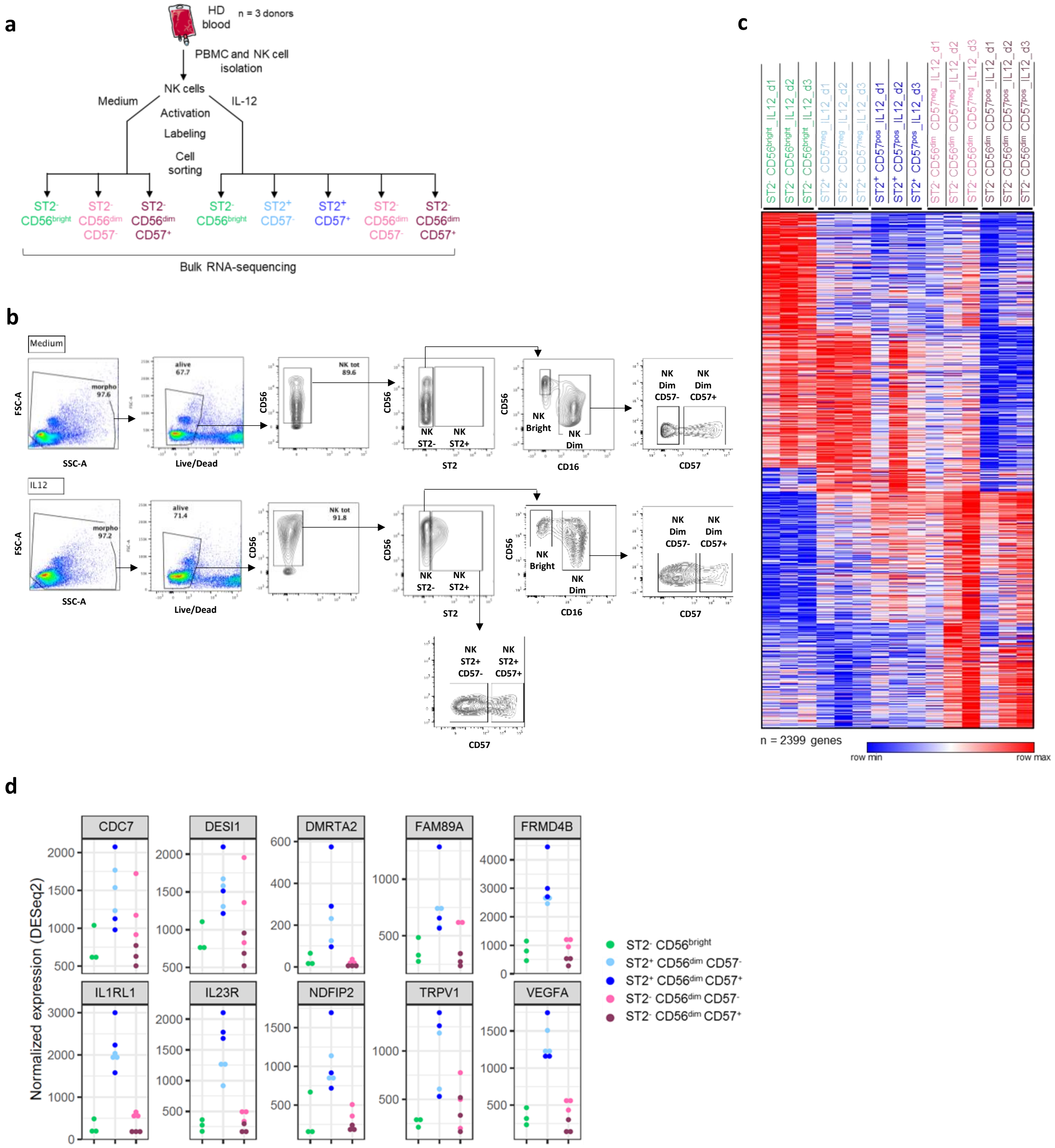
a
HD blood
n = 3 donors
PBMC and NK cell isolation
NK cells
Medium
IL-12
Activation
Labeling
Cell sorting
ST2- CD56bright
ST2- CD56dim CD57-
ST2- CD56dim CD57+
ST2- CD56bright
ST2+ CD57-
ST2+ CD57+
ST2- CD56dim CD57-
ST2- CD56dim CD57+
Bulk RNA-sequencing
b
Medium
IL12
FSC-A
SSC-A
morpho
alive
Live/Dead
CD56
NK tot
NK ST2-
NK ST2+
ST2
NK Bright
NK Dim
CD16
NK Dim CD57-
NK Dim CD57+
CD57
NK ST2+ CD57-
NK ST2+ CD57+
c
ST2- CD56bright_IL12_d1
ST2- CD56bright_IL12_d2
ST2- CD56bright_IL12_d3
ST2+ CD57neg_IL12_d1
ST2+ CD57neg_IL12_d2
ST2+ CD57neg_IL12_d3
ST2+ CD57pos_IL12_d1
ST2+ CD57pos_IL12_d2
ST2+ CD57pos_IL12_d3
ST2- CD56dim CD57neg_IL12_d1
ST2- CD56dim CD57neg_IL12_d2
ST2- CD56dim CD57neg_IL12_d3
ST2- CD56dim CD57pos_IL12_d1
ST2- CD56dim CD57pos_IL12_d2
ST2- CD56dim CD57pos_IL12_d3
n = 2399 genes
row min
row max
d
Normalized expression (DESeq2)
CDC7
DESI1
DMRTA2
FAM89A
FRMD4B
IL1RL1
IL23R
NDFIP2
TRPV1
VEGFA
ST2- CD56bright
ST2+ CD56dim CD57-
ST2+ CD56dim CD57+
ST2- CD56dim CD57-
ST2- CD56dim CD57+

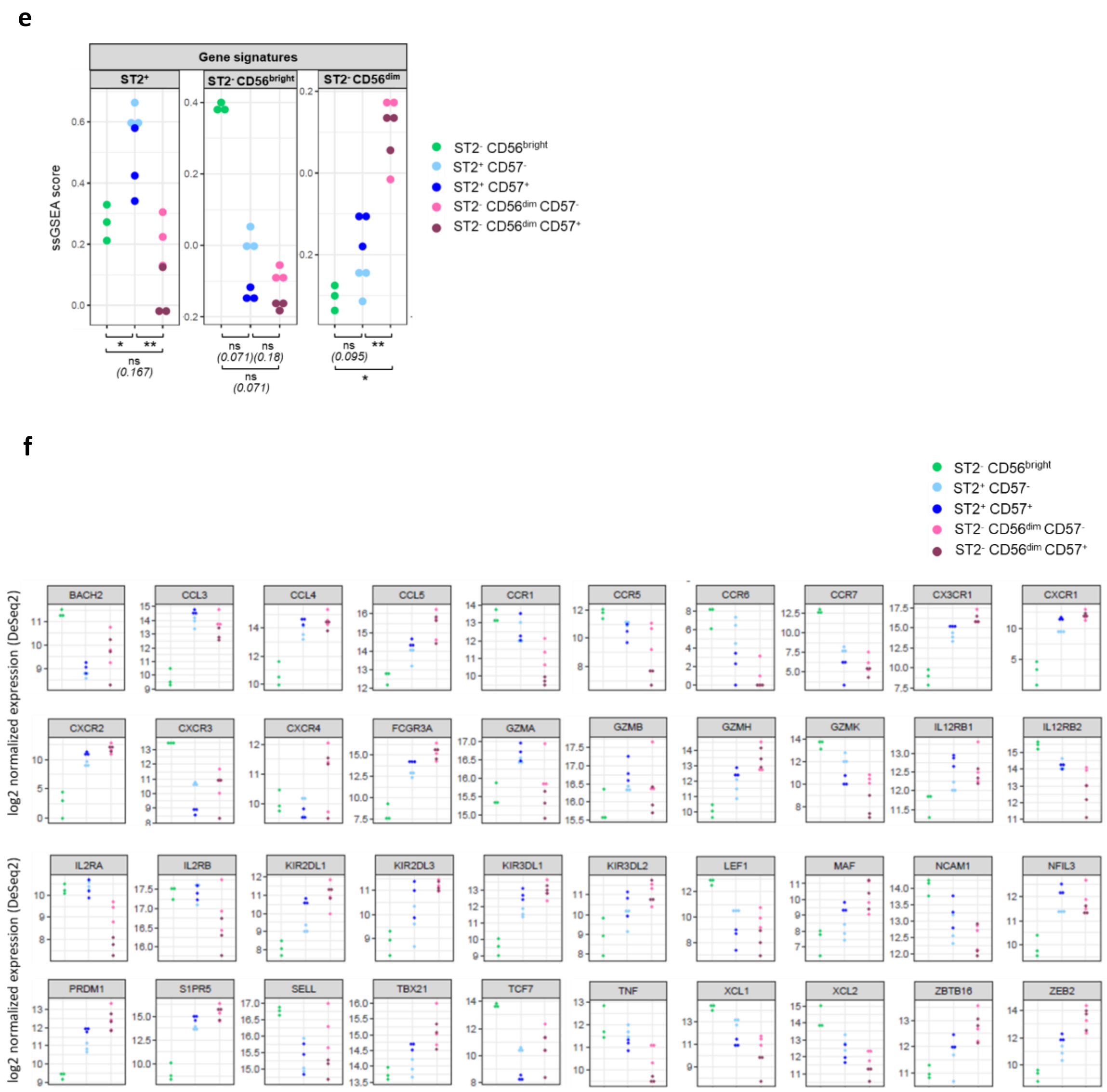


**Supplementary Fig. 4. ST2$^{+}$ NK cells display a unique gene signature compared to CD56$^{bright}$ and CD56$^{dim}$ NK cells**

**a)** Schematic representation of FACS sorting strategy to isolate eight different subsets sorted from healthy donors' blood NK cells based on CD56, CD57 and ST2 surface expression. Three and five subsets were obtained after activation for 24 h with medium or IL-12 respectively.

**b)** Gating strategy to sort CD56$^{bright}$, CD56$^{dim}$ and ST2$^{+}$ NK cells from total blood NK cell suspensions by flow cytometry following activation with Medium or IL-12 for 48 h.

**c)** Heatmap representing all differentially-expressed genes (upregulated and downregulated) (n=2399 genes) in each sorted healthy donors' blood NK cell subset compared to the two other subsets following IL-12 activation (n = 3 individual donors).

**d)** Expression levels of TOP-10 DEG specifically enriched in ST2$^{+}$ NK cells as compared to CD56$^{bright}$ and CD56$^{dim}$ NK cells in each sorted NK cell subset.

**e)** ssGSEA analysis of the NK signatures defined by genes specifically upregulated in each sorted NK cell subset (see Supplementary tables 1-3) following IL-12 activation for 24 h (n = 3 individual healthy donors). Kruskal-Wallis test with Dunn multiple comparisons test was performed.

**f)** Expression levels of DEG (up and down) presented in heatmap in Fig. 3C.

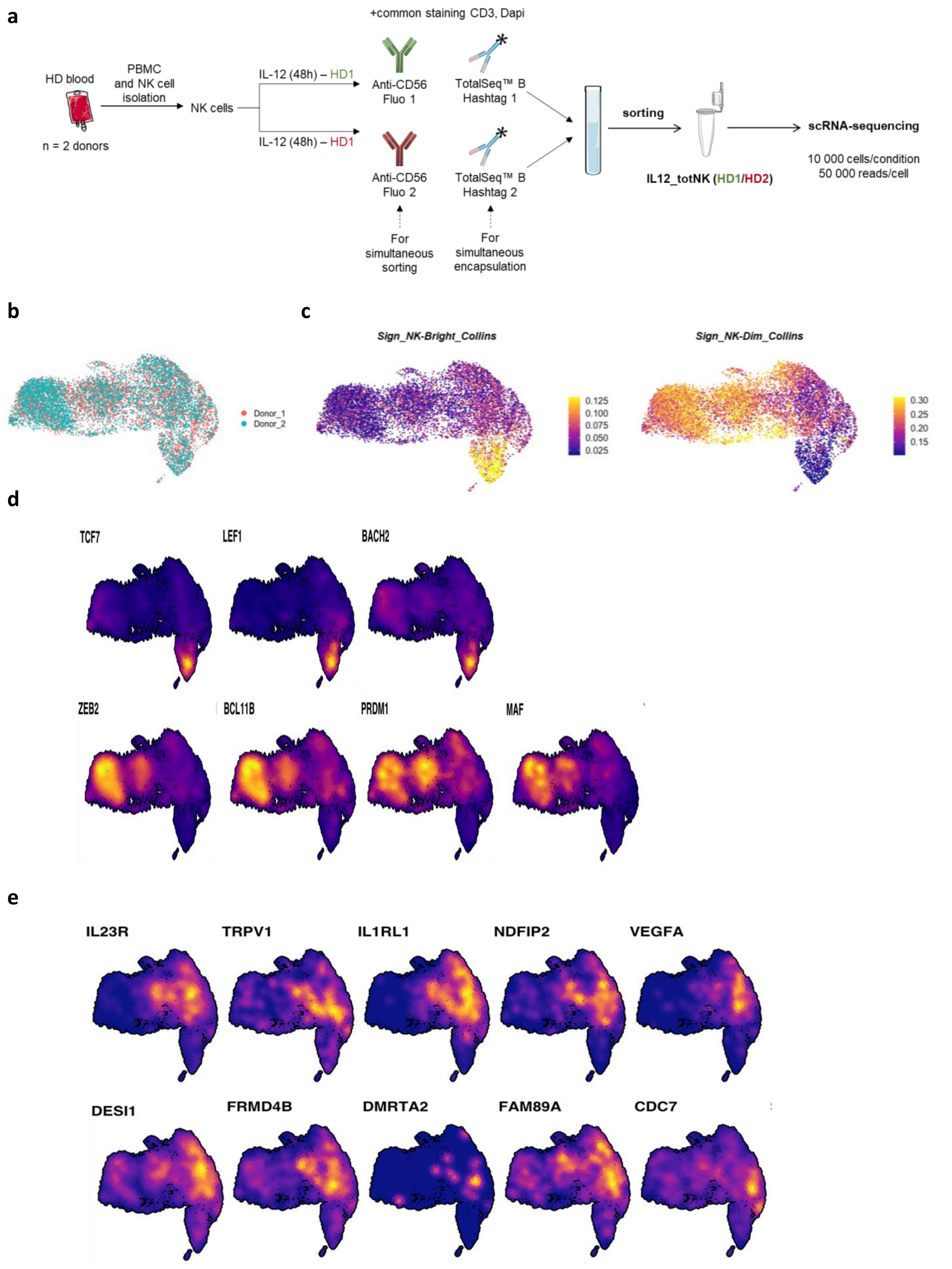


**Supplementary Fig. 5. scRNAseq analysis of IL-12-cultured blood NK cells reveals a distinct subset enriched for the ST2⁺ NK gene signature, characterized by unique transcriptomic features.**

**a)** Schematic representation of strategy to simultaneously sort and encapsulate IL-12 cultured total NK cells from 2 different healthy donors for 48h.

**b)** UMAP representation of blood NK cells from 2 independent donors analyzed by scRNA-seq analysis according to their origin (donor 1 in red, donor 2 in blue).

**c)** UMAP projections of specific gene signatures for CD56bright (left) and CD56dim (right) NK cells identified in Collins *et al.* (16)

**d)** UMAP projections of specific transcription factors of $CD56^{bright}$ (upper panels) and $CD56^{dim}$ (lower panels) NK cells identified in Collins *et al.* (16)

**e)** UMAP projections of TOP10 DEG specifically enriched in $ST2^{+}$ NK cells as compared to $CD56^{bright}$ and $CD56^{dim}$ NK cells from our bulk RNAseq analysis on sorted NK cell subsets (Fig. 3B).

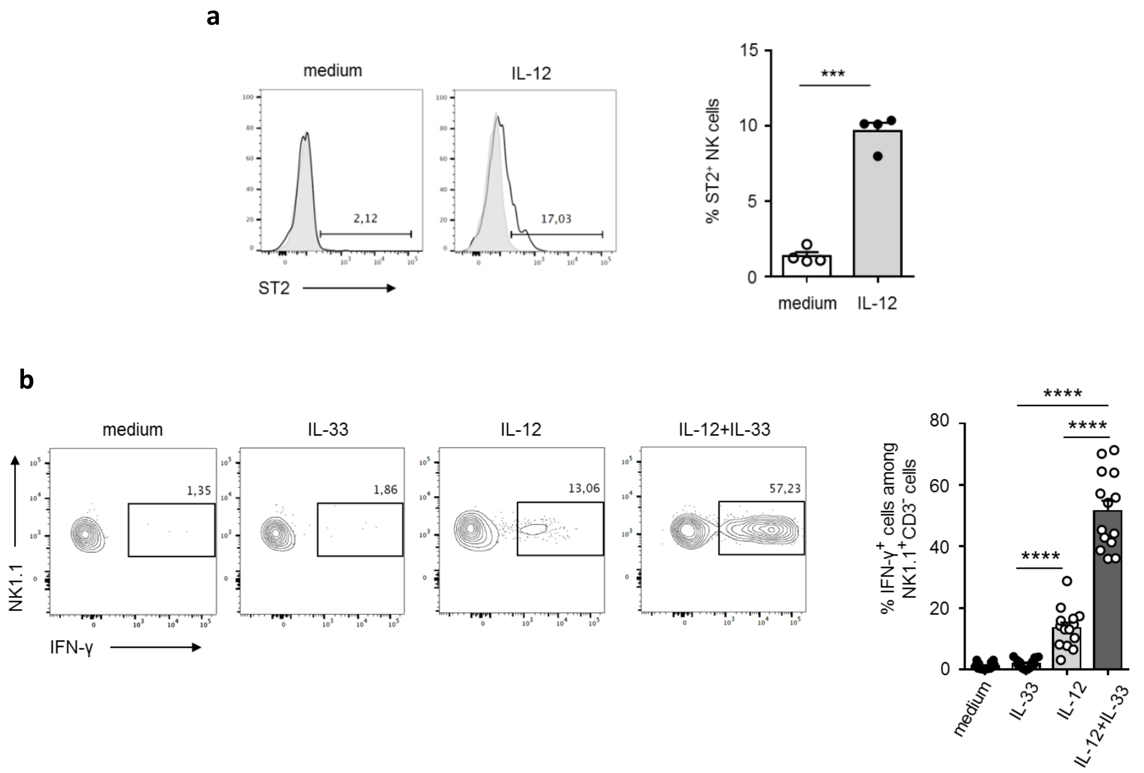


**Supplementary Fig. 6. IL-12 sensitizes mouse splenic NK cells to the production of IFN-γ in response to IL-33**

**a)** Splenic cells were activated with IL-12 (20 ng/mL) or not for 24 h prior to flow cytometry analysis of ST2 surface expression (black line) or corresponding isotypic control (grey). CD45, NK1.1, and CD3 were used to identify NK cells. Representative histogram plots (left) and quantification (%) (right) of $ST2^+$ NK cells after medium or IL-12 culture. Horizontal bars represent the mean + SEM (n = 4 individual mice). Paired two-tailed Student's *t*-test was performed.

**b)** Splenic cells were activated as indicated (20 ng/mL of each cytokine) for 24 h prior to flow cytometry analysis of IFN-γ intracellular expression. CD45, NK1.1, and CD3 were used to identify NK cells. Plots are representative of four individual experiments with 3 to 4 mice/experiment and results for each individual mouse are presented in the right panel. Results are expressed as mean + SEM. One-way repeated measures ANOVA with Tukey's multiple comparisons test was performed.

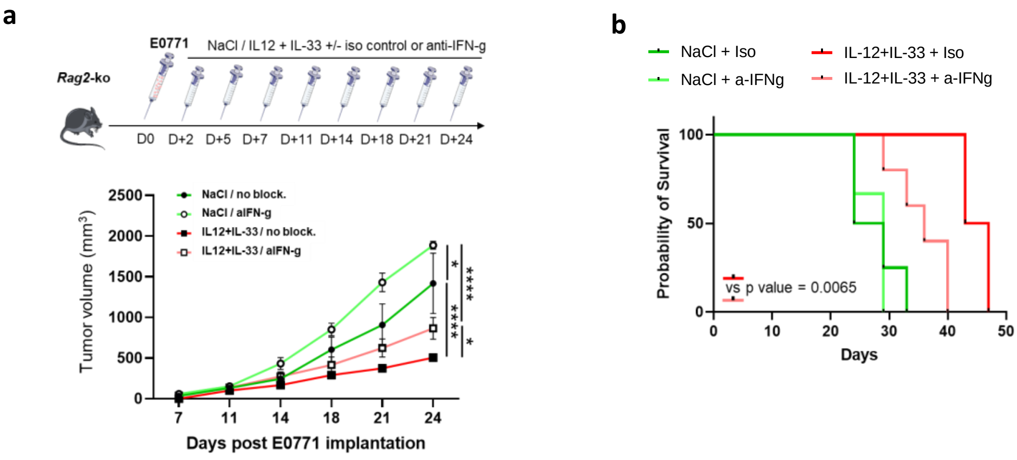


**Supplementary Fig. 7. Antitumoral effect of IL-33 and IL-12 combination is partially dependent on IFN-γ.**

**a)** *Rag2*-KO mice were injected intra-mammary with 2.5x10$^{5}$ E0771 cells on day 0 and then treated in the tumor area with NaCl or a combination of 10 ng/mouse rmIL-12 and 100 ng/mouse rmIL-33 and intraperitoneally with isotype control or blocking anti-IFN-g antibody twice a week from day 2 to day 24. Primary tumor growth was monitored in *Rag2*-KO mice treated with NaCl (n = 4) or with rmIL-12 and rmIL-33 combination (n = 4) or in anti-IFN-g-treated *Rag2*-KO mice treated with NaCl (n = 3) or with rmIL-12 and rmIL-33 combination (n=5). Two-way ANOVA with Tukey's multiple comparisons test was performed.

**b)** Kaplan-Meier survival plots of *Rag2*-KO mice treated with NaCl (n = 4) or with rmIL-12 and rmIL-33 combination (n = 43) or of anti-IFN-γ-treated *Rag2*-KO mice treated with NaCl (n = 3) or with rmIL-12 and rmIL-33 combination (n=5). Mice were sacrificed when longest side of primary tumor reached 17 mm. Log-rank test was performed and the p-value is indicated for *Rag2*-KO mice treated with rmIL-12 + rmIL-33 and isotype control vs anti-IFN-γ antibody mice.

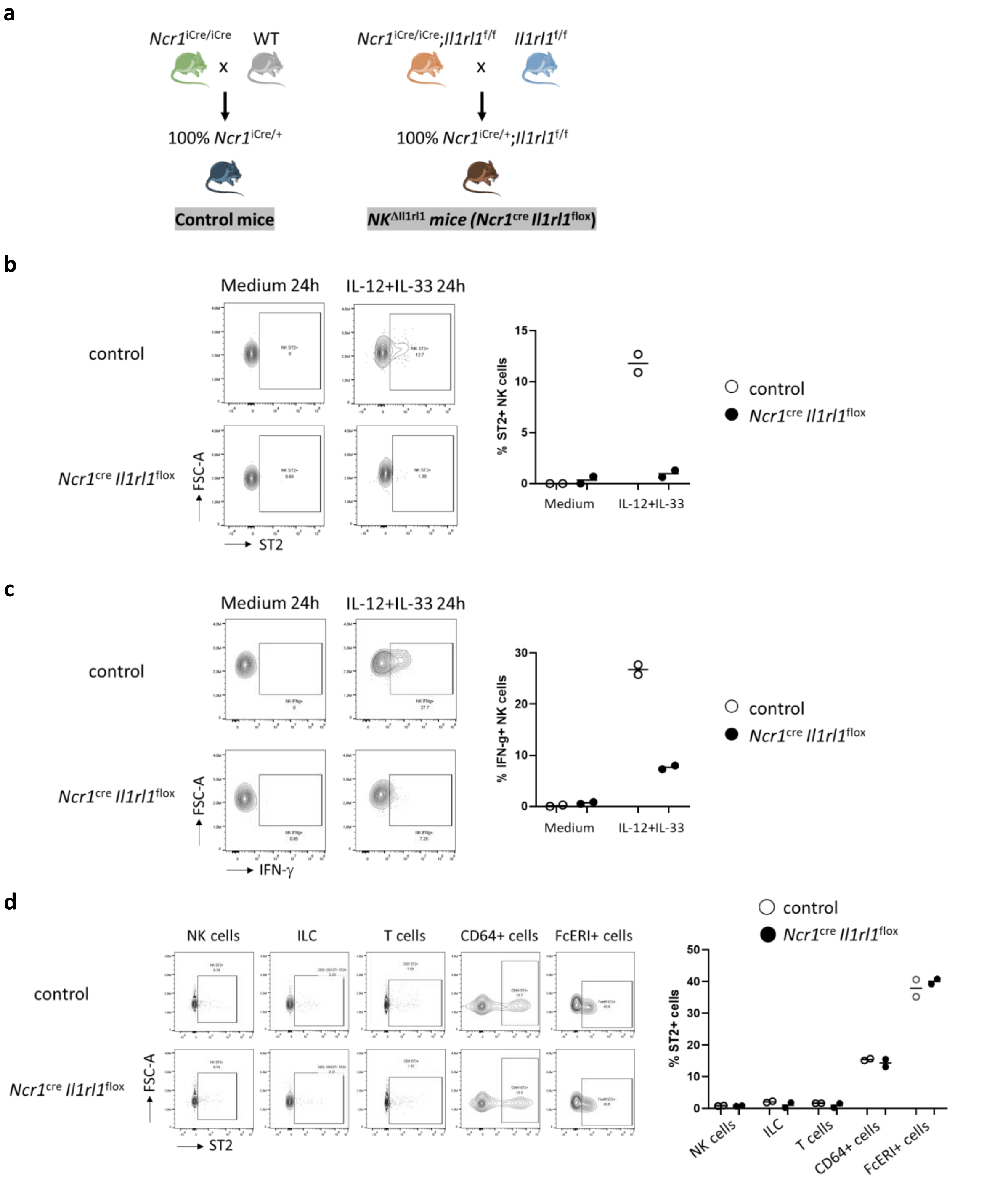


**Supplementary Fig. 8. Validation of *Il1rl1* deletion efficiency in NK cells from the conditional targeting mouse model.**

**a**) Schematic graphs showing the breeding strategies for the conditional knockout mice and their control littermates.

**b**) Analysis of ST2 expression in NK cells from spleen of control (*Ncr1*$^{iCre/+}$) and *Ncr1*$^{cre}$ *Il1rl1*$^{flox}$ mice by flow cytometry after 24h-culture in medium or IL-12+IL-33. Representative dot plots (left) and quantification (%) (right) of ST2$^{+}$ cells after 24h-culture are shown. Horizontal bars represent the median. n = 2 mice per genotype.

**c**) Analysis of IFN-γ expression in NK cells from spleen of control (*Ncr1*$^{iCre/+}$) and *Ncr1*$^{cre}$ *Il1rl1*$^{flox}$ mice by flow cytometry after 24h-culture in medium or IL-12+IL-33. Representative dot plots (left) and quantification (%) (right) of IFN-γ$^{+}$ cells after 24h-culture are shown. Horizontal bars represent the median. n = 2 mice per genotype.

**d**) Analysis of ST2 expression in different immune cells from spleen of control (*Ncr1*$^{iCre/+}$) and *Ncr1*$^{cre}$ *Il1rl1*$^{flox}$ mice by flow cytometry. Representative dot plots (left) and quantification (%) (right) of ST2$^{+}$ cells after isolation from spleen are shown. Horizontal bars represent the median. n = 2 mice per genotype.

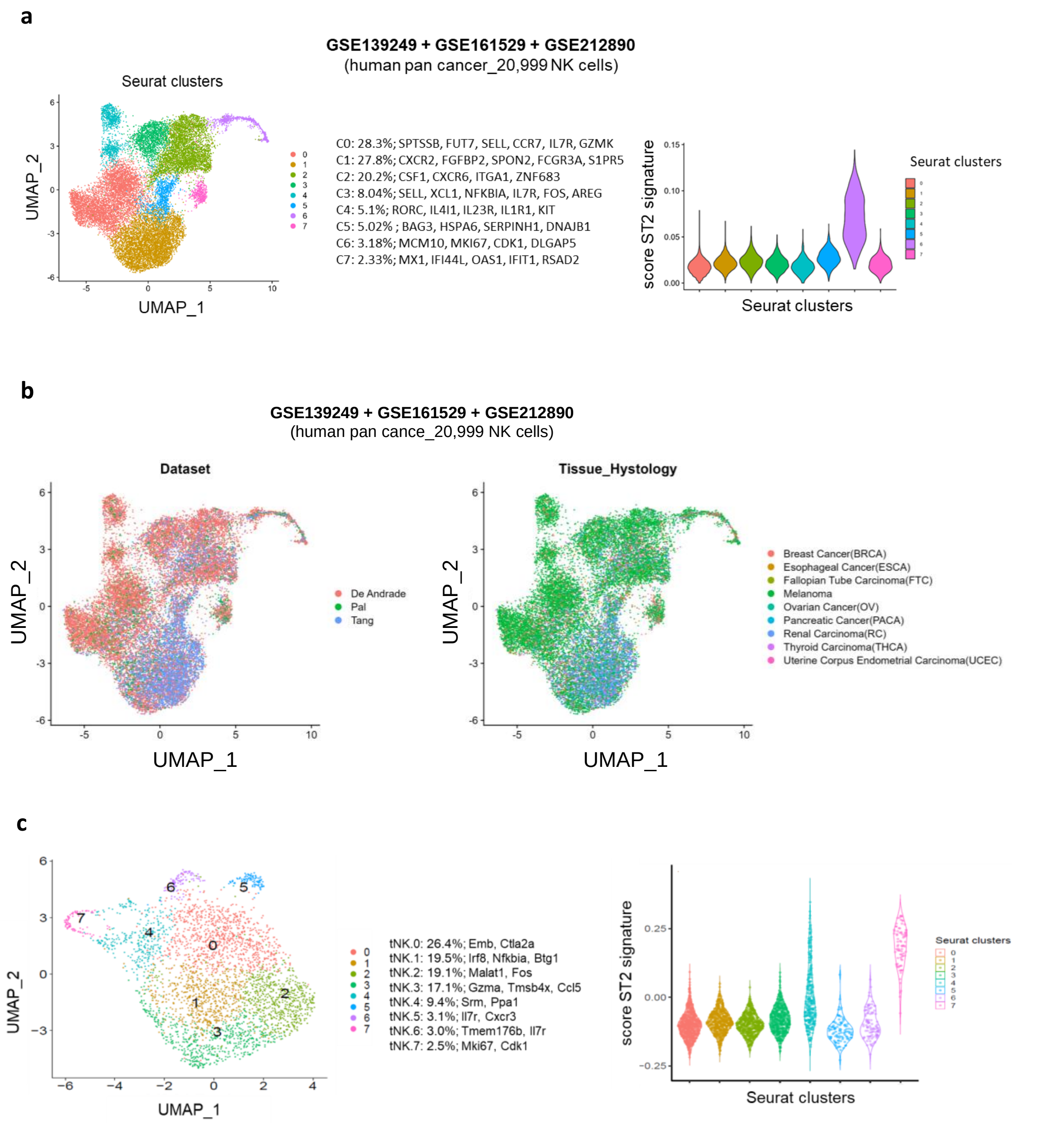


**Supplementary Fig. 9. Unsupervised clustering of public sc-RNAseq datasets of tiNK cells**

**a)** Violin plots representing the distribution of the ST2$^{+}$ NK cells 233-gene signature (see Supplementary table 1) in tiNK cell clusters identified from pan cancer scRNAseq datasets (GSE139249 (36), GSE161529 (37), and GSE212890 (15)).

**b)** UMAP plots of tiNK cells from human tumors sc-RNAseq public datasets (GSE139249 (36), GSE161529 (37), and GSE212890 (15)) according to the study (left) and the tissue histology (right).

**c)** Clustering and UMAP plots (left panel) of NK cells clusters annotated from mouse lymphoma tumors (GSE123534) (38) sc-RNAseq public datasets. Violin plots (right panel) representing the distribution of the human orthologous $ST2^+$ NK cells signature (233 genes, see Supplementary table 1) score in murine tiNK cell clusters.

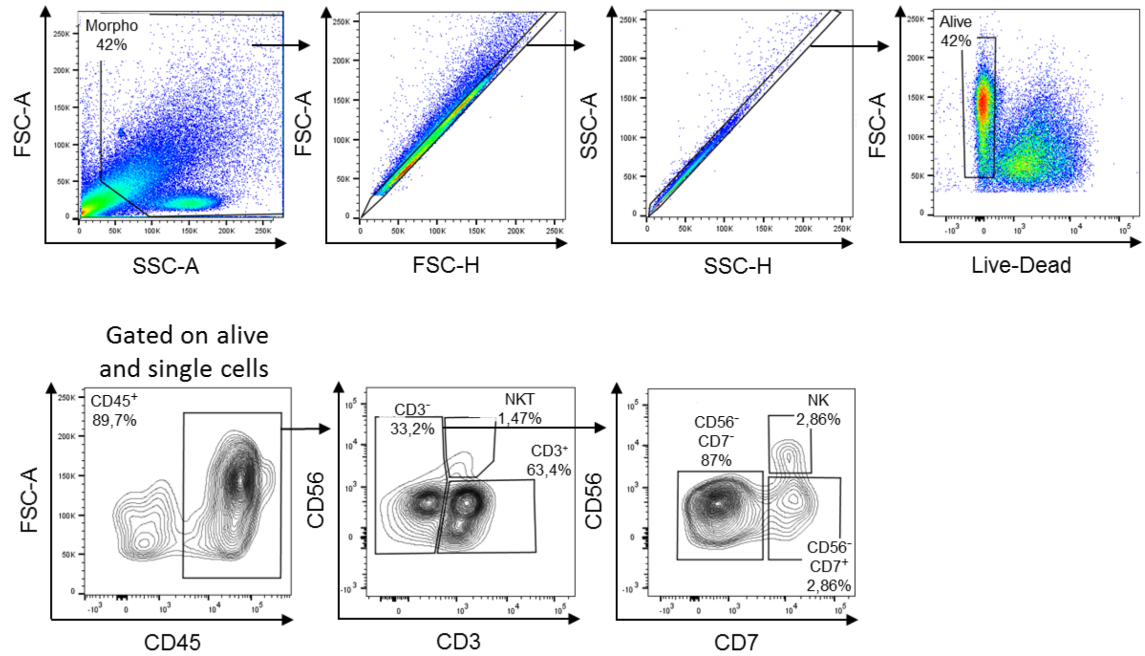


**Supplementary Fig. 10. Gating strategy to identify NK cells in human tumor cell suspensions by flow cytometry**

Tumor cell suspensions were stained for CD45, CD3, CD56, and CD7 expression and tumor-infiltrating NK cells were identified using gating strategy as illustrated. Cell viability was determined using Zombie dye (Biolegend).

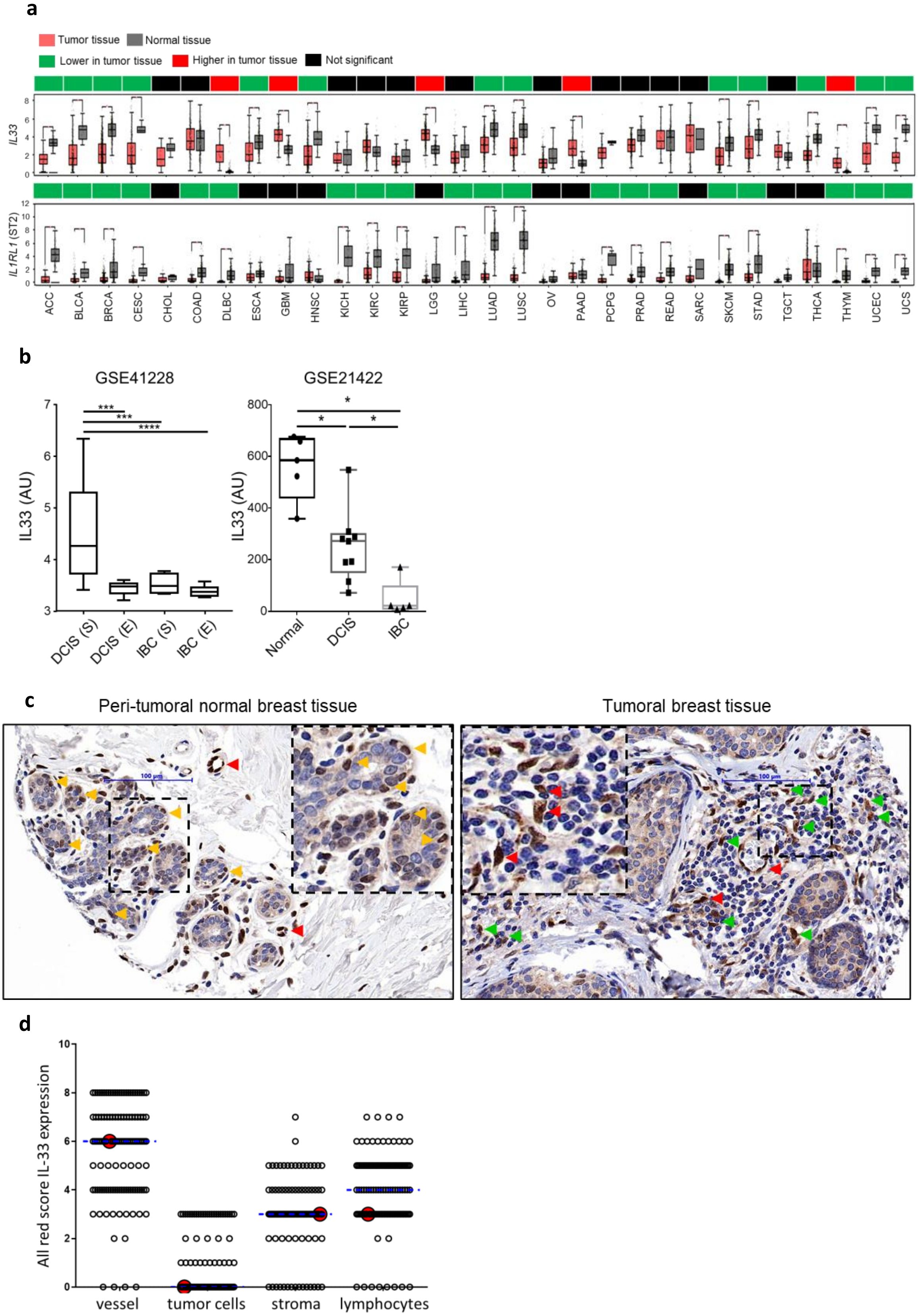


**Supplementary Fig. 11. IL-33 expression decreases during tumor progression and is restricted to vascular cells**

**a,** *IL33* and *IL1RL1* (ST2) gene expression in tumoral (T) (orange) and normal (N) (grey) tissues extracted from TCGA and GTEX databases (ACC: Adrenocortical carcinoma, T=77, N=128; BLCA : Bladder carcinoma, T=404, N=28; BRCA : Breast invasive carcinoma, T=1095, N=291; CESC: Cervical squamous cell carcinoma and endocervical adenocarcinoma, T=306, N=13 ; CHOL: Cholangiocarcinoma, T=36, N=9; COAD : Colorectal adenocarcinoma, T=275, N=349; DLBC: Lymphoid Neoplasm Diffuse Large B-cell Lymphoma, T=47, N=337 ; ESCA: Esophageal carcinoma, T=182, N=286 ; GBM: Glioblastoma multiforme, T=163, N=207 ; HNSC : Head and neck squamous cell carcinoma, T=519, N=44 ; KICH: Kidney Chromophobe, T=66, N=53 ; KIRC : Kidney renal clear cell carcinoma, T=523, N=100 ; KIRP: Kidney renal papillary cell carcinoma, T=286, N=60 ; LGG : Brain lower grade glioma, T=518, N=207 ; LIHC: Liver hepatocellular carcinoma, T=369, N=160 ; LUAD : Lung adenocarcinoma, T=483, N=347 ; LUSC: Lung squamous cell carcinoma, T=486, N=338 ; OV : Ovarian serous cystadenocarcinoma, T=426, N=88 ; PAAD : Pancreatic adenocarcinoma, T=179, N=171 ; PCPG: Pheochromocytoma and Paraganglioma, T=182, N=3 ; PRAD: Prostate adenocarcinoma, T=492, N=152 ; READ: Rectum adenocarcinoma, T=92, N=318 ; SARC: Sarcoma, T=262, N=2 ; SKCM : Skin cutaneous melanoma, T=461, N=558 ; STAD : Stomach adenocarcinoma, T=408, N=211 ; TGCT: Testicular Germ Cell Tumors, T=137, N=165 ; THCA : Thyroid cancer, T=512, N=337; THYM: Thymoma, T=118, N=339 ; UCEC: Uterine Corpus Endometrial Carcinoma, T=174, N=91 ; UCS: Uterine Carcinosarcoma , T=57, N=78). *$p<0.05$

**b,** Analysis of *IL33* expression in laser-microdissected stromal (S) versus epithelial (E) zones from DCIS and IBC lesions (GSE41228) (39) and in healthy mammary tissue versus DCIS and IBC (GSE21422) (40). Kruskal-Wallis test with Steel-Dwas-Fligner multiple comparisons test was performed.

**c,** IL-33 expression was analyzed on breast tumor FFPE slides by Immunohistochemistry (IHC). Non-invasive Ductal Carcinoma *In Situ* (DCIS) and invasive Breast Cancer (IBC) lesions were identified based on anatomopathological tissue observations. Red arrows indicate blood vessels, orange arrows indicate peritumoral normal breast acini and green arrows indicate isolated stromal cells positive for IL-33 staining. Images are shown at 20x magnification. Dotted squares represent enlarges areas.

**d,** Expression of IL-33 quantified in the nuclei of blood vessels, tumor cells, stroma, and lymphocytes from IHC analysis on FFPE breast tumor samples (n=128 patients). The Allred score combines the percentage of positive cells and the intensity of IL-33 nuclear expression. Red dots represent the scores for the patient used to illustrate the staining in supplementary Fig. S11C. Dotted blue lines represent median values in each category.

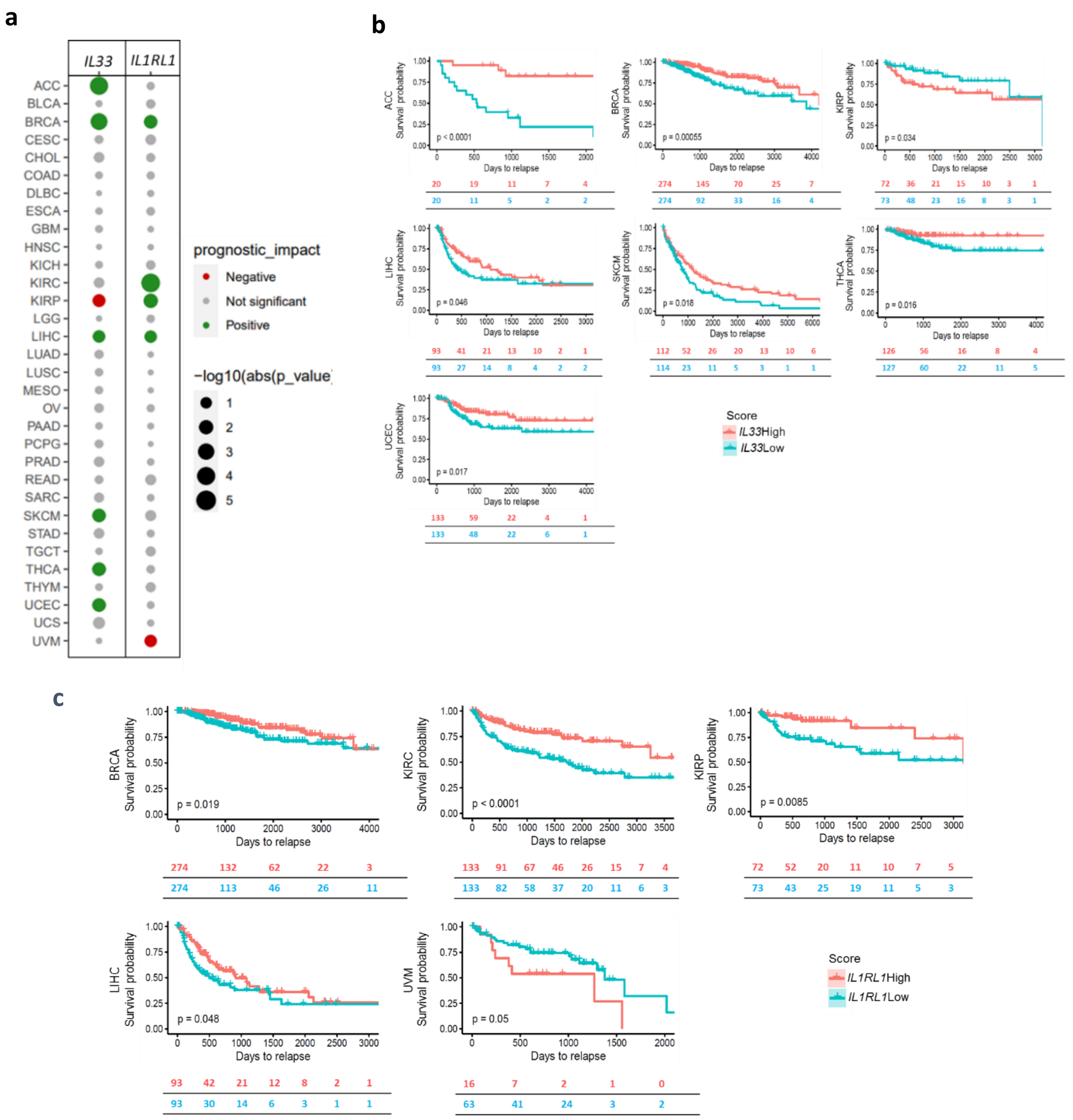


| Variable | | N | Hazard ratio | | p |
|---|---|---|---|---|---|
| **Age_cat** | (25,45] | 150 | | Reference | |
| | (45,65] | 500 | | 0.71 (0.44, 1.16) | 0.171 |
| | (65,75] | 166 | | 0.76 (0.40, 1.45) | 0.406 |
| | (75,90.1] | 106 | | 1.17 (0.58, 2.35) | 0.654 |
| **Molecular_subtype** | Basal | 167 | | Reference | |
| | Her2 | 71 | | 0.80 (0.39, 1.66) | 0.551 |
| | LumA | 495 | | 0.57 (0.35, 0.91) | 0.019 |
| | LumB | 189 | | 0.43 (0.23, 0.80) | 0.008 |
| **Stage** | 1 | 152 | | Reference | |
| | 2 | 536 | | 2.03 (0.95, 4.33) | 0.067 |
| | 3 | 218 | | 4.94 (2.28, 10.70) | <0.001*** |
| | 4 | 16 | | 20.39 (7.94, 52.35) | <0.001*** |
| **IL33_cat** | Low | 229 | | Reference | |
| | Intermediate1 | 227 | | 0.46 (0.26, 0.81) | 0.007** |
| | Intermediate2 | 234 | | 0.65 (0.38, 1.10) | 0.107 |
| | High | 232 | | 0.45 (0.26, 0.79) | 0.006** |

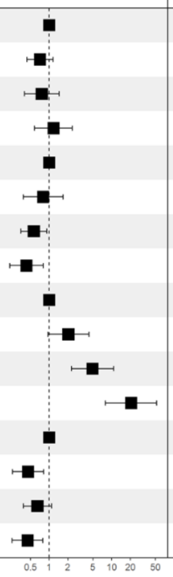

**Supplementary Fig. 12. *IL33* expression correlates with favorable progression-free survival across multiple solid cancers.**

**a)** Summary of p-values associated with the log rank test performed to evaluate prognostic value of *IL33* and *IL1RL1* (ST2) in 32 human TCGA cancer data sets (the same as in Fig. 6C)). Data are represented as a bubble map showing positive (green) or negative (red) impact on progression-free survival. Dots size represents p-values obtained by log-rank test.

**b)** Patients from TCGA database were stratified as high or low for *IL33* and progression-free survival was analyzed in all cancer data sets. Kaplan-Meier survival curves for ACC, BRCA, KIRP, LIHC, SKCM, THCA, and UCEC patients are represented and p-values were obtained with log-rank test.

**c)** Patients from TCGA database were stratified as high or low for *IL1RL1* (ST2) and progression-free survival was analyzed in all cancer data sets. Kaplan-Meier survival curves for BRCA, KIRC, KIRP, LIHC, and UVM patients are represented and p-values were obtained with log-rank test.

**d)** Multivariate Cox analysis of *IL33* impact on prognosis in breast cancers from TCGA dataset regarding the age, molecular subtype, and stage of the tumors p-values were obtained with log-rank test.

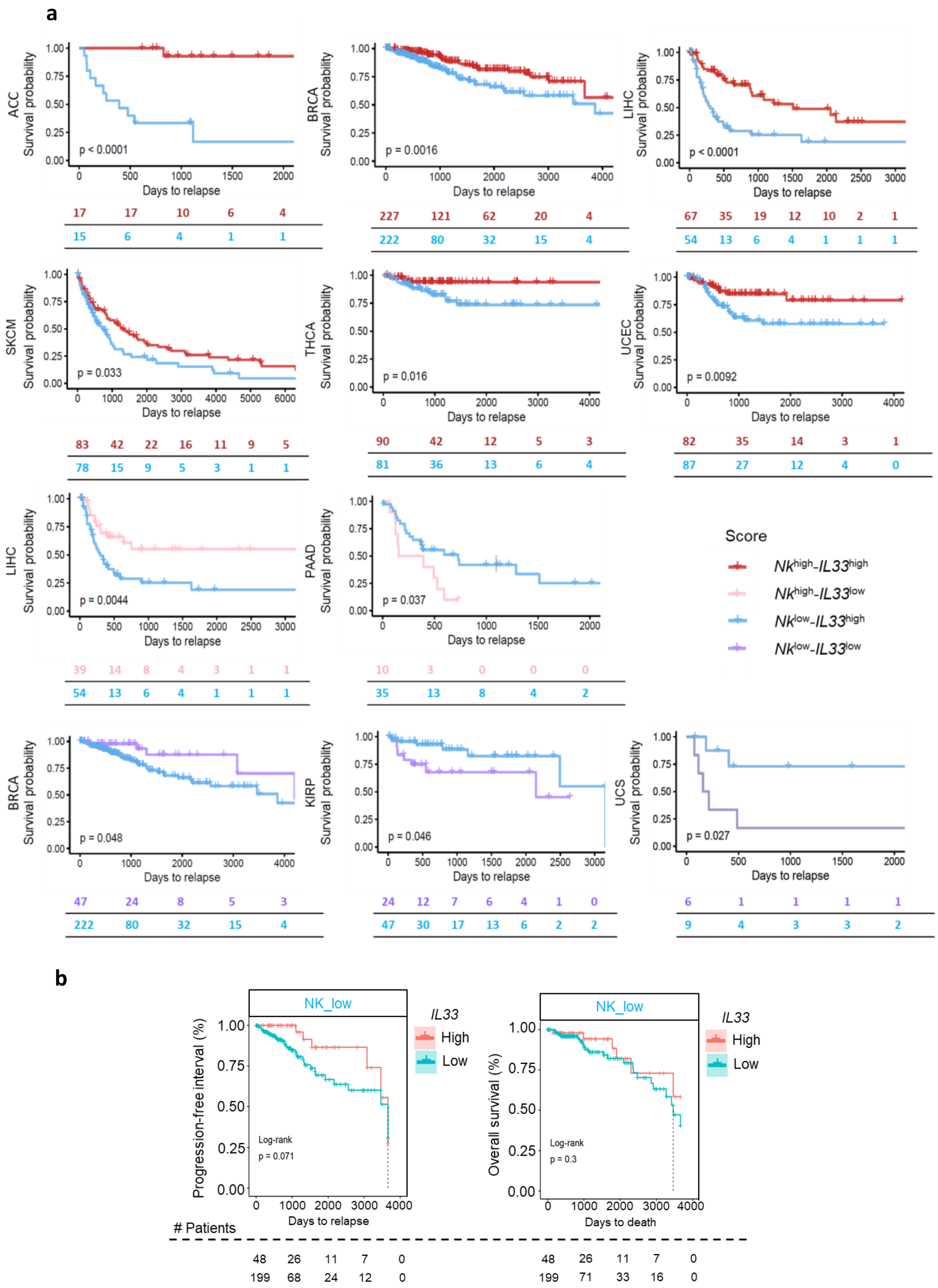


**Supplementary Fig. 13. Pan-cancer survival analysis of the NK cell–IL-33 association in TCGA datasets.**

**a)** Kaplan-Maier curves for progression-free survival for *NK*$^{high}$/*IL33*$^{high}$, *NK*$^{high}$/*IL33*$^{low}$ and *NK*$^{low}$/*IL33*$^{high}$ scores as compared to *NK*$^{low}$/*IL33*$^{low}$ in cancer datasets from TCGA that showed statistical significance in Fig. 6C**.** p-values were obtained with log-rank test. NK signature was defined in (41) (see Supplementary table 8).

**b)** Kaplan-Maier curves for progression-free survival (left panel) and overall survival (right panel) for *NK*$^{low}$ patients stratified according to *IL33* expression in breast cancer dataset from TCGA. p-values were obtained with log-rank test. NK signature was defined in (41) (see Supplementary table 8).

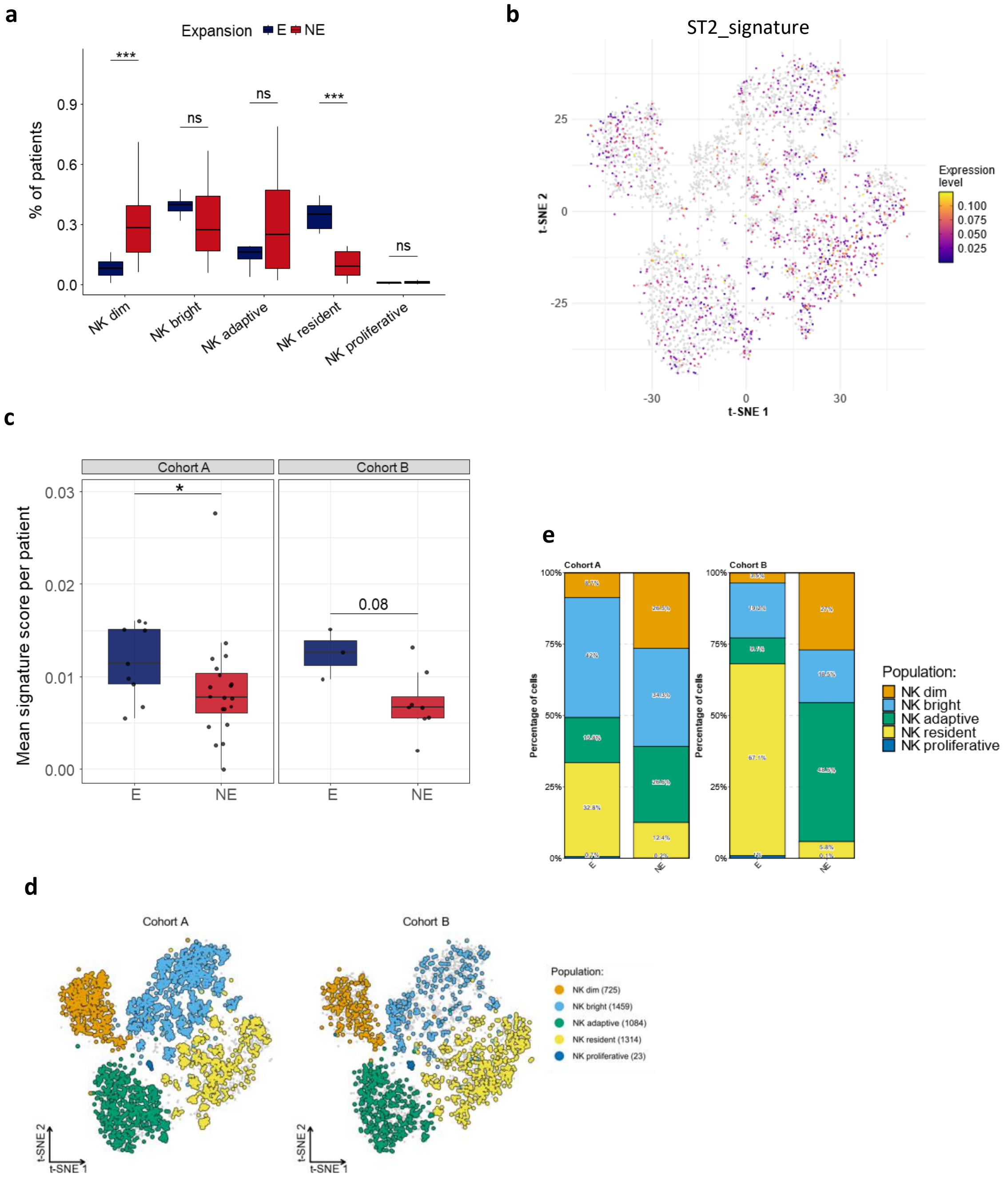


**Supplementary Fig. 14. ST2⁺ NK gene signature contribution across tiNK cell subsets stratified by treatment cohorts and ICB response status.**

**a,** Box plots comparing the relative proportions of patients from Expanded (E) versus Non-Expanded (NE) groups according to the contribution of each cell type, from scRNAseq dataset EGAD00001006608 (42).

**b,** Projection of ST2⁺ NK cell top 10-gene signature enrichment scores (see Supplementary table 1), calculated using the Mann–Whitney U statistic across t-SNE visualization of tiNK, from scRNAseq dataset EGAD00001006608 (42).

**c,** Pseudobulk analysis of mean ST2 NK signature scores per patient according to clonal expansion status, stratified by treatment-arm cohort origin, from scRNAseq dataset EGAD00001006608 (42). Each dot represents the aggregated single-cell expression profile of an individual patient scored for the ST2 NK signature.

**d,** t-SNE plots showing total tiNK cells colored according to subset identity and stratified by treatment-arm cohort origin, from scRNAseq dataset EGAD00001006608 (42).

**e,** Stacked bar plots showing the relative proportions of tiNK subsets according to clonal expansion status and stratified by treatment-arm cohort origin, from scRNAseq dataset EGAD00001006608 (42). Statistical significance was assessed using a two-sided Wilcoxon rank-sum test (*$p$ < 0.05, **$p$ < 0.01, ***$p$ < 0.001).

**Supplementary table 1.** List of $ST2^+$ NK cell signature genes used in this study, Related to Fig. 3B, 3E, 3H, 3I, 5A, 5B, 6H, 6I, Supplementary Fig. S4E, S9A, S9C, S14A-C

| gene | baseMean | log2FoldChange | lfcSE | stat | pvalue | padj |
|---|---|---|---|---|---|---|
| IL23R | 840,503787 | 2,42322895 | 0,24889705 | 9,81765284 | 9,45E-23 | 8,62E-19 |
| TRPV1 | 585,713744 | 1,5368841 | 0,1666148 | 9,18767638 | 4,01E-20 | 1,46E-16 |
| IL1RL1 | 1126,7443 | 2,61252558 | 0,29381202 | 9,01318107 | 2E-19 | 5,21E-16 |
| NDFIP2 | 620,975976 | 1,96393908 | 0,2429768 | 8,14409229 | 3,82E-16 | 4,35E-13 |
| VEGFA | 800,797748 | 2,08492082 | 0,27235779 | 7,70971428 | 1,26E-14 | 9,58E-12 |
| DESI1 | 1249,1091 | 1,1262011 | 0,14710845 | 7,65517735 | 1,93E-14 | 1,34E-11 |
| FRMD4B | 1643,87039 | 1,81739229 | 0,24219981 | 7,53071051 | 5,05E-14 | 2,79E-11 |
| DMRTA2 | 122,468676 | 3,05535882 | 0,41505092 | 7,31571058 | 2,56E-13 | 1,11E-10 |
| FAM89A | 545,929495 | 1,18540785 | 0,16358858 | 7,24271616 | 4,4E-13 | 1,82E-10 |
| CDC7 | 1128,13727 | 1,06925225 | 0,15213337 | 7,02091227 | 2,2E-12 | 6,82E-10 |
| STRIP2 | 140,760635 | 1,29741714 | 0,1881047 | 6,89072171 | 5,55E-12 | 1,33E-09 |
| NCS1 | 468,813606 | 2,07555351 | 0,31228121 | 6,84559957 | 7,62E-12 | 1,76E-09 |
| SLC16A1 | 1512,26695 | 1,9475331 | 0,28782674 | 6,72865566 | 1,71E-11 | 3,55E-09 |
| CLU | 915,149362 | 1,8992889 | 0,28769004 | 6,59283755 | 4,31E-11 | 7,36E-09 |
| CHAC2 | 356,999118 | 1,25300861 | 0,19181008 | 6,53944106 | 6,17E-11 | 1,01E-08 |
| ZNF367 | 984,442668 | 1,89172016 | 0,28799764 | 6,51164619 | 7,43E-11 | 1,18E-08 |
| PSAT1 | 3785,47966 | 1,67376617 | 0,26455322 | 6,44648195 | 1,14E-10 | 1,71E-08 |
| GLB1L2 | 151,981654 | 1,46163105 | 0,22563266 | 6,44192863 | 1,18E-10 | 1,75E-08 |
| WDR76 | 1241,79085 | 1,43758613 | 0,2229631 | 6,43924506 | 1,2E-10 | 1,77E-08 |
| CYCS | 7640,67913 | 1,02715495 | 0,16080527 | 6,38946825 | 1,66E-10 | 2,32E-08 |
| ITGA3 | 253,34202 | 1,30398949 | 0,20420147 | 6,38357806 | 1,73E-10 | 2,39E-08 |
| MT1X | 692,256455 | 1,30721854 | 0,20490144 | 6,36783592 | 1,92E-10 | 2,55E-08 |
| C4orf46 | 658,702544 | 1,03853939 | 0,16290331 | 6,3707641 | 1,88E-10 | 2,55E-08 |
| CDKN2A | 155,551181 | 1,68677994 | 0,26579413 | 6,34882571 | 2,17E-10 | 2,85E-08 |

| | | | | | | |
|---|---|---|---|---|---|---|
| WEE1 | 2447,37932 | 1,44249365 | 0,22840625 | 6,28343908 | 3,31E-10 | 4,14E-08 |
| FANCB | 202,276777 | 1,19938518 | 0,19410873 | 6,18297309 | 6,29E-10 | 7,21E-08 |
| ODC1 | 3337,57699 | 1,214216 | 0,19669728 | 6,16936299 | 6,86E-10 | 7,72E-08 |
| PMAIP1 | 3335,93606 | 1,18029835 | 0,19394516 | 6,13557786 | 8,49E-10 | 9,34E-08 |
| MCM3 | 10000,7616 | 1,06949838 | 0,17412227 | 6,13221471 | 8,67E-10 | 9,35E-08 |
| LIMK2 | 3104,25713 | 1,15832666 | 0,18776431 | 6,12878921 | 8,86E-10 | 9,5E-08 |
| MCM5 | 9509,01304 | 1,19770655 | 0,19494071 | 6,12424439 | 9,11E-10 | 9,66E-08 |
| FANCC | 557,126559 | 1,17140782 | 0,19204919 | 6,10582487 | 1,02E-09 | 1,07E-07 |
| CTNNAL1 | 508,701126 | 1,79673214 | 0,29391591 | 6,09584392 | 1,09E-09 | 1,11E-07 |
| GINS4 | 525,980957 | 1,73755156 | 0,28580647 | 6,05885887 | 1,37E-09 | 1,34E-07 |
| ANKRD9 | 504,828651 | 1,19109652 | 0,19590466 | 6,05394548 | 1,41E-09 | 1,37E-07 |
| GZMA | 74121,1277 | 1,08182451 | 0,17931976 | 6,04154076 | 1,53E-09 | 1,46E-07 |
| NMB | 106,524871 | 1,14403357 | 0,19049991 | 5,99940616 | 1,98E-09 | 1,81E-07 |
| TMEM106C | 2456,27398 | 1,26007271 | 0,21005254 | 5,98257723 | 2,2E-09 | 1,99E-07 |
| MT1E | 44,050544 | 1,95169574 | 0,32328037 | 5,95810717 | 2,55E-09 | 0,00000022 |
| CENPS-CORT | 55,7971453 | 2,46977763 | 0,41342915 | 5,95115155 | 2,66E-09 | 2,27E-07 |
| PRXL2A | 237,702271 | 1,05442176 | 0,17889782 | 5,94511089 | 2,76E-09 | 2,34E-07 |
| MCM7 | 14752,7609 | 1,37191183 | 0,2297699 | 5,94314722 | 2,8E-09 | 2,36E-07 |
| CSTF2 | 2575,74527 | 1,09307847 | 0,18452897 | 5,9067807 | 3,49E-09 | 2,78E-07 |
| PACSIN3 | 25,0768206 | 2,26253594 | 0,38409283 | 5,83189903 | 5,48E-09 | 0,0000004 |
| FAM222A | 180,250612 | 1,40349469 | 0,23981944 | 5,80219939 | 6,55E-09 | 4,57E-07 |
| AL121985.1 | 40,7926711 | 1,16589314 | 0,20293076 | 5,75067325 | 8,89E-09 | 5,85E-07 |
| MARCHF3 | 783,211555 | 1,17359882 | 0,20479762 | 5,73444119 | 9,78E-09 | 6,32E-07 |
| TIMELESS | 4418,15331 | 1,08590411 | 0,18925965 | 5,72959812 | 1,01E-08 | 6,49E-07 |
| AL354718.3 | 26,2533188 | 1,7417743 | 0,29946638 | 5,72789903 | 1,02E-08 | 6,53E-07 |
| ACTN1 | 1511,11114 | 1,74503563 | 0,30449433 | 5,72027976 | 1,06E-08 | 6,79E-07 |
| CCDC74A | 251,973341 | 1,31883746 | 0,2376231 | 5,71927486 | 1,07E-08 | 0,00000068 |
| GINS3 | 467,902103 | 1,33967219 | 0,23328208 | 5,71328543 | 1,11E-08 | 6,94E-07 |

| | | | | | | |
|---|---|---|---|---|---|---|
| H4C2 | 16,5518026 | 1,95322135 | 0,34238088 | 5,70051711 | 1,19E-08 | 7,43E-07 |
| AL133215.2 | 30,1675575 | 1,42266749 | 0,25229359 | 5,63269336 | 1,77E-08 | 0,00000101 |
| WDHD1 | 1158,87928 | 1,46188406 | 0,26192838 | 5,5726197 | 2,51E-08 | 0,00000137 |
| PPIF | 4031,87853 | 1,15942507 | 0,20940214 | 5,55032627 | 2,85E-08 | 0,00000153 |
| TUBB | 35257,5275 | 1,17169715 | 0,21471365 | 5,44816951 | 5,09E-08 | 0,00000244 |
| PTGER3 | 1122,67732 | 1,94573883 | 0,35633685 | 5,42295615 | 5,86E-08 | 0,00000274 |
| NUDT1 | 1001,90806 | 1,00733025 | 0,18719371 | 5,38299465 | 7,33E-08 | 0,00000331 |
| CCDC167 | 992,092596 | 1,07088026 | 0,20061089 | 5,34769926 | 8,91E-08 | 0,00000382 |
| PGD | 6735,92045 | 1,48019965 | 0,27474308 | 5,3438772 | 9,1E-08 | 0,00000388 |
| AL645939.6 | 104,812089 | 1,27138805 | 0,23475732 | 5,33692254 | 9,45E-08 | 0,00000396 |
| RPP25 | 367,831914 | 1,47390104 | 0,27692459 | 5,32409178 | 1,01E-07 | 0,00000418 |
| HDAC9 | 472,156038 | 1,47723458 | 0,28428291 | 5,3202141 | 1,04E-07 | 0,00000427 |
| RPS6KL1 | 45,2426705 | 1,51043197 | 0,28647669 | 5,30331887 | 1,14E-07 | 0,00000452 |
| C17orf58 | 532,027454 | 1,01441751 | 0,19141547 | 5,30104302 | 1,15E-07 | 0,00000454 |
| SLC43A3 | 1071,36813 | 1,13990933 | 0,21499718 | 5,27929886 | 0,00000013 | 0,00000504 |
| SNORD3A | 146,092244 | 1,13049166 | 0,21362145 | 5,27808359 | 1,31E-07 | 0,00000506 |
| MTFP1 | 1468,3341 | 1,157943 | 0,21913998 | 5,27760423 | 1,31E-07 | 0,00000506 |
| ZNRF1 | 3820,68738 | 1,16731492 | 0,2234197 | 5,2740492 | 1,33E-07 | 0,00000511 |
| PPFIA3 | 324,368873 | 1,55820112 | 0,29729033 | 5,27239408 | 1,35E-07 | 0,00000515 |
| UCK2 | 2079,14805 | 1,13481736 | 0,21676644 | 5,22789662 | 1,71E-07 | 0,00000624 |
| AC011447.3 | 57,3186488 | 1,11553004 | 0,21343931 | 5,22242262 | 1,77E-07 | 0,00000638 |
| HASPIN | 554,208648 | 1,68583729 | 0,32282234 | 5,21282615 | 1,86E-07 | 0,00000661 |
| ULBP1 | 149,625849 | 1,51410848 | 0,29748326 | 5,21097287 | 1,88E-07 | 0,00000665 |
| SLC6A9 | 187,199925 | 1,70333199 | 0,33635265 | 5,18997273 | 0,00000021 | 0,00000735 |
| RPA3 | 1636,73819 | 1,00311422 | 0,19366157 | 5,1705612 | 2,33E-07 | 0,00000798 |
| ARHGAP11B | 278,264429 | 1,20368016 | 0,23432331 | 5,16331317 | 2,43E-07 | 0,00000825 |
| CHAC1 | 481,352013 | 1,23939592 | 0,24088971 | 5,14265704 | 2,71E-07 | 0,00000902 |
| CDK4 | 3873,60144 | 1,11749802 | 0,21784441 | 5,12557997 | 2,97E-07 | 0,00000971 |

| | | | | | | |
|---|---|---|---|---|---|---|
| LRR1 | 920,996475 | 1,0036983 | 0,19575589 | 5,12386851 | 2,99E-07 | 0,00000974 |
| AP1S1 | 105,27595 | 1,0450958 | 0,20455019 | 5,11990457 | 3,06E-07 | 0,00000991 |
| SHMT2 | 9828,60995 | 1,1975069 | 0,23618809 | 5,10614466 | 3,29E-07 | 0,0000106 |
| DNAJC9 | 3188,18598 | 1,22199097 | 0,2387053 | 5,09616228 | 3,47E-07 | 0,000011 |
| PSMC3IP | 350,481732 | 1,4581779 | 0,28547784 | 5,0661719 | 4,06E-07 | 0,0000126 |
| GLP1R | 296,527103 | 1,49740713 | 0,30734149 | 5,03306041 | 4,83E-07 | 0,0000144 |
| ICAM1 | 6843,1637 | 1,04169586 | 0,20732701 | 5,03330967 | 4,82E-07 | 0,0000144 |
| TCF19 | 2640,77586 | 1,43029337 | 0,2832562 | 5,02682044 | 4,99E-07 | 0,0000147 |
| CHPF | 3180,3357 | 1,13623626 | 0,22647359 | 5,02329478 | 5,08E-07 | 0,000015 |
| UACA | 101,587297 | 1,28522978 | 0,25825954 | 4,97927636 | 6,38E-07 | 0,0000182 |
| HPSE | 246,082026 | 1,25104351 | 0,24899855 | 4,96053958 | 7,03E-07 | 0,0000195 |
| ZNF90 | 85,3883948 | 1,07121192 | 0,21718338 | 4,94198616 | 7,73E-07 | 0,0000211 |
| CHD5 | 40,98575 | 2,40552677 | 0,45150459 | 4,93564678 | 7,99E-07 | 0,0000216 |
| STEAP1 | 38,756016 | 1,82746385 | 0,36064085 | 4,92882198 | 8,27E-07 | 0,000022 |
| TMEM97 | 753,470908 | 1,44688755 | 0,29464752 | 4,9218635 | 8,57E-07 | 0,0000226 |
| PCNA | 5824,63591 | 1,46770467 | 0,29734296 | 4,91964178 | 8,67E-07 | 0,0000228 |
| DHCR7 | 2278,22228 | 1,01502954 | 0,20677943 | 4,91212558 | 9,01E-07 | 0,0000235 |
| ATAD2 | 3755,90616 | 1,2943769 | 0,26245318 | 4,90082092 | 9,54E-07 | 0,0000246 |
| CKS2 | 1407,44132 | 1,31951811 | 0,26895317 | 4,89882553 | 9,64E-07 | 0,0000248 |
| DSN1 | 1613,8097 | 1,02750526 | 0,21012032 | 4,88135299 | 0,00000105 | 0,0000265 |
| POLD3 | 1710,94088 | 1,00011605 | 0,20433338 | 4,88106533 | 0,00000106 | 0,0000265 |
| PHLDA3 | 77,6403238 | 1,36984895 | 0,27522185 | 4,87388501 | 0,00000109 | 0,0000272 |
| AC079209.1 | 41,3225837 | 1,8582839 | 0,39181777 | 4,86322585 | 0,00000115 | 0,0000283 |
| CENPP | 386,136493 | 1,29006839 | 0,26693782 | 4,84742954 | 0,00000125 | 0,0000303 |
| PKM | 24972,8962 | 1,13454211 | 0,23415197 | 4,83603007 | 0,00000132 | 0,0000319 |
| TESC | 1490,65093 | 1,18929034 | 0,2514482 | 4,83364582 | 0,00000134 | 0,0000322 |
| MCM6 | 4653,0983 | 1,57235012 | 0,32417432 | 4,82704181 | 0,00000139 | 0,000033 |
| MSH6 | 3869,11953 | 1,05660942 | 0,21928593 | 4,80328372 | 0,00000156 | 0,0000361 |

| | | | | | | |
|---|---|---|---|---|---|---|
| RFC4 | 1553,09963 | 1,031953 | 0,21441157 | 4,78866885 | 0,00000168 | 0,0000385 |
| CHAF1A | 2149,62629 | 1,28745474 | 0,27220876 | 4,72032172 | 0,00000235 | 0,0000514 |
| H2AC13 | 51,7125691 | 1,05210436 | 0,22268468 | 4,70945125 | 0,00000248 | 0,0000537 |
| SPTY2D1OS | 42,2396604 | 1,29347563 | 0,27513413 | 4,70686789 | 0,00000252 | 0,0000543 |
| GPT2 | 960,333544 | 1,47635661 | 0,30788401 | 4,69049231 | 0,00000273 | 0,0000578 |
| MCM4 | 6340,26235 | 1,63367485 | 0,34634032 | 4,68454086 | 0,00000281 | 0,0000591 |
| NDC80 | 776,695235 | 1,29472715 | 0,27563296 | 4,67623182 | 0,00000292 | 0,000061 |
| GMNN | 934,054353 | 1,43630881 | 0,30640338 | 4,67506347 | 0,00000294 | 0,0000613 |
| E2F2 | 802,219169 | 1,65972969 | 0,3514081 | 4,66431846 | 0,0000031 | 0,0000641 |
| P2RX5-TAX1BP3 | 1342,34518 | 1,52101022 | 0,3204367 | 4,66403728 | 0,0000031 | 0,0000642 |
| PDLIM1 | 1187,47176 | 1,03361573 | 0,2224437 | 4,6617062 | 0,00000314 | 0,0000647 |
| TMEM154 | 574,226839 | 1,00573313 | 0,2150905 | 4,64587086 | 0,00000339 | 0,0000683 |
| CHEK1 | 825,482205 | 1,73191025 | 0,36953111 | 4,64538288 | 0,00000339 | 0,0000683 |
| TREML2 | 235,222525 | 1,45436846 | 0,32219791 | 4,64228458 | 0,00000345 | 0,000069 |
| EME1 | 314,144262 | 1,57289255 | 0,33841067 | 4,62692244 | 0,00000371 | 0,0000731 |
| MCM2 | 8689,40083 | 1,61040092 | 0,34557034 | 4,62485422 | 0,00000375 | 0,0000737 |
| GINS2 | 1161,25528 | 1,8507702 | 0,39762798 | 4,61960492 | 0,00000384 | 0,0000753 |
| ORC6 | 684,604254 | 1,57585647 | 0,33937762 | 4,61923814 | 0,00000385 | 0,0000753 |
| EPOP | 77,9565208 | 1,32065759 | 0,28593559 | 4,61440444 | 0,00000394 | 0,0000769 |
| POLD2 | 3182,659 | 1,11906962 | 0,24281803 | 4,60941177 | 0,00000404 | 0,0000783 |
| TMPO-AS1 | 283,780703 | 1,71981254 | 0,36963206 | 4,59856749 | 0,00000425 | 0,0000818 |
| IFNG | 6396,66052 | 1,45636818 | 0,31848246 | 4,58996006 | 0,00000443 | 0,0000847 |
| RASD1 | 606,327077 | 1,55681397 | 0,33333489 | 4,58647174 | 0,00000451 | 0,0000858 |
| DHCR24 | 7856,12245 | 1,11331258 | 0,24256226 | 4,58412271 | 0,00000456 | 0,0000863 |
| TFDP1 | 4133,97738 | 1,30153913 | 0,28366491 | 4,57958167 | 0,00000466 | 0,0000877 |
| AP1S3 | 369,438627 | 1,3518563 | 0,29480193 | 4,56559322 | 0,00000498 | 0,0000928 |
| BRCA1 | 930,477735 | 1,39840757 | 0,30518946 | 4,56130619 | 0,00000508 | 0,0000945 |
| SLC7A1 | 2343,405 | 1,10529514 | 0,24349859 | 4,55492539 | 0,00000524 | 0,0000965 |

| | | | | | | |
|---|---|---|---|---|---|---|
| DHFR | 2151,02731 | 1,55704343 | 0,33954021 | 4,5402195 | 0,00000562 | 0,00010235 |
| ATAD5 | 805,154511 | 1,10530992 | 0,2427956 | 4,53887588 | 0,00000566 | 0,00010269 |
| RANP1 | 16,5239844 | 1,32440045 | 0,28989191 | 4,51475446 | 0,00000634 | 0,00011231 |
| BZW1P2 | 22,4181532 | 1,12484237 | 0,47018698 | 4,51482661 | 0,00000634 | 0,00011231 |
| S100P | 87,3413641 | 1,82640658 | 0,3899944 | 4,51308538 | 0,00000639 | 0,00011298 |
| STAP2 | 242,784642 | 1,09058482 | 0,24286098 | 4,51313801 | 0,00000639 | 0,00011298 |
| FEN1 | 5528,4219 | 1,38578289 | 0,30732008 | 4,49108564 | 0,00000709 | 0,00012315 |
| E2F1 | 2025,17146 | 1,7319208 | 0,38283899 | 4,47953839 | 0,00000748 | 0,00012825 |
| AMOTL1 | 496,757411 | 1,02596064 | 0,22667277 | 4,47446543 | 0,00000766 | 0,00013064 |
| TUBAP2 | 38,2765439 | 1,53130728 | 0,34105492 | 4,47299192 | 0,00000771 | 0,00013098 |
| DTYMK | 2010,46089 | 1,02403528 | 0,22848002 | 4,47138208 | 0,00000777 | 0,00013168 |
| NET1 | 461,954204 | 1,2765251 | 0,28560765 | 4,46968512 | 0,00000783 | 0,00013211 |
| CENPN | 782,642068 | 1,53981103 | 0,34494963 | 4,46054026 | 0,00000818 | 0,00013737 |
| AC092718.4 | 63,7504111 | 1,4920021 | 0,33635788 | 4,45531478 | 0,00000838 | 0,0001405 |
| LIPG | 5,89092867 | 1,93128208 | 0,49660756 | 4,42884074 | 0,00000947 | 0,00015449 |
| AC002116.2 | 25,5918918 | 1,19588495 | 0,27276091 | 4,41044863 | 0,0000103 | 0,00016411 |
| UNG | 2362,03983 | 1,32634165 | 0,29975081 | 4,40692892 | 0,0000105 | 0,00016607 |
| RAD54B | 266,75802 | 1,16939837 | 0,26554358 | 4,39421167 | 0,0000111 | 0,00017323 |
| ORC1 | 691,813587 | 1,6666604 | 0,37795945 | 4,3824423 | 0,0000117 | 0,0001824 |
| WDR62 | 766,596341 | 1,45094258 | 0,33134967 | 4,37240158 | 0,0000123 | 0,00018922 |
| CLSPN | 1091,54245 | 1,67501566 | 0,38017063 | 4,36742483 | 0,0000126 | 0,00019244 |
| SCD | 1345,74203 | 1,35532059 | 0,31142925 | 4,35959145 | 0,000013 | 0,00019813 |
| WWC1 | 9,77638638 | 1,85241231 | 0,49501321 | 4,3548267 | 0,0000133 | 0,00020148 |
| SOCS1 | 5198,56908 | 1,17308325 | 0,27012128 | 4,34892689 | 0,0000137 | 0,00020477 |
| RNF157 | 4457,47736 | 1,28354426 | 0,30207097 | 4,33638366 | 0,0000145 | 0,00021452 |
| TUBBP1 | 8,52784924 | 1,50653657 | 0,34610586 | 4,33590032 | 0,0000145 | 0,00021481 |
| FANCI | 4280,14541 | 1,21764881 | 0,28034532 | 4,33554899 | 0,0000145 | 0,00021498 |
| AC125611.3 | 368,112707 | 1,08632677 | 0,25174306 | 4,32071096 | 0,0000156 | 0,00022702 |

| | | | | | | |
|---|---|---|---|---|---|---|
| CHAF1B | 957,421705 | 1,31490521 | 0,30328712 | 4,31485291 | 0,000016 | 0,00023054 |
| BAG2 | 521,183057 | 1,05120342 | 0,24327784 | 4,30709311 | 0,0000165 | 0,00023727 |
| CENPH | 576,293978 | 1,26026279 | 0,2925594 | 4,30587288 | 0,0000166 | 0,0002382 |
| DUT | 4484,90877 | 1,2199032 | 0,28278906 | 4,29737298 | 0,0000173 | 0,00024617 |
| EIF4EBP1 | 1072,5969 | 1,20523444 | 0,27988365 | 4,29345192 | 0,0000176 | 0,00024939 |
| FOXRED2 | 439,287578 | 1,4681439 | 0,34151791 | 4,2862685 | 0,0000182 | 0,00025619 |
| ASRGL1 | 198,029471 | 1,164368 | 0,2682786 | 4,28482002 | 0,0000183 | 0,00025747 |
| PIR | 89,949917 | 1,65357487 | 0,38886029 | 4,27296141 | 0,0000193 | 0,00026824 |
| AL353135.2 | 14,2246629 | 1,62814621 | 0,3797066 | 4,26149956 | 0,0000203 | 0,00027982 |
| ADM2 | 283,026873 | 1,01121121 | 0,23931693 | 4,24890986 | 0,0000215 | 0,0002927 |
| PSPH | 366,751383 | 1,07179051 | 0,25048272 | 4,23729156 | 0,0000226 | 0,00030506 |
| GGH | 647,481775 | 1,55943325 | 0,36825399 | 4,23288001 | 0,0000231 | 0,00031019 |
| LRP8 | 2217,7952 | 1,10561244 | 0,26234263 | 4,22833651 | 0,0000235 | 0,00031513 |
| TLCD3A | 160,982726 | 1,43553228 | 0,33736838 | 4,22412508 | 0,000024 | 0,00031955 |
| SMTN | 270,762571 | 1,126887 | 0,26663656 | 4,22404222 | 0,000024 | 0,00031955 |
| TUBA1B | 60578,4037 | 1,20501359 | 0,2849711 | 4,22013433 | 0,0000244 | 0,00032373 |
| KNTC1 | 1787,11296 | 1,11477093 | 0,26350939 | 4,22033697 | 0,0000244 | 0,00032373 |
| KIF18A | 417,335287 | 1,31788213 | 0,31169429 | 4,21896245 | 0,0000245 | 0,0003247 |
| AC004816.1 | 14,306372 | 1,65661376 | 0,39618463 | 4,202751 | 0,0000264 | 0,0003456 |
| MASTL | 864,89905 | 1,02468418 | 0,24379548 | 4,18746011 | 0,0000282 | 0,00036448 |
| DTL | 2028,0242 | 1,63425478 | 0,39575132 | 4,13128745 | 0,0000361 | 0,00044953 |
| POLE2 | 409,461561 | 1,53115468 | 0,369125 | 4,1304297 | 0,0000362 | 0,00044998 |
| TUBG1 | 2203,28342 | 1,02452182 | 0,24807305 | 4,1290456 | 0,0000364 | 0,00045177 |
| FAM72A | 220,279994 | 1,15187674 | 0,28159687 | 4,12615562 | 0,0000369 | 0,00045593 |
| TEDC2 | 431,086628 | 1,64867763 | 0,39625457 | 4,12497492 | 0,0000371 | 0,00045766 |
| P2RX5 | 22126,469 | 1,50476936 | 0,3635052 | 4,12437677 | 0,0000372 | 0,00045823 |
| ARHGAP11A | 1730,04945 | 1,25299442 | 0,30371393 | 4,12055956 | 0,0000378 | 0,00046369 |
| CDC25A | 699,947246 | 1,81511065 | 0,44397787 | 4,10696078 | 0,0000401 | 0,00048693 |

| | | | | | | |
|---|---|---|---|---|---|---|
| EXO1 | 713,754785 | 1,68789674 | 0,40358897 | 4,10386851 | 0,0000406 | 0,00049185 |
| TUBA1C | 3622,14416 | 1,09707551 | 0,26829002 | 4,09586354 | 0,0000421 | 0,0005068 |
| B4GALNT1 | 10,60565 | 2,15643736 | 0,49332317 | 4,08855336 | 0,0000434 | 0,00052029 |
| CCDC74B | 104,686214 | 1,11756828 | 0,31918457 | 4,08830793 | 0,0000435 | 0,00052045 |
| EZH2 | 1948,0683 | 1,01462741 | 0,24857909 | 4,0719677 | 0,0000466 | 0,00054938 |
| AC007952.4 | 77,532975 | 1,16366725 | 0,29066496 | 4,06902588 | 0,0000472 | 0,00055426 |
| DCTPP1 | 2597,99471 | 1,11717813 | 0,27411709 | 4,0667583 | 0,0000477 | 0,00055819 |
| WARS1 | 28550,4623 | 1,08044355 | 0,26992507 | 4,0640571 | 0,0000482 | 0,00056313 |
| BRI3BP | 1335,34543 | 1,04957051 | 0,25882536 | 4,04814242 | 0,0000516 | 0,00059269 |
| CDC6 | 1410,79264 | 1,63800346 | 0,40051232 | 4,04157944 | 0,0000531 | 0,00060609 |
| HMGA1 | 1798,81596 | 1,24562942 | 0,30887844 | 4,02957083 | 0,0000559 | 0,00062962 |
| RFC3 | 814,51005 | 1,30070738 | 0,32074068 | 4,02572616 | 0,0000568 | 0,00063763 |
| CKS1B | 1636,82627 | 1,22029692 | 0,30256448 | 4,02273939 | 0,0000575 | 0,00064432 |
| HELLS | 911,07159 | 1,542782 | 0,38126874 | 4,01564698 | 0,0000593 | 0,00066022 |
| TCAM1P | 25,8600269 | 1,02260311 | 0,47239221 | 4,01360092 | 0,0000598 | 0,00066435 |
| EDA2R | 20,8387114 | 1,82530656 | 0,43498774 | 4,00318835 | 0,0000625 | 0,00069177 |
| HMGB2 | 8675,27654 | 1,00558925 | 0,251096 | 4,00110979 | 0,000063 | 0,00069702 |
| CDT1 | 1368,90856 | 1,61856181 | 0,40331384 | 3,97889661 | 0,0000692 | 0,00075425 |
| FZD5 | 41,668731 | 1,30012035 | 0,32147065 | 3,97882493 | 0,0000693 | 0,00075425 |
| CENPU | 902,785896 | 1,51007305 | 0,38358497 | 3,96187243 | 0,0000744 | 0,00080079 |
| RAD51AP1 | 490,051196 | 1,42774933 | 0,35682273 | 3,9615959 | 0,0000745 | 0,00080124 |
| PAQR4 | 1091,18271 | 1,35723293 | 0,34178392 | 3,95995024 | 0,000075 | 0,00080489 |
| HDC | 30,8090943 | 2,33219202 | 0,48736777 | 3,9581798 | 0,0000755 | 0,0008096 |
| POLR3G | 120,074947 | 1,3316655 | 0,33481957 | 3,95740384 | 0,0000758 | 0,00081112 |
| XRCC2 | 370,305949 | 1,40619326 | 0,35690285 | 3,94694686 | 0,0000792 | 0,0008401 |
| UBE2T | 901,685734 | 1,49786465 | 0,37668238 | 3,94064866 | 0,0000813 | 0,00085717 |
| AC011511.5 | 26,6442249 | 1,29618176 | 0,33396073 | 3,94086894 | 0,0000812 | 0,00085717 |
| DONSON | 949,085575 | 1,09582141 | 0,27649212 | 3,94055698 | 0,0000813 | 0,00085717 |

| | | | | | | |
|---|---|---|---|---|---|---|
| CDKN1A | 2383,78661 | 1,5829976 | 0,40776128 | 3,94030883 | 0,0000814 | 0,00085756 |
| MTFR2 | 320,179139 | 1,39504069 | 0,35070288 | 3,93936383 | 0,0000817 | 0,00085995 |
| ELFN1-AS1 | 19,3850066 | 1,67751136 | 0,42667302 | 3,92620031 | 0,0000863 | 0,00089647 |
| PHGDH | 1154,77976 | 1,46015007 | 0,36922628 | 3,92534315 | 0,0000866 | 0,00089916 |
| THSD8 | 364,425606 | 1,19026425 | 0,30716875 | 3,92487223 | 0,0000868 | 0,0009004 |
| STIL | 666,799496 | 1,43452061 | 0,36383328 | 3,92234751 | 0,0000877 | 0,00090731 |
| ATF3 | 271,145157 | 1,08657261 | 0,28875395 | 3,91571619 | 0,0000901 | 0,00092945 |
| ERCC6L | 216,289004 | 1,44576283 | 0,36810749 | 3,91545896 | 0,0000902 | 0,00092992 |
| TTK | 499,963879 | 1,57513627 | 0,39649228 | 3,91459441 | 0,0000906 | 0,00093175 |
| SLC35G1 | 79,1619926 | 1,16368713 | 0,29752546 | 3,91427716 | 0,0000907 | 0,00093185 |
| TIPIN | 806,5352 | 1,15405363 | 0,29753383 | 3,89998518 | 0,0000962 | 0,00097923 |
| RAB3D | 1128,09782 | 1,01239501 | 0,26084934 | 3,89714692 | 0,0000973 | 0,00098867 |
| HSF5 | 107,39634 | 1,25325507 | 0,32196466 | 3,89543348 | 0,000098 | 0,00099502 |

**Supplementary table 2.** List of $ST2^{-}$ $CD56^{bright}$ NK cell signature genes used in this study, Related to Fig. 3B, 3E, 3H, 3I, 5A, 5B, Supplementary Fig. S4E, S9A

| gene | baseMean | log2FoldChange | lfcSE | stat | pvalue | padj |
|---|---|---|---|---|---|---|
| MARCKSL1 | 2140,369762 | 1,740882999 | 0,116707873 | 14,91550027 | 2,61E-50 | 4,39E-47 |
| SGSM3 | 8319,280225 | 1,301010684 | 0,099219228 | 13,1115542 | 2,83E-39 | 2,71E-36 |
| TMEM123 | 8464,060835 | 1,625281824 | 0,129219167 | 12,58132481 | 2,68E-36 | 2,16E-33 |
| LRRC75B | 456,6176087 | 2,416210354 | 0,198847881 | 12,15071388 | 5,69E-34 | 3,70E-31 |
| EVA1B | 263,1092393 | 2,229323471 | 0,188111393 | 11,85127264 | 2,12E-32 | 1,16E-29 |
| IL4R | 3450,852494 | 1,499947728 | 0,126745269 | 11,84412664 | 2,31E-32 | 1,22E-29 |
| CYSLTR1 | 649,5300538 | 1,515430844 | 0,129159975 | 11,73268544 | 8,67E-32 | 4,26E-29 |
| MAN1C1 | 565,7106081 | 3,237435298 | 0,286233564 | 11,29381958 | 1,41E-29 | 6,17E-27 |
| ANKRD13C | 2134,558285 | 1,002724262 | 0,089928491 | 11,15065986 | 7,11E-29 | 2,87E-26 |
| P2RX4 | 2152,637806 | 1,272120915 | 0,115924558 | 10,97495534 | 5,04E-28 | 1,92E-25 |
| CD82 | 977,0473825 | 2,847624791 | 0,264383983 | 10,78862923 | 3,90E-27 | 1,38E-24 |
| PPARD | 1041,824462 | 1,179517604 | 0,111025009 | 10,62431354 | 2,30E-26 | 7,85E-24 |
| CD44 | 17644,69997 | 2,466187162 | 0,235781727 | 10,46120997 | 1,30E-25 | 4,37E-23 |
| CACNB1 | 672,7485259 | 1,620194479 | 0,155622145 | 10,41380983 | 2,14E-25 | 7,09E-23 |
| AKTIP | 1505,369451 | 1,445792576 | 0,139440905 | 10,37395255 | 3,26E-25 | 1,06E-22 |
| FUCA1 | 2045,337861 | 1,842417316 | 0,17993438 | 10,23962265 | 1,32E-24 | 3,97E-22 |
| LINC00996 | 319,627706 | 2,609455123 | 0,256710702 | 10,15786989 | 3,06E-24 | 8,56E-22 |
| FXYD5 | 4272,477726 | 1,196803035 | 0,121577711 | 9,843066925 | 7,34E-23 | 1,85E-20 |
| AFAP1L2 | 218,6357125 | 3,082245531 | 0,312139865 | 9,810966072 | 1,01E-22 | 2,51E-20 |
| RAB38 | 243,3177113 | 3,055826055 | 0,313133124 | 9,801295845 | 1,11E-22 | 2,70E-20 |
| AHI1 | 1559,477129 | 2,477023515 | 0,255683885 | 9,685122861 | 3,49E-22 | 7,81E-20 |
| SOX4 | 196,001897 | 2,237665613 | 0,23699926 | 9,444790764 | 3,56E-21 | 7,25E-19 |
| KIAA1211L | 88,75908207 | 2,71897675 | 0,294775003 | 9,213852772 | 3,15E-20 | 5,97E-18 |
| IGFLR1 | 3854,076961 | 1,143615923 | 0,124148381 | 9,209966716 | 3,26E-20 | 6,09E-18 |

| | | | | | | |
|---|---|---|---|---|---|---|
| TLE3 | 1567,104006 | 2,590685551 | 0,282744437 | 9,166413642 | 4,89E-20 | 8,80E-18 |
| RARA | 600,9572647 | 1,318907813 | 0,145484053 | 9,062144833 | 1,28E-19 | 2,22E-17 |
| STBD1 | 219,2555171 | 1,822615346 | 0,209614914 | 8,717454048 | 2,85E-18 | 4,41E-16 |
| FUT8-AS1 | 67,90486061 | 2,855822847 | 0,328357416 | 8,695225091 | 3,46E-18 | 5,29E-16 |
| MMP25 | 1398,742723 | 1,40354112 | 0,162428028 | 8,639506398 | 5,65E-18 | 8,43E-16 |
| FAM102A | 2599,12843 | 1,914468847 | 0,222371703 | 8,607765357 | 7,45E-18 | 1,09E-15 |
| AL034397.3 | 264,4978444 | 2,197245925 | 0,256021938 | 8,601655494 | 7,86E-18 | 1,14E-15 |
| TMEM200A | 142,3148917 | 2,89847412 | 0,337566616 | 8,571643818 | 1,02E-17 | 1,47E-15 |
| KIR2DL4 | 5176,736346 | 3,205020743 | 0,372271606 | 8,56145511 | 1,11E-17 | 1,56E-15 |
| SPTBN1 | 5050,407322 | 1,454201938 | 0,169895179 | 8,562040027 | 1,11E-17 | 1,56E-15 |
| GRAMD4 | 1739,584588 | 1,256608365 | 0,148997408 | 8,426004572 | 3,58E-17 | 4,74E-15 |
| AXIN2 | 122,846397 | 4,198344719 | 0,496186938 | 8,424218323 | 3,63E-17 | 4,79E-15 |
| TRAF5 | 5084,644085 | 1,264733035 | 0,151119937 | 8,371710643 | 5,68E-17 | 7,29E-15 |
| AC136475.3 | 181,4399858 | 3,322393183 | 0,393086018 | 8,331814546 | 7,96E-17 | 1,00E-14 |
| AC008691.1 | 15,3365338 | 3,974901787 | 0,471562234 | 8,271214677 | 1,33E-16 | 1,61E-14 |
| PRAM1 | 324,8694184 | 3,248314446 | 0,39599038 | 8,151417564 | 3,60E-16 | 4,12E-14 |
| FOXC1 | 42,81840529 | 4,257531481 | 0,513555488 | 8,146888353 | 3,73E-16 | 4,25E-14 |
| FUT8 | 857,1317759 | 2,256320072 | 0,277545555 | 8,121125486 | 4,62E-16 | 5,20E-14 |
| AREG | 49,15236748 | 2,404981214 | 0,298520266 | 8,0837208 | 6,28E-16 | 7,04E-14 |
| ANKRD6 | 380,9412248 | 1,408207328 | 0,176085911 | 8,082307646 | 6,36E-16 | 7,08E-14 |
| HVCN1 | 1192,528257 | 2,419885321 | 0,301033606 | 8,025608906 | 1,01E-15 | 1,09E-13 |
| MPZL3 | 2867,960136 | 1,96187017 | 0,244903273 | 8,005583963 | 1,19E-15 | 1,27E-13 |
| LRFN1 | 903,3538527 | 1,46235046 | 0,183524361 | 7,969598699 | 1,59E-15 | 1,66E-13 |
| BACH2 | 1055,397026 | 2,043278573 | 0,256745168 | 7,956655052 | 1,77E-15 | 1,83E-13 |
| IFITM10 | 79,33410328 | 2,819037525 | 0,358126867 | 7,908149319 | 2,61E-15 | 2,67E-13 |
| LMNA | 764,4263779 | 3,588425802 | 0,460469059 | 7,795625921 | 6,41E-15 | 6,37E-13 |
| MAML2 | 405,3947041 | 1,969330901 | 0,254612167 | 7,733434442 | 1,05E-14 | 1,01E-12 |
| TIAM1 | 2217,418454 | 3,446940657 | 0,443312808 | 7,712998259 | 1,23E-14 | 1,18E-12 |

| | | | | | | |
|---|---|---|---|---|---|---|
| LDLRAD2 | 37,78263563 | 2,864315899 | 0,37090984 | 7,706069704 | 1,30E-14 | 1,23E-12 |
| PRCD | 141,6670188 | 2,886854421 | 0,384456276 | 7,657090385 | 1,90E-14 | 1,76E-12 |
| KLHL6 | 2736,886324 | 1,449384166 | 0,189438168 | 7,656948369 | 1,90E-14 | 1,76E-12 |
| ZFHX3 | 162,5904485 | 2,099702068 | 0,274363937 | 7,648660179 | 2,03E-14 | 1,87E-12 |
| FLNB | 1219,316738 | 2,293855562 | 0,300509328 | 7,632018925 | 2,31E-14 | 2,10E-12 |
| C19orf38 | 314,3637498 | 2,865738545 | 0,381649932 | 7,60950385 | 2,75E-14 | 2,46E-12 |
| TEC | 310,8560866 | 2,442201965 | 0,320801105 | 7,608939573 | 2,76E-14 | 2,46E-12 |
| DOK7 | 143,5755317 | 2,752814433 | 0,364946495 | 7,596460325 | 3,04E-14 | 2,68E-12 |
| LDOC1 | 547,4136479 | 1,203333137 | 0,159103248 | 7,587139728 | 3,27E-14 | 2,85E-12 |
| IER5L | 207,4254029 | 3,928176017 | 0,512131998 | 7,561144111 | 4,00E-14 | 3,43E-12 |
| STARD10 | 965,4029696 | 1,517324896 | 0,202110986 | 7,524072375 | 5,31E-14 | 4,52E-12 |
| CAPN12 | 1574,643552 | 1,986846274 | 0,264452466 | 7,514838013 | 5,70E-14 | 4,81E-12 |
| RRM2P3 | 33,88671537 | 1,906501022 | 0,255080085 | 7,514772312 | 5,70E-14 | 4,81E-12 |
| PLK2 | 98,37650677 | 2,344139862 | 0,312635658 | 7,475686672 | 7,68E-14 | 6,37E-12 |
| PRPF40B | 405,9369242 | 1,255426299 | 0,168137217 | 7,466021351 | 8,27E-14 | 6,83E-12 |
| IL7R | 1849,842269 | 3,752113699 | 0,497696008 | 7,41606311 | 1,21E-13 | 9,85E-12 |
| FAAH | 346,5778786 | 1,680102357 | 0,22747979 | 7,382156404 | 1,56E-13 | 1,24E-11 |
| TNFSF9 | 132,6527701 | 3,637122844 | 0,475030088 | 7,330987379 | 2,28E-13 | 1,78E-11 |
| MBOAT7 | 1550,633095 | 1,332621699 | 0,182816461 | 7,288449341 | 3,14E-13 | 2,39E-11 |
| DST | 209,6756475 | 1,816584497 | 0,250223788 | 7,282464592 | 3,28E-13 | 2,48E-11 |
| ZBTB46 | 128,1886131 | 3,641740392 | 0,496162257 | 7,228772817 | 4,87E-13 | 3,61E-11 |
| MAFF | 2257,925472 | 1,879053686 | 0,260093808 | 7,224527248 | 5,03E-13 | 3,70E-11 |
| ZNF217 | 1450,119102 | 1,287753617 | 0,178824246 | 7,201997933 | 5,93E-13 | 4,32E-11 |
| UNC93B1 | 3043,964328 | 1,750329949 | 0,243608534 | 7,184379072 | 6,75E-13 | 4,90E-11 |
| TPCN1 | 1699,033407 | 1,53151534 | 0,213456563 | 7,17768457 | 7,09E-13 | 5,11E-11 |
| ZNF296 | 727,0590777 | 1,305088768 | 0,182369891 | 7,156348565 | 8,29E-13 | 5,90E-11 |
| PLEKHN1 | 206,4025607 | 2,085181376 | 0,292403136 | 7,122297134 | 1,06E-12 | 7,46E-11 |
| SMG6 | 1585,486005 | 1,057027036 | 0,148854315 | 7,10138665 | 1,24E-12 | 8,59E-11 |

| | | | | | | |
|---|---|---|---|---|---|---|
| XYLT1 | 1465,872637 | 1,156545617 | 0,163384884 | 7,077618012 | 1,47E-12 | 1,01E-10 |
| SLC39A10 | 1623,403389 | 1,212661365 | 0,173634103 | 6,988704433 | 2,77E-12 | 1,87E-10 |
| HID1 | 39,66821395 | 2,205264018 | 0,320151044 | 6,966474531 | 3,25E-12 | 2,16E-10 |
| RRAD | 58,48146211 | 2,953436907 | 0,428687381 | 6,90543153 | 5,01E-12 | 3,28E-10 |
| MPG | 2611,401069 | 1,003775495 | 0,145291907 | 6,904810892 | 5,03E-12 | 3,28E-10 |
| FBXL16 | 35,19675056 | 3,730713076 | 0,511691174 | 6,896414366 | 5,33E-12 | 3,47E-10 |
| PLPP1 | 233,9239877 | 1,219327306 | 0,178811692 | 6,843240224 | 7,74E-12 | 4,92E-10 |
| PLXNA4 | 961,9891073 | 2,158678049 | 0,318267672 | 6,83276983 | 8,33E-12 | 5,26E-10 |
| TBXT | 24,26215193 | 5,06557221 | 0,664143384 | 6,814910581 | 9,43E-12 | 5,87E-10 |
| METTL24 | 29,66113697 | 2,986686143 | 0,454225853 | 6,799242063 | 1,05E-11 | 6,45E-10 |
| GPR68 | 3491,357851 | 1,549838753 | 0,22953245 | 6,781940691 | 1,19E-11 | 7,20E-10 |
| SNX30 | 333,4991655 | 1,840690351 | 0,272887511 | 6,764333072 | 1,34E-11 | 8,09E-10 |
| PDE6G | 738,1239368 | 1,855810087 | 0,275709691 | 6,751372835 | 1,46E-11 | 8,76E-10 |
| AL158211.5 | 55,05019248 | 3,18301925 | 0,471540867 | 6,699949582 | 2,08E-11 | 1,22E-09 |
| THEM4 | 1339,204355 | 1,240865036 | 0,187320759 | 6,627140101 | 3,42E-11 | 1,94E-09 |
| GATA6 | 148,983322 | 3,806964818 | 0,557597993 | 6,618724163 | 3,62E-11 | 2,05E-09 |
| SPIN3 | 579,5682663 | 2,150287743 | 0,325929848 | 6,604060126 | 4,00E-11 | 2,23E-09 |
| GOLGA6L10 | 65,06390646 | 1,759747802 | 0,268302629 | 6,598321389 | 4,16E-11 | 2,31E-09 |
| GRAMD1B | 353,2229995 | 2,83284927 | 0,428341998 | 6,586356155 | 4,51E-11 | 2,49E-09 |
| CD300C | 662,9534538 | 2,141666401 | 0,329522641 | 6,574886891 | 4,87E-11 | 2,67E-09 |
| S100A13 | 51,69556697 | 1,57700331 | 0,241041752 | 6,54188058 | 6,08E-11 | 3,27E-09 |
| EXT1 | 752,9336282 | 1,058061661 | 0,162404986 | 6,516422519 | 7,20E-11 | 3,81E-09 |
| SH3D21 | 63,91059322 | 2,212082091 | 0,339848211 | 6,508575435 | 7,59E-11 | 4,00E-09 |
| ERCC6 | 1325,076914 | 1,196918771 | 0,184243302 | 6,498806178 | 8,10E-11 | 4,24E-09 |
| SUOX | 514,9009048 | 1,922142601 | 0,295842521 | 6,492744365 | 8,43E-11 | 4,39E-09 |
| IRF2BPL | 951,7592032 | 1,293991659 | 0,199419253 | 6,490984681 | 8,53E-11 | 4,43E-09 |
| AL157935.2 | 12,34158825 | 2,475195319 | 0,383015829 | 6,451818604 | 1,11E-10 | 5,67E-09 |
| GPR35 | 331,9485478 | 1,26233446 | 0,196491366 | 6,414399391 | 1,41E-10 | 7,11E-09 |

| | | | | | | |
|---|---|---|---|---|---|---|
| ERMP1 | 6432,990792 | 1,050931817 | 0,164670882 | 6,400523906 | 1,55E-10 | 7,69E-09 |
| PPP1R9A | 493,7632895 | 1,981004521 | 0,309699355 | 6,391466414 | 1,64E-10 | 8,14E-09 |
| TAF9B | 820,1818757 | 1,285949427 | 0,201407611 | 6,384789994 | 1,72E-10 | 8,42E-09 |
| LRP5 | 94,50272901 | 3,172582403 | 0,489045222 | 6,380459222 | 1,77E-10 | 8,61E-09 |
| ITGAX | 12415,82202 | 2,079106969 | 0,329943746 | 6,35380944 | 2,10E-10 | 1,01E-08 |
| KLRC2 | 726,1948602 | 1,217769369 | 0,191269433 | 6,347808523 | 2,18E-10 | 1,04E-08 |
| ZNF629 | 237,397175 | 1,314223877 | 0,207922122 | 6,339709292 | 2,30E-10 | 1,09E-08 |
| CXCR3 | 3176,952781 | 2,727739219 | 0,432006382 | 6,289496674 | 3,18E-10 | 1,47E-08 |
| DND1P1 | 139,7377777 | 1,591147032 | 0,253169883 | 6,281714343 | 3,35E-10 | 1,54E-08 |
| RNASE6 | 49,3304587 | 1,8336818 | 0,294251275 | 6,241170135 | 4,34E-10 | 1,95E-08 |
| LINC02273 | 120,6854075 | 2,104712474 | 0,334746798 | 6,237198999 | 4,45E-10 | 2,00E-08 |
| NOD2 | 627,9389415 | 1,373877227 | 0,221025473 | 6,236011813 | 4,49E-10 | 2,01E-08 |
| ZEB1 | 804,8908063 | 1,574559748 | 0,252463737 | 6,234338719 | 4,54E-10 | 2,02E-08 |
| LINC00271 | 60,35176693 | 2,032485041 | 0,326992021 | 6,22232025 | 4,90E-10 | 2,18E-08 |
| COL4A4 | 136,6893954 | 2,877693674 | 0,464567299 | 6,190854914 | 5,98E-10 | 2,60E-08 |
| RHOB | 142,5630848 | 1,646035215 | 0,266532796 | 6,153458318 | 7,58E-10 | 3,23E-08 |
| WNT11 | 357,6166372 | 2,750818924 | 0,447235458 | 6,140192481 | 8,24E-10 | 3,48E-08 |
| ITM2C | 313,5989235 | 3,078454925 | 0,495274515 | 6,138832445 | 8,31E-10 | 3,51E-08 |
| PATJ | 832,4500235 | 2,026383599 | 0,331901279 | 6,116181619 | 9,58E-10 | 3,98E-08 |
| CLSTN3 | 2662,810035 | 1,451565636 | 0,237248483 | 6,116238957 | 9,58E-10 | 3,98E-08 |
| SUPT3H | 215,3559279 | 1,029318783 | 0,169361342 | 6,078184656 | 1,22E-09 | 4,95E-08 |
| TP53I11 | 303,9506515 | 1,927634969 | 0,330736618 | 6,072075791 | 1,26E-09 | 5,13E-08 |
| ITM2A | 2462,385495 | 1,136864047 | 0,187341643 | 6,070902966 | 1,27E-09 | 5,16E-08 |
| PI16 | 42,04003912 | 4,176725001 | 0,65065832 | 6,049442295 | 1,45E-09 | 5,84E-08 |
| BEX4 | 1500,538493 | 1,086656664 | 0,180329332 | 6,028447297 | 1,66E-09 | 6,56E-08 |
| RGS10 | 579,2833941 | 1,532265566 | 0,254414155 | 6,020657274 | 1,74E-09 | 6,81E-08 |
| AC002383.1 | 10,8397303 | 4,32815078 | 0,680315877 | 6,015521954 | 1,79E-09 | 7,00E-08 |
| CD55 | 3052,146888 | 1,09862975 | 0,18266267 | 6,015323622 | 1,80E-09 | 7,00E-08 |

| | | | | | | |
|---|---|---|---|---|---|---|
| BEX2 | 385,7628247 | 1,249224469 | 0,208923302 | 6,002445603 | 1,94E-09 | 7,52E-08 |
| XIRP1 | 57,22071206 | 3,683841193 | 0,573027442 | 5,982503354 | 2,20E-09 | 8,41E-08 |
| CDHR1 | 819,9741477 | 2,677331521 | 0,444707403 | 5,959096751 | 2,54E-09 | 9,51E-08 |
| CRYBG3 | 125,5555099 | 2,063911456 | 0,35247776 | 5,942522032 | 2,81E-09 | 1,05E-07 |
| RAB43 | 2036,021922 | 1,001743046 | 0,168646318 | 5,941980609 | 2,82E-09 | 1,05E-07 |
| TMEM38A | 82,82198594 | 3,210889069 | 0,538726068 | 5,937159296 | 2,90E-09 | 1,07E-07 |
| ARMC5 | 1780,369751 | 1,389518757 | 0,234393229 | 5,927321137 | 3,08E-09 | 1,14E-07 |
| CLEC12A-AS1 | 46,22362818 | 1,931710213 | 0,330869574 | 5,923534282 | 3,15E-09 | 1,16E-07 |
| RELL1 | 724,4778869 | 1,283650735 | 0,216901888 | 5,921056996 | 3,20E-09 | 1,17E-07 |
| ACVR2A | 307,8483385 | 1,435989021 | 0,243605706 | 5,915557483 | 3,31E-09 | 1,21E-07 |
| RCBTB2 | 3063,875493 | 2,182264742 | 0,368944091 | 5,907303203 | 3,48E-09 | 1,27E-07 |
| AL022322.2 | 67,76634459 | 1,801975645 | 0,307307468 | 5,888891436 | 3,89E-09 | 1,41E-07 |
| SCARF1 | 156,4142594 | 2,842376226 | 0,482660895 | 5,876623721 | 4,19E-09 | 1,50E-07 |
| BGLAP | 96,04635907 | 3,538994106 | 0,589813919 | 5,862238586 | 4,57E-09 | 1,63E-07 |
| SV2A | 48,2437807 | 2,445816786 | 0,44854443 | 5,852427352 | 4,84E-09 | 1,71E-07 |
| SMIM10L2A | 64,30658026 | 3,508776996 | 0,594553252 | 5,837677632 | 5,29E-09 | 1,87E-07 |
| PLEC | 2056,099146 | 1,861878277 | 0,320559179 | 5,836288724 | 5,34E-09 | 1,88E-07 |
| FXYD7 | 38,81660948 | 3,469733075 | 0,578446437 | 5,830312225 | 5,53E-09 | 1,94E-07 |
| RAB3GAP1 | 663,7483876 | 1,089492619 | 0,187676911 | 5,809340557 | 6,27E-09 | 2,17E-07 |
| EXD2 | 489,2651708 | 1,22891323 | 0,21183615 | 5,806800475 | 6,37E-09 | 2,20E-07 |
| PRDM8 | 497,0459425 | 1,594431879 | 0,274486306 | 5,790720871 | 7,01E-09 | 2,41E-07 |
| AL357033.4 | 101,5975193 | 1,590155519 | 0,27616053 | 5,765928866 | 8,12E-09 | 2,74E-07 |
| LTC4S | 42,46148258 | 2,210105354 | 0,382728867 | 5,760233058 | 8,40E-09 | 2,82E-07 |
| COL9A2 | 666,9338283 | 2,977350103 | 0,517402243 | 5,751054742 | 8,87E-09 | 2,96E-07 |
| AL031736.1 | 103,687593 | 1,950697715 | 0,33843021 | 5,742233571 | 9,34E-09 | 3,10E-07 |
| GAB1 | 175,9057184 | 2,737081522 | 0,477591498 | 5,708355806 | 1,14E-08 | 3,73E-07 |
| SSBP2 | 719,3021611 | 1,973183258 | 0,345798583 | 5,705182996 | 1,16E-08 | 3,79E-07 |
| DEPDC7 | 126,9076612 | 2,000178345 | 0,352564952 | 5,698178526 | 1,21E-08 | 3,93E-07 |

| | | | | | | |
|---|---|---|---|---|---|---|
| AHR | 1030,419847 | 2,406975171 | 0,421207467 | 5,693457474 | 1,24E-08 | 4,01E-07 |
| PLCH2 | 1656,044125 | 1,331782742 | 0,23544833 | 5,659271744 | 1,52E-08 | 4,83E-07 |
| AL132996.1 | 6,930815148 | 3,865822551 | 0,658816638 | 5,655974876 | 1,55E-08 | 4,91E-07 |
| CRTAM | 3258,918725 | 2,879944593 | 0,498895482 | 5,635140157 | 1,75E-08 | 5,49E-07 |
| EPAS1 | 235,0594084 | 1,780244369 | 0,315446327 | 5,624532859 | 1,86E-08 | 5,82E-07 |
| RPS6KA2 | 603,4370845 | 2,081404584 | 0,375350496 | 5,615442365 | 1,96E-08 | 6,13E-07 |
| AC021066.1 | 445,2279295 | 1,806776785 | 0,322754978 | 5,588460676 | 2,29E-08 | 7,08E-07 |
| SPECC1 | 1212,882897 | 1,008531761 | 0,180580378 | 5,582913532 | 2,37E-08 | 7,28E-07 |
| MTDHP3 | 18,53111258 | 3,651885327 | 0,630130448 | 5,57811123 | 2,43E-08 | 7,47E-07 |
| AL603783.1 | 43,40415723 | 2,919117033 | 0,516839812 | 5,577366312 | 2,44E-08 | 7,48E-07 |
| NBPF15 | 523,6238388 | 1,003156505 | 0,179966039 | 5,577463011 | 2,44E-08 | 7,48E-07 |
| AC073332.1 | 50,46757368 | 2,079539706 | 0,374584686 | 5,568107727 | 2,58E-08 | 7,88E-07 |
| AC011445.1 | 149,0232011 | 1,448193495 | 0,261854134 | 5,539578792 | 3,03E-08 | 9,14E-07 |
| TCF7 | 3711,66176 | 2,801294186 | 0,501319966 | 5,532091697 | 3,16E-08 | 9,52E-07 |
| AL022341.2 | 64,00754303 | 2,406238912 | 0,436859003 | 5,525605382 | 3,28E-08 | 9,82E-07 |
| NELL2 | 528,3413071 | 2,958904755 | 0,536983549 | 5,519567819 | 3,40E-08 | 1,01E-06 |
| GPR155 | 584,2908913 | 1,362453331 | 0,246781698 | 5,518453369 | 3,42E-08 | 1,02E-06 |
| GORASP1 | 727,874936 | 1,159369536 | 0,212809321 | 5,517578996 | 3,44E-08 | 1,02E-06 |
| MAML3 | 227,3004088 | 2,910517687 | 0,520119301 | 5,507021751 | 3,65E-08 | 1,07E-06 |
| NCKAP5L | 557,8377929 | 1,019341234 | 0,185736999 | 5,495749561 | 3,89E-08 | 1,14E-06 |
| NBPF9 | 286,8213266 | 1,104871219 | 0,201721668 | 5,487263097 | 4,08E-08 | 1,19E-06 |
| AC068580.4 | 1237,365465 | 1,09636765 | 0,199802887 | 5,481917224 | 4,21E-08 | 1,22E-06 |
| PRSS2 | 20,42490575 | 3,435662521 | 0,640356433 | 5,468191576 | 4,55E-08 | 1,31E-06 |
| SEC22B4P | 427,8475782 | 1,068157178 | 0,195364259 | 5,467566083 | 4,56E-08 | 1,32E-06 |
| AC136475.9 | 40,38923353 | 2,683705638 | 0,492099609 | 5,467015518 | 4,58E-08 | 1,32E-06 |
| KLHDC9 | 16,03763148 | 2,554000388 | 0,46130946 | 5,46690161 | 4,58E-08 | 1,32E-06 |
| PLA2G6 | 1633,729643 | 2,025368206 | 0,371533287 | 5,451318684 | 5,00E-08 | 1,42E-06 |
| NUDT16P1 | 132,0159874 | 2,394800836 | 0,440749668 | 5,433913468 | 5,51E-08 | 1,56E-06 |

| | | | | | | |
|---|---|---|---|---|---|---|
| IFITM3 | 7923,417658 | 1,451337304 | 0,267556571 | 5,43083565 | 5,61E-08 | 1,58E-06 |
| MTM1 | 1010,089782 | 1,886289418 | 0,347656721 | 5,420132252 | 5,96E-08 | 1,68E-06 |
| GPR82 | 337,6226796 | 2,635019703 | 0,481344315 | 5,419871453 | 5,96E-08 | 1,68E-06 |
| ATP6V0E2-AS1 | 108,2815423 | 1,244807485 | 0,230542859 | 5,410393293 | 6,29E-08 | 1,76E-06 |
| LINC01871 | 1830,467521 | 1,540975424 | 0,285096179 | 5,408546324 | 6,35E-08 | 1,78E-06 |
| LSR | 887,6666794 | 2,840944598 | 0,518173198 | 5,401460927 | 6,61E-08 | 1,85E-06 |
| AL034550.2 | 33,02488769 | 3,172151524 | 0,559528571 | 5,395879874 | 6,82E-08 | 1,90E-06 |
| AC092368.3 | 24,78169397 | 1,512806742 | 0,283628302 | 5,379991707 | 7,45E-08 | 2,06E-06 |
| TPD52L1 | 51,71867641 | 2,422752867 | 0,455664353 | 5,373551856 | 7,72E-08 | 2,13E-06 |
| CD2 | 9315,600078 | 1,454866474 | 0,272244595 | 5,363995817 | 8,14E-08 | 2,24E-06 |
| PFKFB3 | 1945,504031 | 1,000897819 | 0,187849186 | 5,329454109 | 9,85E-08 | 2,65E-06 |
| LINC02446 | 16,30915541 | 2,489644747 | 0,459313591 | 5,324837854 | 1,01E-07 | 2,72E-06 |
| ANKDD1B | 17,25372673 | 1,915061785 | 0,364217268 | 5,324244425 | 1,01E-07 | 2,72E-06 |
| EPHX1 | 54,71961291 | 2,458322408 | 0,471892105 | 5,292055384 | 1,21E-07 | 3,18E-06 |
| EPHA1 | 89,4450334 | 1,590839766 | 0,303668555 | 5,283878035 | 1,26E-07 | 3,32E-06 |
| PALD1 | 72,54013307 | 2,564431066 | 0,484668735 | 5,283191913 | 1,27E-07 | 3,33E-06 |
| GSN | 11477,76579 | 1,080001767 | 0,204600157 | 5,282113632 | 1,28E-07 | 3,34E-06 |
| SLC17A9 | 1020,722123 | 2,025156259 | 0,383167497 | 5,274721728 | 1,33E-07 | 3,48E-06 |
| ZNF667 | 63,99101562 | 3,275472613 | 0,616486896 | 5,226061181 | 1,73E-07 | 4,42E-06 |
| SPACA9 | 113,6159267 | 2,142499804 | 0,408667207 | 5,222418166 | 1,77E-07 | 4,50E-06 |
| CSF2 | 167,7107889 | 3,291839501 | 0,596805233 | 5,214685992 | 1,84E-07 | 4,67E-06 |
| ZNF471 | 57,49951783 | 2,226779851 | 0,434540854 | 5,214527878 | 1,84E-07 | 4,67E-06 |
| ARMCX2 | 60,84843318 | 1,884167269 | 0,360777333 | 5,214744992 | 1,84E-07 | 4,67E-06 |
| EGR3 | 43,79934598 | 2,259073472 | 0,432640676 | 5,206379511 | 1,93E-07 | 4,85E-06 |
| TTC23 | 29,86479356 | 1,512348688 | 0,295236661 | 5,206164654 | 1,93E-07 | 4,85E-06 |
| BEND5 | 33,96012583 | 2,096798976 | 0,408007954 | 5,199110676 | 2,00E-07 | 5,03E-06 |
| TRPM8 | 20,34419262 | 3,04250246 | 0,678290192 | 5,195196907 | 2,05E-07 | 5,12E-06 |

| | | | | | | |
|---|---|---|---|---|---|---|
| WDR86-AS1 | 5,171232188 | 3,921026082 | 0,684325649 | 5,193465345 | 2,06E-07 | 5,16E-06 |
| SLC49A3 | 485,7237538 | 1,312431718 | 0,253372909 | 5,184921568 | 2,16E-07 | 5,39E-06 |
| TUBA8 | 95,46203126 | 2,550813663 | 0,490022933 | 5,170630572 | 2,33E-07 | 5,79E-06 |
| RNF144A | 1772,527623 | 1,289472324 | 0,250586361 | 5,162317942 | 2,44E-07 | 6,03E-06 |
| SPRY2 | 473,2348813 | 2,107772018 | 0,408888309 | 5,147224675 | 2,64E-07 | 6,51E-06 |
| DDIT4 | 5551,55676 | 1,362385866 | 0,26530604 | 5,144237245 | 2,69E-07 | 6,61E-06 |
| SLC44A1 | 1742,706472 | 1,401151872 | 0,273392571 | 5,125205954 | 2,97E-07 | 7,23E-06 |
| ASPH | 249,8983469 | 1,797842619 | 0,35231292 | 5,105993649 | 3,29E-07 | 7,92E-06 |
| AC009404.1 | 43,90260786 | 1,29603386 | 0,254206162 | 5,101933528 | 3,36E-07 | 8,08E-06 |
| ENPP3 | 119,5988475 | 2,667883964 | 0,57024697 | 5,093649621 | 3,51E-07 | 8,39E-06 |
| CD27 | 111,4046967 | 3,032675704 | 0,579396161 | 5,06209634 | 4,15E-07 | 9,84E-06 |
| CPNE7 | 81,84795563 | 1,838326943 | 0,363674022 | 5,0473816 | 4,48E-07 | 1,05E-05 |
| LINC02391 | 9,538284846 | 2,131988111 | 0,422294001 | 5,046154098 | 4,51E-07 | 1,06E-05 |
| MFGE8 | 399,3689337 | 1,248535075 | 0,247873273 | 5,045848157 | 4,52E-07 | 1,06E-05 |
| AC004687.3 | 18,53533533 | 3,779035129 | 0,68637623 | 5,04379158 | 4,56E-07 | 1,07E-05 |
| SLC29A4 | 26,32821855 | 2,370306808 | 0,468206092 | 5,043496764 | 4,57E-07 | 1,07E-05 |
| TRO | 241,9601661 | 1,037435872 | 0,207147565 | 5,021933504 | 5,12E-07 | 1,19E-05 |
| AC011298.1 | 12,69198312 | 3,717939806 | 0,684965986 | 5,015416738 | 5,29E-07 | 1,22E-05 |
| AC136475.5 | 16,14527145 | 2,820416285 | 0,563897138 | 5,015266466 | 5,30E-07 | 1,22E-05 |
| COX6A2 | 11,17509384 | 3,840578061 | 0,686505729 | 4,995326291 | 5,87E-07 | 1,35E-05 |
| AC097658.2 | 6,571846465 | 2,389027138 | 0,470204318 | 4,983528216 | 6,24E-07 | 1,42E-05 |
| PLCB4 | 8,966236706 | 3,114548233 | 0,662253761 | 4,977668738 | 6,44E-07 | 1,46E-05 |
| FXYD2 | 28,56930801 | 3,374607679 | 0,679677749 | 4,973572964 | 6,57E-07 | 1,49E-05 |
| MS4A1 | 103,7166229 | 2,651249066 | 0,544912467 | 4,960162463 | 7,04E-07 | 1,59E-05 |
| SRGAP3 | 102,6996534 | 2,09325268 | 0,424847963 | 4,953356723 | 7,29E-07 | 1,64E-05 |
| IL9RP3 | 32,18469003 | 2,965586928 | 0,614113692 | 4,945474988 | 7,60E-07 | 1,70E-05 |
| EGR2 | 775,1497977 | 1,551361391 | 0,312263 | 4,943510388 | 7,67E-07 | 1,72E-05 |
| CASS4 | 376,6299138 | 1,107652538 | 0,224036719 | 4,943290919 | 7,68E-07 | 1,72E-05 |

| | | | | | | |
|---|---|---|---|---|---|---|
| CAV1 | 12,3243057 | 3,000225407 | 0,669579558 | 4,936992385 | 7,93E-07 | 1,77E-05 |
| SEPTIN11 | 3368,522987 | 1,230166792 | 0,249533059 | 4,93445316 | 8,04E-07 | 1,79E-05 |
| DAPP1 | 453,4751878 | 1,810976987 | 0,378679342 | 4,933798851 | 8,06E-07 | 1,79E-05 |
| KCTD9 | 622,9600217 | 1,554305672 | 0,315884338 | 4,932125087 | 8,13E-07 | 1,81E-05 |
| NR4A3 | 60,41180688 | 2,390151939 | 0,482092819 | 4,931824622 | 8,15E-07 | 1,81E-05 |
| AL390719.1 | 21,22682362 | 2,173717679 | 0,438728664 | 4,924569418 | 8,45E-07 | 1,87E-05 |
| AC093890.1 | 84,29101585 | 1,850806298 | 0,37780216 | 4,919792993 | 8,66E-07 | 1,92E-05 |
| LINC01862 | 165,8059183 | 3,407342196 | 0,647630517 | 4,918208601 | 8,73E-07 | 1,93E-05 |
| NPTX2 | 11,95920836 | 2,674611464 | 0,647295525 | 4,903056914 | 9,44E-07 | 2,07E-05 |
| AC246785.2 | 54,40084889 | 1,103043325 | 0,224853183 | 4,902309554 | 9,47E-07 | 2,07E-05 |
| GNG7 | 32,13907297 | 2,162926917 | 0,455478684 | 4,901646418 | 9,50E-07 | 2,08E-05 |
| FHL1 | 356,3105433 | 1,851953834 | 0,378469867 | 4,898587642 | 9,65E-07 | 2,11E-05 |
| TNFSF11 | 290,4070602 | 2,662973414 | 0,535577669 | 4,898267525 | 9,67E-07 | 2,11E-05 |
| E2F3 | 2431,09033 | 1,106357893 | 0,227053379 | 4,881546688 | 1,05E-06 | 2,27E-05 |
| SERPINE1 | 24,51291118 | 3,069040041 | 0,611586793 | 4,879453656 | 1,06E-06 | 2,29E-05 |
| AL512625.1 | 136,192134 | 1,160967664 | 0,239929247 | 4,87860271 | 1,07E-06 | 2,30E-05 |
| LEF1 | 2008,647706 | 2,61189181 | 0,527151981 | 4,872815174 | 1,10E-06 | 2,36E-05 |
| DCAF4 | 454,5373992 | 1,152808831 | 0,236887987 | 4,871929897 | 1,11E-06 | 2,37E-05 |
| PIK3R6 | 267,3136241 | 2,728241743 | 0,551094203 | 4,869987304 | 1,12E-06 | 2,39E-05 |
| TNFSF10 | 2868,030902 | 1,088892881 | 0,223705676 | 4,865157215 | 1,14E-06 | 2,45E-05 |
| A1BG | 168,3731151 | 1,775440392 | 0,365395825 | 4,855055251 | 1,20E-06 | 2,56E-05 |
| AC016026.1 | 86,36717872 | 1,052261414 | 0,217943588 | 4,848636915 | 1,24E-06 | 2,64E-05 |
| PGBD1 | 81,40353523 | 1,717651033 | 0,362680229 | 4,847121884 | 1,25E-06 | 2,65E-05 |
| MMRN1 | 259,7084239 | 3,018811421 | 0,620565044 | 4,841825249 | 1,29E-06 | 2,72E-05 |
| CCR2 | 145,1278363 | 2,860956803 | 0,574266949 | 4,83652106 | 1,32E-06 | 2,78E-05 |
| RGS9BP | 34,79056254 | 2,361203239 | 0,496212139 | 4,830697487 | 1,36E-06 | 2,86E-05 |
| LGALS9C | 706,5233468 | 1,509910795 | 0,322621982 | 4,828798788 | 1,37E-06 | 2,88E-05 |
| SUCLG2-AS1 | 107,4804422 | 1,589844771 | 0,330428536 | 4,814550363 | 1,48E-06 | 3,07E-05 |

| | | | | | | |
|---|---|---|---|---|---|---|
| PGGHG | 7170,992647 | 1,625079721 | 0,338907522 | 4,784349433 | 1,72E-06 | 3,54E-05 |
| EFHC2 | 306,4969901 | 1,666384281 | 0,349250247 | 4,779806872 | 1,75E-06 | 3,61E-05 |
| GNG4 | 13,38166738 | 3,909794942 | 0,683043142 | 4,773653198 | 1,81E-06 | 3,71E-05 |
| UBE2Q2P2 | 175,2414379 | 1,404562422 | 0,295652145 | 4,76984627 | 1,84E-06 | 3,77E-05 |
| TNFRSF10D | 45,93736542 | 2,132218736 | 0,478037828 | 4,756439585 | 1,97E-06 | 4,02E-05 |
| ZNF667-AS1 | 191,2023662 | 2,490988306 | 0,525674202 | 4,7365302 | 2,17E-06 | 4,38E-05 |
| CUEDC1 | 35,54985212 | 2,292075626 | 0,481057487 | 4,724454283 | 2,31E-06 | 4,63E-05 |
| EGLN3 | 44,92095041 | 2,24550515 | 0,479632739 | 4,711643955 | 2,46E-06 | 4,90E-05 |
| INSC | 48,77728206 | 3,098483111 | 0,623070967 | 4,709394843 | 2,48E-06 | 4,95E-05 |
| EMILIN1 | 128,8480127 | 1,639857566 | 0,347426603 | 4,704790036 | 2,54E-06 | 5,05E-05 |
| MAL | 39,57340648 | 3,464877836 | 0,669770372 | 4,694105682 | 2,68E-06 | 5,29E-05 |
| RORC | 26,13664694 | 2,777760689 | 0,587674155 | 4,684977087 | 2,80E-06 | 5,50E-05 |
| PTK2 | 351,4638039 | 2,477553128 | 0,521587879 | 4,684769133 | 2,80E-06 | 5,50E-05 |
| GYPE | 142,7763228 | 1,891528989 | 0,401385726 | 4,684246085 | 2,81E-06 | 5,51E-05 |
| AGAP3 | 949,5526516 | 1,041453001 | 0,222716515 | 4,677427311 | 2,90E-06 | 5,67E-05 |
| NLRP6 | 362,0418812 | 1,688567398 | 0,382816417 | 4,676545909 | 2,92E-06 | 5,69E-05 |
| CCR3 | 57,93643764 | 2,803902646 | 0,582266843 | 4,674047945 | 2,95E-06 | 5,75E-05 |
| MYO7A | 33,55637115 | 3,737964548 | 0,684756816 | 4,66937877 | 3,02E-06 | 5,86E-05 |
| NEIL1 | 1497,44199 | 1,453142161 | 0,310129545 | 4,667908927 | 3,04E-06 | 5,90E-05 |
| ZMAT4 | 216,068965 | 2,640467911 | 0,558154412 | 4,643769643 | 3,42E-06 | 6,54E-05 |
| GATA6-AS1 | 11,54308557 | 3,066840867 | 0,629454234 | 4,630479567 | 3,65E-06 | 6,93E-05 |
| KLRC1 | 9685,022474 | 1,004146993 | 0,21694846 | 4,629659228 | 3,66E-06 | 6,95E-05 |
| TNFSF4 | 139,7637985 | 2,011664798 | 0,427744978 | 4,626434948 | 3,72E-06 | 7,02E-05 |
| JUP | 58,15443641 | 2,526880336 | 0,546290366 | 4,616890407 | 3,90E-06 | 7,28E-05 |
| BTBD6P1 | 302,4428418 | 2,666889986 | 0,563167772 | 4,614916427 | 3,93E-06 | 7,32E-05 |
| KLHL29 | 33,08738174 | 1,946606948 | 0,416347485 | 4,61404557 | 3,95E-06 | 7,34E-05 |
| LINC01619 | 61,33308047 | 2,086410116 | 0,459723481 | 4,610883455 | 4,01E-06 | 7,44E-05 |
| PER2 | 289,3529778 | 1,149889389 | 0,249251779 | 4,609300045 | 4,04E-06 | 7,49E-05 |

| | | | | | | |
|---|---|---|---|---|---|---|
| NAB2 | 282,4119613 | 1,650115906 | 0,354901133 | 4,605926938 | 4,11E-06 | 7,58E-05 |
| PDK4 | 62,19623696 | 2,612015818 | 0,625512614 | 4,597283482 | 4,28E-06 | 7,87E-05 |
| OR2W3 | 8,537974039 | 2,71906961 | 0,575346458 | 4,588680942 | 4,46E-06 | 8,17E-05 |
| RNA5-8SN2 | 83,66427081 | 1,424126424 | 0,309865244 | 4,576279252 | 4,73E-06 | 8,61E-05 |
| TRIM47 | 230,944992 | 2,515587478 | 0,57857664 | 4,576015626 | 4,74E-06 | 8,62E-05 |
| PDE9A | 28,46432687 | 2,11835182 | 0,463611347 | 4,5757863 | 4,74E-06 | 8,62E-05 |
| RAMP1 | 1416,771374 | 2,003743871 | 0,436861022 | 4,569071259 | 4,90E-06 | 8,87E-05 |
| FES | 14096,02622 | 1,218324888 | 0,267207028 | 4,565206498 | 4,99E-06 | 9,01E-05 |
| AC012435.2 | 34,70171456 | 2,115485105 | 0,462083466 | 4,556526026 | 5,20E-06 | 9,35E-05 |
| CAMK1 | 712,271817 | 1,318506905 | 0,289591149 | 4,541444188 | 5,59E-06 | 9,97E-05 |
| CKMT2 | 21,75960717 | 2,975202182 | 0,64829 | 4,540229443 | 5,62E-06 | 0,000100095 |
| CYP2E1 | 41,85003279 | 1,868522619 | 0,4155738 | 4,538337814 | 5,67E-06 | 0,000100908 |
| MCF2L | 98,84718277 | 2,428602121 | 0,531561162 | 4,537497838 | 5,69E-06 | 0,000101221 |
| HDAC11 | 197,7692003 | 1,658304137 | 0,367992609 | 4,521049269 | 6,15E-06 | 0,000108744 |
| AC092171.3 | 4,066207505 | 3,35422722 | 0,680565705 | 4,517883731 | 6,25E-06 | 0,000110189 |
| LINC01232 | 60,31393813 | 1,043585838 | 0,231126021 | 4,512542166 | 6,41E-06 | 0,000112804 |
| UBR5-AS1 | 44,37993739 | 1,718926174 | 0,381103323 | 4,504273936 | 6,66E-06 | 0,000116878 |
| WNT6 | 5,052215367 | 3,331058752 | 0,675564228 | 4,500825894 | 6,77E-06 | 0,000118378 |
| DIO3OS | 25,74357413 | 3,117712453 | 0,653715159 | 4,496033198 | 6,92E-06 | 0,000120658 |
| FNBP1L | 19,56801807 | 2,777742174 | 0,598214518 | 4,494566007 | 6,97E-06 | 0,000121283 |
| JHY | 255,9646704 | 1,520655684 | 0,340394715 | 4,494675056 | 6,97E-06 | 0,000121283 |
| XCL2 | 7701,308734 | 1,762187493 | 0,390203808 | 4,49211141 | 7,05E-06 | 0,000122479 |
| CSTF2T | 3840,28611 | 1,005275341 | 0,22378182 | 4,490371606 | 7,11E-06 | 0,000123377 |
| PRKAR2B | 149,6290659 | 2,645619867 | 0,579069628 | 4,480688106 | 7,44E-06 | 0,000128777 |
| PLS3 | 37,18201753 | 2,874024246 | 0,664410669 | 4,478669708 | 7,51E-06 | 0,000129778 |
| CITED4 | 54,10013354 | 1,04923707 | 0,233115122 | 4,465966524 | 7,97E-06 | 0,00013737 |
| MGAT5B | 72,42671081 | 2,704211638 | 0,596935983 | 4,462644629 | 8,10E-06 | 0,00013928 |
| BMS1P1 | 120,6512927 | 1,155984636 | 0,261025163 | 4,453962399 | 8,43E-06 | 0,000144666 |

| | | | | | | |
|---|---|---|---|---|---|---|
| DOCK1 | 19,87347216 | 2,858351435 | 0,618826085 | 4,453308541 | 8,46E-06 | 0,000144984 |
| MPV17L | 18,52482589 | 2,933379987 | 0,630792433 | 4,438740939 | 9,05E-06 | 0,000153713 |
| KANSL1L-AS1 | 57,80704568 | 1,175597107 | 0,265021626 | 4,438865951 | 9,04E-06 | 0,000153713 |
| FAM117B | 398,6916273 | 1,703561568 | 0,38370197 | 4,431446356 | 9,36E-06 | 0,00015834 |
| AC009133.3 | 89,18313721 | 1,272998877 | 0,288556644 | 4,428772591 | 9,48E-06 | 0,000159779 |
| AC008014.1 | 81,58722507 | 1,565423115 | 0,352980341 | 4,420221681 | 9,86E-06 | 0,00016568 |
| PAOX | 277,1754213 | 1,163805828 | 0,264060277 | 4,410719008 | 1,03E-05 | 0,000172546 |
| P4HA2 | 62,19190001 | 2,074598291 | 0,470662803 | 4,403734272 | 1,06E-05 | 0,000177903 |
| CCDC121 | 55,15150199 | 1,36373148 | 0,310334972 | 4,399393755 | 1,09E-05 | 0,000180899 |
| IL3 | 7,263746933 | 3,032465126 | 0,685089311 | 4,39804176 | 1,09E-05 | 0,000181879 |
| HOXA5 | 29,22251332 | 2,663573703 | 0,628071499 | 4,396650806 | 1,10E-05 | 0,000182746 |
| TNFRSF11A | 900,3745533 | 3,043200414 | 0,652565419 | 4,393640665 | 1,11E-05 | 0,000184991 |
| PCNX2 | 244,8780474 | 1,353694323 | 0,30927685 | 4,37962019 | 1,19E-05 | 0,000196172 |
| KRT81 | 219,3263906 | 1,852860789 | 0,420643076 | 4,376345734 | 1,21E-05 | 0,000198816 |
| AC123912.4 | 57,7867272 | 3,215074261 | 0,670750072 | 4,370088695 | 1,24E-05 | 0,000203932 |
| BMF | 108,750818 | 1,85337081 | 0,424311083 | 4,354033382 | 1,34E-05 | 0,000217868 |
| HSPG2 | 17,117014 | 2,187090022 | 0,677115263 | 4,335947056 | 1,45E-05 | 0,000235248 |
| SLC5A2 | 15,49691469 | 2,370559637 | 0,57441768 | 4,324722237 | 1,53E-05 | 0,000247153 |
| RASSF2 | 2129,258914 | 1,4305824 | 0,330369984 | 4,321751259 | 1,55E-05 | 0,000250104 |
| CEP68 | 758,1683459 | 1,69136332 | 0,390381931 | 4,320330329 | 1,56E-05 | 0,000251116 |
| NEK6 | 534,3137535 | 1,209174135 | 0,281884172 | 4,313087311 | 1,61E-05 | 0,000259282 |
| AL136456.1 | 15,04858929 | 2,661349463 | 0,605572047 | 4,297893561 | 1,72E-05 | 0,000275504 |
| CYGB | 340,0955644 | 2,588938845 | 0,590623579 | 4,292018938 | 1,77E-05 | 0,000282226 |
| UBASH3A | 71,54453592 | 2,887247625 | 0,635371147 | 4,289809408 | 1,79E-05 | 0,000284823 |
| CSPG4P12 | 36,1560947 | 1,837829876 | 0,435732667 | 4,287569921 | 1,81E-05 | 0,000287255 |
| NFIX | 60,68310508 | 2,522417396 | 0,573915819 | 4,28194832 | 1,85E-05 | 0,000293454 |
| PECAM1 | 2770,661028 | 1,234226615 | 0,29372387 | 4,27280116 | 1,93E-05 | 0,000304801 |
| TTLL10 | 55,17994789 | 3,165533647 | 0,677545215 | 4,266103255 | 1,99E-05 | 0,000312867 |

| | | | | | | |
|---|---|---|---|---|---|---|
| CARD10 | 195,7946157 | 3,162507469 | 0,681788746 | 4,254657742 | 2,09E-05 | 0,000328161 |
| BASP1 | 89,17467743 | 2,459015851 | 0,534617824 | 4,252059772 | 2,12E-05 | 0,000331342 |
| AC093157.1 | 224,0533513 | 1,180037207 | 0,276519418 | 4,250369494 | 2,13E-05 | 0,000333594 |
| AC079779.1 | 58,29398965 | 2,129901217 | 0,592652134 | 4,246268326 | 2,17E-05 | 0,000338968 |
| WASIR2 | 8,636171076 | 2,893649754 | 0,687707799 | 4,245385515 | 2,18E-05 | 0,000339256 |
| CELSR1 | 44,13666181 | 2,481443117 | 0,603597699 | 4,245644287 | 2,18E-05 | 0,000339256 |
| PKIA | 78,85937452 | 2,204573982 | 0,521138808 | 4,238829798 | 2,25E-05 | 0,00034824 |
| PDGFA | 21,16065015 | 2,465664476 | 0,576959244 | 4,220608974 | 2,44E-05 | 0,000376173 |
| BEX5 | 89,58593359 | 1,367130389 | 0,322809632 | 4,216438277 | 2,48E-05 | 0,000382609 |
| CLEC3B | 9,930720348 | 2,837184486 | 0,656641233 | 4,211959984 | 2,53E-05 | 0,000389978 |
| AK8 | 8,325888658 | 2,258884646 | 0,536740694 | 4,201875242 | 2,65E-05 | 0,000405598 |
| AC005083.1 | 9,836329602 | 2,579657893 | 0,645084134 | 4,201039134 | 2,66E-05 | 0,00040679 |
| INPP5F | 829,2714252 | 1,197923774 | 0,28491769 | 4,190833732 | 2,78E-05 | 0,000423278 |
| HOXA-AS3 | 29,76508842 | 2,251069243 | 0,535071714 | 4,188372622 | 2,81E-05 | 0,000427249 |
| PDCD4 | 11931,50272 | 1,092133935 | 0,260805255 | 4,186306354 | 2,84E-05 | 0,000430181 |
| FUT6 | 3,392327544 | 2,858222968 | 0,677162137 | 4,18386062 | 2,87E-05 | 0,000433859 |
| AL022238.4 | 222,1358653 | 1,115870744 | 0,267411959 | 4,183134943 | 2,88E-05 | 0,00043492 |
| CSPG4P10 | 50,37921758 | 1,015984266 | 0,242249925 | 4,177084633 | 2,95E-05 | 0,000445644 |
| IGLV2-18 | 3,924084235 | 2,611510241 | 0,663150589 | 4,173288094 | 3,00E-05 | 0,000451786 |
| TLCD2 | 26,04526924 | 1,633004101 | 0,410515131 | 4,169477262 | 3,05E-05 | 0,000458039 |
| AC025164.2 | 773,1323373 | 1,46411102 | 0,353146935 | 4,16917934 | 3,06E-05 | 0,000458298 |
| AC068775.1 | 884,7281123 | 1,09932535 | 0,260790028 | 4,165733973 | 3,10E-05 | 0,000463895 |
| MVB12B | 297,7737086 | 1,406102328 | 0,347569303 | 4,156290574 | 3,23E-05 | 0,000479572 |
| TTC24 | 68,77921346 | 2,267286146 | 0,562949671 | 4,149431782 | 3,33E-05 | 0,000492719 |
| EPS8 | 127,5161962 | 1,415747403 | 0,355555925 | 4,13439404 | 3,56E-05 | 0,000521907 |
| CARD19 | 1476,40296 | 1,100074256 | 0,26648286 | 4,126729715 | 3,68E-05 | 0,000537257 |
| PLD1 | 141,5092009 | 1,383133178 | 0,339080294 | 4,125699483 | 3,70E-05 | 0,000538894 |
| RETREG1 | 94,16902923 | 2,07510904 | 0,512433511 | 4,114913381 | 3,87E-05 | 0,000560662 |

| | | | | | | |
|---|---|---|---|---|---|---|
| MCC | 133,7813786 | 1,435245654 | 0,362675489 | 4,109445865 | 3,97E-05 | 0,000572868 |
| STXBP1 | 166,6807384 | 1,773062233 | 0,430974083 | 4,105823205 | 4,03E-05 | 0,000580547 |
| ZEB1-AS1 | 164,9176877 | 1,566477904 | 0,388516549 | 4,102143778 | 4,09E-05 | 0,000589145 |
| NECTIN3 | 65,65558893 | 2,109821104 | 0,522511446 | 4,100453329 | 4,12E-05 | 0,000593043 |
| BMP8B | 158,3523606 | 1,797732771 | 0,428864179 | 4,094667742 | 4,23E-05 | 0,000605456 |
| TMEM14C | 657,2655348 | 1,810962691 | 0,442101686 | 4,077193222 | 4,56E-05 | 0,000647929 |
| EPHA4 | 1050,413239 | 1,557123797 | 0,38537098 | 4,065706375 | 4,79E-05 | 0,000676663 |
| FAM30A | 12,43469716 | 2,62472664 | 0,687276262 | 4,064282968 | 4,82E-05 | 0,00068015 |
| LGALS3BP | 455,2933785 | 2,01736213 | 0,498440034 | 4,063485753 | 4,83E-05 | 0,000681702 |
| CNR2 | 2219,199414 | 2,444235085 | 0,585082679 | 4,058692112 | 4,93E-05 | 0,000692456 |
| ZNF843 | 15,55036673 | 2,364271664 | 0,592664421 | 4,057948375 | 4,95E-05 | 0,000693699 |
| AC036176.1 | 50,40960634 | 1,284314447 | 0,317099019 | 4,055862318 | 4,99E-05 | 0,000698464 |
| CYP4F22 | 115,5672067 | 1,023809783 | 0,254208061 | 4,05374212 | 5,04E-05 | 0,000703365 |
| SIX4 | 5,032002761 | 2,972599722 | 0,687818501 | 4,049103353 | 5,14E-05 | 0,000714977 |
| DTX1 | 44,33735311 | 2,597561359 | 0,615597976 | 4,038247269 | 5,39E-05 | 0,000744769 |
| PCBP3 | 14,05993545 | 2,732766327 | 0,655409566 | 4,033752225 | 5,49E-05 | 0,000757091 |
| COL15A1 | 46,51217042 | 2,389977593 | 0,542911588 | 4,032154516 | 5,53E-05 | 0,000761736 |
| AL138828.1 | 4,740326734 | 2,687186184 | 0,658844736 | 4,030351437 | 5,57E-05 | 0,000767079 |
| AL135818.2 | 33,98399152 | 1,492962774 | 0,370720041 | 4,027813163 | 5,63E-05 | 0,000774348 |
| SERPINB9P1 | 37,81299154 | 1,718317668 | 0,425459062 | 4,022438429 | 5,76E-05 | 0,00079062 |
| H3-2 | 22,33116103 | 1,607520491 | 0,393806175 | 4,018284767 | 5,86E-05 | 0,00080359 |
| LTB | 7685,961389 | 1,885747477 | 0,465054567 | 4,015437499 | 5,93E-05 | 0,000811148 |
| MEST | 60,3144696 | 1,50647934 | 0,379673843 | 4,006372315 | 6,17E-05 | 0,00083779 |
| HOXA10 | 474,6676523 | 1,586323541 | 0,395748298 | 4,00277112 | 6,26E-05 | 0,000848934 |
| KL | 6,183304183 | 2,588017678 | 0,687354751 | 3,999147664 | 6,36E-05 | 0,000861015 |
| SELL | 61133,19386 | 1,046801587 | 0,263254611 | 3,99281777 | 6,53E-05 | 0,000881824 |
| NR2F6 | 281,6955243 | 1,506947823 | 0,37795197 | 3,99255195 | 6,54E-05 | 0,000882222 |
| RGS6 | 4,488837285 | 2,58082069 | 0,666501808 | 3,987713893 | 6,67E-05 | 0,000897999 |

| | | | | | | |
|---|---|---|---|---|---|---|
| YWHAEP7 | 38,3378469 | 2,636322159 | 0,645170748 | 3,984996126 | 6,75E-05 | 0,00090592 |
| CHMP4C | 34,73443287 | 2,848940037 | 0,671522152 | 3,970921553 | 7,16E-05 | 0,000954791 |
| ABTB2 | 42,18544019 | 1,874409248 | 0,473912327 | 3,968552991 | 7,23E-05 | 0,000961785 |

**Supplementary table 3.** List of ST2$^{-}$ CD56$^{dim}$ NK cell signature genes used in this study, Related to Fig. 3B, 3E, 3H, 3I, 5A, 5B, Supplementary Fig. S4E, S9A

| gene | baseMean | log2FoldChange | lfcSE | stat | pvalue | padj |
|---|---|---|---|---|---|---|
| FGFBP2 | 8144,153518 | 3,202150845 | 0,320967185 | 10,00234058 | 1,49E-23 | 9,84E-20 |
| RNF166 | 8329,004271 | 1,572015459 | 0,157231451 | 10,01458146 | 1,32E-23 | 9,84E-20 |
| RASGRP2 | 4442,914764 | 1,388044602 | 0,146010973 | 9,525583883 | 1,64E-21 | 6,51E-18 |
| DPEP2 | 1526,11809 | 1,609829657 | 0,190756897 | 8,483359095 | 2,19E-17 | 4,34E-14 |
| AL353622.1 | 297,1803946 | 1,685555006 | 0,200170725 | 8,443427493 | 3,08E-17 | 5,56E-14 |
| KLF3 | 2317,609775 | 1,540112022 | 0,183942681 | 8,40503739 | 4,28E-17 | 7,07E-14 |
| CEP78 | 4905,929088 | 1,044810306 | 0,126366368 | 8,259780569 | 1,46E-16 | 2,07E-13 |
| LINC00944 | 44,76489599 | 3,21482747 | 0,372462803 | 8,19595732 | 2,49E-16 | 3,29E-13 |
| DYRK1B | 672,4335557 | 1,118315017 | 0,137280114 | 8,165574615 | 3,20E-16 | 3,86E-13 |
| P2RY8 | 7887,303735 | 1,486780838 | 0,189003159 | 7,86287299 | 3,75E-15 | 3,24E-12 |
| GPA33 | 161,1699077 | 2,418507998 | 0,327658938 | 7,546200536 | 4,48E-14 | 3,07E-11 |
| AC245407.2 | 406,0213999 | 1,859932789 | 0,248929722 | 7,516379506 | 5,63E-14 | 3,72E-11 |
| TMEM71 | 1574,466452 | 1,954352228 | 0,268749796 | 7,358825676 | 1,86E-13 | 1,12E-10 |
| AL138824.1 | 129,6429159 | 2,422070604 | 0,336058541 | 7,33023171 | 2,30E-13 | 1,30E-10 |
| LINC00861 | 3061,05769 | 1,53354884 | 0,210763177 | 7,323411026 | 2,42E-13 | 1,33E-10 |
| KLF2 | 6970,676213 | 1,560628347 | 0,216566128 | 7,216426459 | 5,34E-13 | 2,71E-10 |
| WAKMAR2 | 491,0159819 | 1,331281734 | 0,186211266 | 7,186350801 | 6,65E-13 | 3,30E-10 |
| RASSF1-AS1 | 238,3238734 | 1,416845962 | 0,196731197 | 7,170698638 | 7,46E-13 | 3,52E-10 |
| MCM3AP-AS1 | 326,1585037 | 1,138706429 | 0,161235728 | 7,053431474 | 1,75E-12 | 7,37E-10 |
| AC005332.2 | 94,85373685 | 2,095310385 | 0,293433439 | 6,976915203 | 3,02E-12 | 1,11E-09 |
| BISPR | 1681,320381 | 1,060832931 | 0,152324166 | 6,960487766 | 3,39E-12 | 1,22E-09 |
| SELPLG | 21499,71541 | 1,283065741 | 0,185716133 | 6,934799516 | 4,07E-12 | 1,39E-09 |
| OAS1 | 1195,66649 | 1,211755432 | 0,175809612 | 6,871013811 | 6,37E-12 | 2,07E-09 |
| AL662844.4 | 967,2315141 | 1,207738148 | 0,177139872 | 6,840280625 | 7,90E-12 | 2,53E-09 |

| | | | | | | |
|---|---|---|---|---|---|---|
| IFIT2 | 1117,024786 | 2,133274979 | 0,315695154 | 6,834721634 | 8,22E-12 | 2,59E-09 |
| RASSF1 | 11231,77473 | 1,301637387 | 0,192592714 | 6,758344733 | 1,40E-11 | 3,90E-09 |
| STMN3 | 137,9011327 | 2,372036606 | 0,352935904 | 6,73337158 | 1,66E-11 | 4,44E-09 |
| AC073130.3 | 171,1204581 | 1,663210743 | 0,247089371 | 6,720080459 | 1,82E-11 | 4,56E-09 |
| HSBP1L1 | 213,0495397 | 1,096455411 | 0,163684666 | 6,690234449 | 2,23E-11 | 5,46E-09 |
| KIFC3 | 745,2824955 | 2,12908314 | 0,321380577 | 6,679504561 | 2,40E-11 | 5,66E-09 |
| AL592295.5 | 158,8162533 | 1,063282332 | 0,160343558 | 6,661930468 | 2,70E-11 | 6,16E-09 |
| HEATR9 | 73,23745131 | 2,555095193 | 0,387866469 | 6,65222127 | 2,89E-11 | 6,36E-09 |
| AC055839.2 | 298,5800799 | 1,540998324 | 0,233523327 | 6,636494986 | 3,21E-11 | 7,00E-09 |
| NDRG1 | 1077,893796 | 1,288213421 | 0,195811703 | 6,60289876 | 4,03E-11 | 8,38E-09 |
| PLEKHG3 | 8342,958982 | 1,360164232 | 0,206675366 | 6,586993772 | 4,49E-11 | 9,18E-09 |
| PARP12 | 6641,277294 | 1,200871467 | 0,18278299 | 6,561481629 | 5,33E-11 | 1,07E-08 |
| FGR | 17120,1803 | 1,4607729 | 0,222708926 | 6,556003566 | 5,53E-11 | 1,09E-08 |
| ZFHX2-AS1 | 761,7303199 | 1,065192305 | 0,162302597 | 6,536673788 | 6,29E-11 | 1,19E-08 |
| THRA | 284,6004293 | 1,592549138 | 0,2455052 | 6,533669042 | 6,42E-11 | 1,20E-08 |
| ICAM2 | 3207,579092 | 1,124983666 | 0,172111497 | 6,529980219 | 6,58E-11 | 1,22E-08 |
| MHENCR | 224,0159648 | 1,361939827 | 0,209491017 | 6,521826752 | 6,95E-11 | 1,28E-08 |
| RENBP | 199,7535366 | 1,483512111 | 0,229877225 | 6,484212226 | 8,92E-11 | 1,62E-08 |
| TTC38 | 8039,041486 | 1,003443505 | 0,159254813 | 6,304267394 | 2,90E-10 | 4,54E-08 |
| NATD1 | 671,0286088 | 1,052857164 | 0,167698493 | 6,275188997 | 3,49E-10 | 5,25E-08 |
| CTSF | 1165,042147 | 1,941891463 | 0,322774126 | 6,247305527 | 4,18E-10 | 6,09E-08 |
| SFXN3 | 1817,137939 | 1,061585468 | 0,170287364 | 6,239116516 | 4,40E-10 | 6,35E-08 |
| DLGAP1-AS1 | 330,5404792 | 1,279252449 | 0,205533663 | 6,233047148 | 4,57E-10 | 6,41E-08 |
| TTLL3 | 2088,501966 | 1,21159376 | 0,194420136 | 6,232022848 | 4,60E-10 | 6,41E-08 |
| AC244157.2 | 11,20080967 | 3,129269147 | 0,478217362 | 6,211872602 | 5,24E-10 | 6,97E-08 |
| PATL2 | 4331,057083 | 2,073692218 | 0,339076125 | 6,142247184 | 8,14E-10 | 1,01E-07 |
| SNAI3 | 591,4513799 | 1,664734223 | 0,274430177 | 6,141609469 | 8,17E-10 | 1,01E-07 |
| ZMYND10 | 90,55930117 | 1,479130226 | 0,24144219 | 6,144421879 | 8,03E-10 | 1,01E-07 |

| | | | | | | |
|---|---|---|---|---|---|---|
| ZNF276 | 8281,915905 | 1,111608858 | 0,18130732 | 6,140495038 | 8,23E-10 | 1,01E-07 |
| TRIM22 | 16223,92092 | 1,132720837 | 0,186423812 | 6,070675197 | 1,27E-09 | 1,47E-07 |
| AL139246.5 | 210,2164121 | 1,289055593 | 0,214307842 | 6,02576554 | 1,68E-09 | 1,81E-07 |
| BTN3A3 | 7879,327212 | 1,001665785 | 0,167620932 | 5,979652651 | 2,24E-09 | 2,27E-07 |
| CCDC88C | 5657,368226 | 1,023107223 | 0,171581502 | 5,974768528 | 2,30E-09 | 2,33E-07 |
| SLAMF6 | 4918,979345 | 1,245100796 | 0,210034001 | 5,943449696 | 2,79E-09 | 2,74E-07 |
| FCGR2A | 84,56463711 | 2,695351865 | 0,468057246 | 5,934925165 | 2,94E-09 | 2,83E-07 |
| CDC14B | 109,090885 | 2,543603082 | 0,417798191 | 5,929046971 | 3,05E-09 | 2,91E-07 |
| DEGS2 | 171,5630993 | 2,213052873 | 0,37606735 | 5,890150226 | 3,86E-09 | 3,54E-07 |
| FAM13A-AS1 | 86,14944071 | 1,321665729 | 0,224337026 | 5,88294354 | 4,03E-09 | 3,60E-07 |
| LINC002481 | 1047,622692 | 1,077615531 | 0,183488496 | 5,87767994 | 4,16E-09 | 3,65E-07 |
| NLRP1 | 11216,96825 | 1,109626639 | 0,189979648 | 5,855860413 | 4,75E-09 | 4,06E-07 |
| EFEMP2 | 692,2367593 | 1,341301299 | 0,231765649 | 5,815466941 | 6,05E-09 | 4,96E-07 |
| AC006369.1 | 101,647403 | 1,990066631 | 0,35352843 | 5,813465678 | 6,12E-09 | 4,96E-07 |
| TRGV10 | 614,0797893 | 1,416106431 | 0,241708205 | 5,811665992 | 6,19E-09 | 4,97E-07 |
| PRR29 | 306,0053646 | 1,386872293 | 0,238445664 | 5,79664335 | 6,77E-09 | 5,35E-07 |
| ZFP36L2 | 18485,19308 | 1,390293461 | 0,239873773 | 5,788011561 | 7,12E-09 | 5,58E-07 |
| METTL7A | 1308,623924 | 1,73695712 | 0,300854347 | 5,780965031 | 7,43E-09 | 5,80E-07 |
| SEPTIN4 | 75,97567771 | 2,698194832 | 0,452517403 | 5,75264716 | 8,79E-09 | 6,63E-07 |
| AL391987.4 | 20,05432848 | 2,197526065 | 0,378590689 | 5,734426497 | 9,78E-09 | 7,21E-07 |
| BCL9L | 506,2869482 | 1,200452362 | 0,211099255 | 5,706615393 | 1,15E-08 | 8,25E-07 |
| C3orf18 | 189,7095457 | 1,494737274 | 0,261434877 | 5,677302098 | 1,37E-08 | 9,52E-07 |
| DNAI2 | 26,98773266 | 2,158699082 | 0,377378171 | 5,675041668 | 1,39E-08 | 9,62E-07 |
| AC008555.4 | 342,8896863 | 1,305132998 | 0,230246837 | 5,671671372 | 1,41E-08 | 9,74E-07 |
| AC083862.3 | 93,04402387 | 1,605951846 | 0,285482801 | 5,666288702 | 1,46E-08 | 9,98E-07 |
| AOAH | 15162,22478 | 1,177424409 | 0,212304547 | 5,666244743 | 1,46E-08 | 9,98E-07 |
| SCIMP | 212,0349226 | 2,783228915 | 0,450582846 | 5,664457042 | 1,47E-08 | 1,01E-06 |
| A2M-AS1 | 33,70740077 | 2,023754653 | 0,369565554 | 5,654876374 | 1,56E-08 | 1,04E-06 |

| | | | | | | |
|---|---|---|---|---|---|---|
| PROCR | 189,2710625 | 1,782388913 | 0,308461157 | 5,655020804 | 1,56E-08 | 1,04E-06 |
| HLA-DPB1 | 3374,062135 | 1,455744026 | 0,254509239 | 5,591878485 | 2,25E-08 | 1,37E-06 |
| THEMIS2 | 6429,654639 | 1,318426828 | 0,243695999 | 5,581361969 | 2,39E-08 | 1,45E-06 |
| TTYH2 | 119,2345397 | 1,912936214 | 0,330242527 | 5,570418879 | 2,54E-08 | 1,50E-06 |
| FCHO2 | 354,5028786 | 1,313542877 | 0,235595382 | 5,550212185 | 2,85E-08 | 1,64E-06 |
| SYNE1 | 7987,350074 | 1,037980237 | 0,187983316 | 5,549434563 | 2,87E-08 | 1,64E-06 |
| ANO8 | 97,04963606 | 1,230117118 | 0,221829997 | 5,547661657 | 2,90E-08 | 1,65E-06 |
| AL731571.1 | 505,1583218 | 1,013699118 | 0,182893325 | 5,543829085 | 2,96E-08 | 1,66E-06 |
| ARVCF | 833,3125729 | 2,408541464 | 0,439432257 | 5,524252065 | 3,31E-08 | 1,81E-06 |
| SMAD7 | 2965,651036 | 1,11518809 | 0,202018956 | 5,511917559 | 3,55E-08 | 1,91E-06 |
| UCP3 | 56,41155725 | 1,573797693 | 0,285729882 | 5,507325675 | 3,64E-08 | 1,95E-06 |
| LIME1 | 942,3741088 | 1,479021292 | 0,267520994 | 5,506762372 | 3,65E-08 | 1,95E-06 |
| AC133552.2 | 182,303471 | 1,054617444 | 0,192419877 | 5,500330174 | 3,79E-08 | 1,99E-06 |
| NUAK2 | 1515,700571 | 1,857743056 | 0,334990288 | 5,499802944 | 3,80E-08 | 2,00E-06 |
| AP001372.2 | 131,5564837 | 1,148468837 | 0,210424808 | 5,469934728 | 4,50E-08 | 2,30E-06 |
| AL160269.1 | 17,96116089 | 2,732927978 | 0,50061574 | 5,464840779 | 4,63E-08 | 2,36E-06 |
| MC1R | 484,7686664 | 1,07744908 | 0,196701472 | 5,458544783 | 4,80E-08 | 2,40E-06 |
| AC015911.3 | 80,38809773 | 1,337714978 | 0,247579608 | 5,45226435 | 4,97E-08 | 2,47E-06 |
| CLIC3 | 9459,677847 | 1,481450713 | 0,279632872 | 5,441172466 | 5,29E-08 | 2,57E-06 |
| PRR5 | 1701,267869 | 1,29351842 | 0,235458066 | 5,439890922 | 5,33E-08 | 2,59E-06 |
| C8G | 45,28819125 | 1,659126755 | 0,308272489 | 5,436019139 | 5,45E-08 | 2,63E-06 |
| VIPR2 | 140,3095731 | 2,66019421 | 0,459253371 | 5,43117563 | 5,60E-08 | 2,68E-06 |
| AMZ2P1 | 484,3583591 | 1,158245381 | 0,213752863 | 5,421662242 | 5,90E-08 | 2,77E-06 |
| TMEM191A | 149,1107394 | 1,328338074 | 0,245783935 | 5,419654336 | 5,97E-08 | 2,77E-06 |
| AC093010.2 | 133,6541972 | 1,438341241 | 0,262590937 | 5,414098905 | 6,16E-08 | 2,82E-06 |
| LINC01801 | 99,14112127 | 1,834082888 | 0,337712839 | 5,409646324 | 6,31E-08 | 2,86E-06 |
| CXXC4 | 7,761928407 | 3,341815889 | 0,592915329 | 5,401131918 | 6,62E-08 | 2,96E-06 |
| COL6A2 | 1751,980505 | 2,12446352 | 0,391644071 | 5,398399147 | 6,72E-08 | 3,00E-06 |

| | | | | | | |
|---|---|---|---|---|---|---|
| LINC00565 | 51,79099498 | 2,056481532 | 0,380334703 | 5,383884267 | 7,29E-08 | 3,17E-06 |
| AC021188.1 | 181,0353108 | 1,059744646 | 0,197249743 | 5,382760231 | 7,34E-08 | 3,18E-06 |
| ZNF683 | 1353,273478 | 2,206563322 | 0,407730698 | 5,365112229 | 8,09E-08 | 3,39E-06 |
| HCG27 | 22,85714239 | 2,036093232 | 0,387623816 | 5,365334247 | 8,08E-08 | 3,39E-06 |
| LINC00943 | 31,06372798 | 3,170285752 | 0,524871488 | 5,363474766 | 8,16E-08 | 3,40E-06 |
| AC015911.11 | 643,1416956 | 1,104234871 | 0,207477992 | 5,354340959 | 8,59E-08 | 3,53E-06 |
| RASA3 | 14648,7847 | 1,227943988 | 0,230592384 | 5,350087119 | 8,79E-08 | 3,59E-06 |
| MXD4 | 4367,991898 | 1,008548164 | 0,189256948 | 5,34861958 | 8,86E-08 | 3,60E-06 |
| TSPAN32 | 1680,349979 | 1,79932189 | 0,336957099 | 5,343106309 | 9,14E-08 | 3,68E-06 |
| CARMIL3 | 114,0695523 | 2,419235059 | 0,442115745 | 5,337392111 | 9,43E-08 | 3,77E-06 |
| AC008878.3 | 449,526451 | 1,286863372 | 0,24350788 | 5,302891786 | 1,14E-07 | 4,37E-06 |
| DBP | 1128,896084 | 1,281597154 | 0,24346066 | 5,294708607 | 1,19E-07 | 4,55E-06 |
| BBS2 | 1317,400007 | 1,010588346 | 0,191435061 | 5,286580295 | 1,25E-07 | 4,69E-06 |
| LEXM | 191,9846548 | 1,794520893 | 0,34204509 | 5,279096728 | 1,30E-07 | 4,84E-06 |
| ZBP1 | 2571,005471 | 1,248883048 | 0,237996161 | 5,27659424 | 1,32E-07 | 4,86E-06 |
| ADGRG1 | 24045,91616 | 2,134403793 | 0,408032775 | 5,273768679 | 1,34E-07 | 4,92E-06 |
| DGKD | 7479,620703 | 1,010624725 | 0,191786194 | 5,273526694 | 1,34E-07 | 4,92E-06 |
| KLF9 | 95,97313961 | 1,239751056 | 0,234357911 | 5,271941387 | 1,35E-07 | 4,95E-06 |
| AL645933.3 | 72,22059645 | 1,355849793 | 0,256626744 | 5,26739582 | 1,38E-07 | 5,05E-06 |
| LINC00299 | 1692,671679 | 1,19453555 | 0,22752385 | 5,265035978 | 1,40E-07 | 5,10E-06 |
| PHOSPHO1 | 155,6505971 | 1,102879801 | 0,209418228 | 5,260066897 | 1,44E-07 | 5,22E-06 |
| VASH1 | 473,3051921 | 1,054677812 | 0,194473484 | 5,257675292 | 1,46E-07 | 5,28E-06 |
| AL365272.1 | 94,70735444 | 1,693224309 | 0,322038434 | 5,241850783 | 1,59E-07 | 5,63E-06 |
| KDM7A | 806,6152019 | 1,000202276 | 0,191750705 | 5,240127635 | 1,60E-07 | 5,67E-06 |
| AC026748.1 | 43,85490864 | 2,347312926 | 0,472272602 | 5,23597803 | 1,64E-07 | 5,77E-06 |
| AC004408.2 | 17,0308697 | 2,529599962 | 0,597942865 | 5,232422875 | 1,67E-07 | 5,85E-06 |
| NMUR1 | 2270,621657 | 1,684435022 | 0,324979943 | 5,230283425 | 1,69E-07 | 5,91E-06 |
| LY9 | 479,5011827 | 2,118962474 | 0,391487208 | 5,228320333 | 1,71E-07 | 5,95E-06 |

| | | | | | | |
|---|---|---|---|---|---|---|
| TSPOAP1 | 3229,099018 | 1,739234161 | 0,330229789 | 5,221416885 | 1,78E-07 | 6,12E-06 |
| CDC42-AS1 | 31,27226099 | 1,79126482 | 0,358407045 | 5,218163321 | 1,81E-07 | 6,20E-06 |
| AL590560.3 | 30,87246874 | 2,067096386 | 0,398254939 | 5,208159089 | 1,91E-07 | 6,44E-06 |
| C9orf139 | 1590,305386 | 1,005446539 | 0,192864819 | 5,19139162 | 2,09E-07 | 6,92E-06 |
| AC023908.3 | 109,2582264 | 1,041530409 | 0,20068853 | 5,186997842 | 2,14E-07 | 7,04E-06 |
| LINC00987 | 56,43830273 | 2,101577291 | 0,388294364 | 5,160013637 | 2,47E-07 | 7,86E-06 |
| SLC43A2 | 246,8527537 | 1,658615653 | 0,308859536 | 5,157946714 | 2,50E-07 | 7,91E-06 |
| LINC00528 | 455,6077964 | 1,04871322 | 0,2034194 | 5,149984738 | 2,61E-07 | 8,16E-06 |
| AC009133.1 | 143,5085013 | 1,162587828 | 0,223297213 | 5,144641867 | 2,68E-07 | 8,37E-06 |
| ATP8A1 | 2191,085121 | 1,016084287 | 0,197428 | 5,142980308 | 2,70E-07 | 8,42E-06 |
| SORL1 | 15644,15909 | 1,237782888 | 0,241760048 | 5,13654422 | 2,80E-07 | 8,59E-06 |
| AC008115.3 | 12,80872293 | 1,749697322 | 0,341449513 | 5,121549903 | 3,03E-07 | 9,07E-06 |
| TLR1 | 991,0522492 | 1,011983935 | 0,19931512 | 5,097226114 | 3,45E-07 | 1,00E-05 |
| NME4 | 398,7327413 | 1,185412126 | 0,246136621 | 5,078882468 | 3,80E-07 | 1,08E-05 |
| LINC02580 | 103,136312 | 1,927552183 | 0,381713593 | 5,075481389 | 3,87E-07 | 1,09E-05 |
| FCMR | 3082,195454 | 1,525517697 | 0,30064693 | 5,06963301 | 3,99E-07 | 1,12E-05 |
| ADAMTS10 | 1817,806232 | 1,476035116 | 0,291159586 | 5,06860452 | 4,01E-07 | 1,12E-05 |
| AXIN1 | 2828,725292 | 1,107000537 | 0,21911331 | 5,054831921 | 4,31E-07 | 1,19E-05 |
| PPM1N | 111,0466079 | 1,86650293 | 0,365824576 | 5,048706704 | 4,45E-07 | 1,22E-05 |
| AC017104.6 | 281,1891744 | 1,924773209 | 0,390438273 | 5,043971505 | 4,56E-07 | 1,24E-05 |
| AC084018.2 | 126,5141249 | 1,028639788 | 0,204398709 | 5,041739495 | 4,61E-07 | 1,26E-05 |
| ADHFE1 | 1716,108649 | 1,300878706 | 0,261954489 | 5,026136826 | 5,00E-07 | 1,34E-05 |
| CALCOCO1 | 8382,387021 | 1,036633552 | 0,20669138 | 5,018369601 | 5,21E-07 | 1,38E-05 |
| YPEL3 | 5250,87534 | 1,182445039 | 0,23676953 | 5,002213151 | 5,67E-07 | 1,48E-05 |
| AC020656.2 | 18,1122843 | 1,867029789 | 0,377135657 | 4,988305634 | 6,09E-07 | 1,55E-05 |
| AC087741.1 | 361,5262212 | 1,184731562 | 0,240995022 | 4,978428649 | 6,41E-07 | 1,62E-05 |
| ASPRV1 | 58,1557482 | 1,222666349 | 0,244679157 | 4,968705871 | 6,74E-07 | 1,69E-05 |
| MAP2K6 | 110,4938357 | 1,232926195 | 0,251886494 | 4,966838123 | 6,81E-07 | 1,70E-05 |

| | | | | | | |
|---|---|---|---|---|---|---|
| AC245100.8 | 344,612282 | 1,124740314 | 0,228653101 | 4,952502178 | 7,33E-07 | 1,79E-05 |
| IL13RA1 | 81,74065765 | 2,92075194 | 0,514790173 | 4,939134248 | 7,85E-07 | 1,90E-05 |
| AC134669.1 | 372,670439 | 1,042972577 | 0,211665566 | 4,925676539 | 8,41E-07 | 1,98E-05 |
| IFIT1 | 167,5701181 | 1,990677333 | 0,432179887 | 4,91394281 | 8,93E-07 | 2,08E-05 |
| BTN3A1 | 11982,80094 | 1,076439364 | 0,219833583 | 4,910972132 | 9,06E-07 | 2,10E-05 |
| C13orf46 | 219,4481016 | 1,604370654 | 0,343040709 | 4,908335023 | 9,19E-07 | 2,12E-05 |
| IL24 | 14,13899697 | 1,829517236 | 0,368439861 | 4,904722928 | 9,36E-07 | 2,15E-05 |
| TXNIP | 87373,28167 | 1,198611463 | 0,24632488 | 4,892742155 | 9,94E-07 | 2,25E-05 |
| PPP2R5C | 10495,0938 | 1,212018087 | 0,248196189 | 4,881872109 | 1,05E-06 | 2,35E-05 |
| PRR5-ARHGAP8 | 30,04910832 | 2,233599954 | 0,459502379 | 4,857117848 | 1,19E-06 | 2,60E-05 |
| TRIM73 | 113,5137089 | 1,326773633 | 0,270785003 | 4,832941441 | 1,35E-06 | 2,83E-05 |
| MST1L | 35,53330089 | 1,843963043 | 0,39785379 | 4,826925457 | 1,39E-06 | 2,90E-05 |
| AC009093.10 | 70,90865127 | 1,303881935 | 0,271860718 | 4,825710292 | 1,40E-06 | 2,91E-05 |
| NR1D2 | 1901,099841 | 1,061516657 | 0,220954674 | 4,805364677 | 1,54E-06 | 3,14E-05 |
| SLFN12L | 388,6582447 | 1,016747552 | 0,212823659 | 4,804996628 | 1,55E-06 | 3,14E-05 |
| DAPK2 | 315,8775139 | 2,269275673 | 0,46000889 | 4,803430519 | 1,56E-06 | 3,16E-05 |
| AL162457.1 | 60,01120291 | 1,80702327 | 0,371410401 | 4,798244632 | 1,60E-06 | 3,22E-05 |
| NHSL2 | 1708,883997 | 1,543158551 | 0,327110587 | 4,794206828 | 1,63E-06 | 3,27E-05 |
| UNC5CL | 93,2975135 | 1,57197324 | 0,329561404 | 4,788580335 | 1,68E-06 | 3,33E-05 |
| AL162458.1 | 165,1223187 | 1,127248748 | 0,23570295 | 4,789075463 | 1,68E-06 | 3,33E-05 |
| AC010319.1 | 43,13147385 | 1,502921542 | 0,313091723 | 4,786588652 | 1,70E-06 | 3,36E-05 |
| DLGAP1-AS2 | 58,74548733 | 1,427003542 | 0,297323381 | 4,786150878 | 1,70E-06 | 3,36E-05 |
| ZNF208 | 25,17114763 | 1,843243073 | 0,394563582 | 4,783253633 | 1,72E-06 | 3,40E-05 |
| AC243829.1 | 40,37717938 | 2,533252857 | 0,52705675 | 4,780751422 | 1,75E-06 | 3,43E-05 |
| CCDC65 | 186,1724057 | 2,427617547 | 0,513646417 | 4,779372081 | 1,76E-06 | 3,45E-05 |
| SAT1 | 3820,59353 | 1,023904591 | 0,214341376 | 4,773329599 | 1,81E-06 | 3,52E-05 |
| ZNF831 | 1478,234274 | 1,148277783 | 0,240483824 | 4,772403024 | 1,82E-06 | 3,53E-05 |
| AC018809.1 | 30,58052564 | 1,127351416 | 0,235538174 | 4,758501657 | 1,95E-06 | 3,71E-05 |

| | | | | | | |
|---|---|---|---|---|---|---|
| FRMPD3 | 25,27426586 | 2,242602646 | 0,492422158 | 4,745952303 | 2,08E-06 | 3,90E-05 |
| C1orf21 | 3800,709353 | 1,152410123 | 0,243213472 | 4,745811927 | 2,08E-06 | 3,90E-05 |
| AC099489.1 | 97,36576781 | 1,972012994 | 0,414899076 | 4,739572019 | 2,14E-06 | 3,99E-05 |
| PLCD1 | 824,4581922 | 1,334017381 | 0,282996846 | 4,739078882 | 2,15E-06 | 3,99E-05 |
| SIRT4 | 98,34753543 | 1,075089499 | 0,227828607 | 4,733254327 | 2,21E-06 | 4,07E-05 |
| AP003068.2 | 69,49145467 | 1,045748728 | 0,221320079 | 4,731441888 | 2,23E-06 | 4,09E-05 |
| MTSS1 | 2929,102962 | 1,654146357 | 0,351096483 | 4,727358026 | 2,27E-06 | 4,15E-05 |
| AL590560.2 | 120,164635 | 1,814906046 | 0,398136941 | 4,708077351 | 2,50E-06 | 4,48E-05 |
| AC124016.2 | 116,854978 | 1,002133543 | 0,214280142 | 4,704480502 | 2,55E-06 | 4,54E-05 |
| AC025279.1 | 104,2758127 | 1,381314656 | 0,294837335 | 4,703887732 | 2,55E-06 | 4,55E-05 |
| AL773545.3 | 27,83370224 | 1,160252068 | 0,246928811 | 4,693325472 | 2,69E-06 | 4,74E-05 |
| PSD2 | 17,42910173 | 1,849647571 | 0,390947834 | 4,681260096 | 2,85E-06 | 4,95E-05 |
| AC023794.4 | 11,47496082 | 1,601383024 | 0,339309844 | 4,680697826 | 2,86E-06 | 4,95E-05 |
| COLGALT2 | 870,3854753 | 1,353701533 | 0,298558899 | 4,673802619 | 2,96E-06 | 5,05E-05 |
| ATG9B | 111,1433503 | 1,447918986 | 0,316929422 | 4,673272382 | 2,96E-06 | 5,05E-05 |
| FCGBP | 212,86402 | 1,405748668 | 0,304885636 | 4,664369744 | 3,10E-06 | 5,24E-05 |
| LINC00954 | 421,6966756 | 1,163749282 | 0,248363315 | 4,660070335 | 3,16E-06 | 5,31E-05 |
| SULF2 | 74,31806132 | 2,635487063 | 0,508138654 | 4,656477327 | 3,22E-06 | 5,39E-05 |
| PROX2 | 60,31267389 | 1,344959973 | 0,29151711 | 4,63502145 | 3,57E-06 | 5,86E-05 |
| HSPA7 | 98,06520416 | 2,172463402 | 0,47461447 | 4,6257089 | 3,73E-06 | 6,05E-05 |
| AC092070.2 | 271,9266929 | 1,044125945 | 0,225528349 | 4,625397712 | 3,74E-06 | 6,05E-05 |
| IFT172 | 739,4401397 | 1,046835161 | 0,226649969 | 4,61768802 | 3,88E-06 | 6,22E-05 |
| LRRN1 | 199,3817219 | 1,67634591 | 0,347861825 | 4,613286527 | 3,96E-06 | 6,30E-05 |
| AC107884.1 | 149,9546605 | 1,098534222 | 0,238546131 | 4,607222473 | 4,08E-06 | 6,43E-05 |
| LRFN2 | 17,33028618 | 2,70347866 | 0,589952673 | 4,602924909 | 4,17E-06 | 6,54E-05 |
| LINC00891 | 99,51639127 | 1,211701174 | 0,27010786 | 4,596032167 | 4,31E-06 | 6,73E-05 |
| FCGRT | 188,1205837 | 1,714697178 | 0,365707243 | 4,593517411 | 4,36E-06 | 6,79E-05 |
| LINC-PINT | 939,3163 | 1,052297189 | 0,231429274 | 4,586378535 | 4,51E-06 | 6,97E-05 |

| | | | | | | |
|---|---|---|---|---|---|---|
| UNC45B | 67,03005005 | 1,69186889 | 0,370619754 | 4,578527977 | 4,68E-06 | 7,19E-05 |
| FCGR2B | 178,910675 | 2,286289763 | 0,494827636 | 4,573149683 | 4,80E-06 | 7,34E-05 |
| AC010883.2 | 20,891705 | 1,984927898 | 0,446983453 | 4,562862989 | 5,05E-06 | 7,61E-05 |
| OPRD1 | 7,304310896 | 2,099294135 | 0,457757727 | 4,558500739 | 5,15E-06 | 7,75E-05 |
| GPRASP1 | 292,3099084 | 1,65327553 | 0,351655055 | 4,558420211 | 5,15E-06 | 7,75E-05 |
| PRSS40A | 19,76720964 | 2,409698917 | 0,534900922 | 4,549321221 | 5,38E-06 | 8,02E-05 |
| AC211476.12 | 52,38657243 | 1,113237922 | 0,242282423 | 4,548969966 | 5,39E-06 | 8,02E-05 |
| RTCA-AS1 | 86,85366497 | 1,308702077 | 0,290320204 | 4,545842086 | 5,47E-06 | 8,10E-05 |
| CCDC114 | 62,46447889 | 1,596195989 | 0,350021629 | 4,545020059 | 5,49E-06 | 8,12E-05 |
| AL353748.3 | 22,42083074 | 1,620815243 | 0,371708927 | 4,53980668 | 5,63E-06 | 8,27E-05 |
| AC015911.10 | 459,6133838 | 1,073288441 | 0,239562632 | 4,536841491 | 5,71E-06 | 8,35E-05 |
| AP000977.1 | 104,011099 | 1,901781356 | 0,427486371 | 4,535458552 | 5,75E-06 | 8,40E-05 |
| P2RY6 | 102,6792379 | 2,707602737 | 0,563857161 | 4,529894188 | 5,90E-06 | 8,57E-05 |
| FRAT1 | 637,5847984 | 1,044245964 | 0,230046085 | 4,523603261 | 6,08E-06 | 8,78E-05 |
| MAN1B1-DT | 90,43292212 | 1,17205302 | 0,266235582 | 4,521034025 | 6,15E-06 | 8,87E-05 |
| DLG4 | 58,07181287 | 1,200925482 | 0,267174539 | 4,506728855 | 6,58E-06 | 9,29E-05 |
| KRT2 | 24,47030025 | 1,962531153 | 0,45089209 | 4,503145747 | 6,70E-06 | 9,42E-05 |
| CD68 | 340,2825221 | 2,88669529 | 0,563558472 | 4,496760346 | 6,90E-06 | 9,66E-05 |
| AC015911.8 | 32,80239405 | 1,306274083 | 0,294328639 | 4,493626462 | 7,00E-06 | 9,77E-05 |
| TLR8 | 63,06937408 | 2,818121661 | 0,544355561 | 4,493254349 | 7,01E-06 | 9,78E-05 |
| ZNF540 | 192,2027719 | 1,014889414 | 0,226165718 | 4,492982309 | 7,02E-06 | 9,79E-05 |
| AC114490.1 | 68,7453404 | 1,42591301 | 0,315481587 | 4,492207674 | 7,05E-06 | 9,81E-05 |
| ZEB2 | 5870,147002 | 1,905679567 | 0,428102631 | 4,490713028 | 7,10E-06 | 9,85E-05 |
| SGSM1 | 415,2285547 | 1,377020187 | 0,307687888 | 4,487318345 | 7,21E-06 | 9,94E-05 |
| HELZ2 | 1472,231384 | 1,081424223 | 0,242364264 | 4,476578523 | 7,58E-06 | 0,000102887 |
| AC010175.1 | 27,9370756 | 1,801279253 | 0,435967861 | 4,468299007 | 7,88E-06 | 0,000105815 |
| AZIN2 | 87,19863322 | 1,544194386 | 0,352785449 | 4,467055383 | 7,93E-06 | 0,000106216 |
| C11orf21 | 3310,830889 | 1,394613926 | 0,312979942 | 4,45381841 | 8,44E-06 | 0,00011133 |

| | | | | | | |
|---|---|---|---|---|---|---|
| KLRC4 | 422,5199519 | 1,085847636 | 0,246319418 | 4,448547078 | 8,65E-06 | 0,000113795 |
| AC008750.7 | 26,8318274 | 1,615791861 | 0,364256153 | 4,44163176 | 8,93E-06 | 0,000116874 |
| SERPINA1 | 465,6871084 | 2,858234421 | 0,560442559 | 4,440305821 | 8,98E-06 | 0,00011723 |
| AC022182.2 | 150,9571258 | 1,087910973 | 0,246185903 | 4,437422784 | 9,10E-06 | 0,000118576 |
| RNF43 | 227,8255452 | 1,259155595 | 0,285216739 | 4,43645344 | 9,15E-06 | 0,000119033 |
| KLRC4-KLRK1 | 1187,083535 | 1,055601882 | 0,244961622 | 4,435937577 | 9,17E-06 | 0,00011924 |
| LINC00612 | 9,628627634 | 2,245717697 | 0,47550919 | 4,434097608 | 9,25E-06 | 0,000120027 |
| ZNF600 | 6875,594315 | 1,157301054 | 0,259148588 | 4,428094487 | 9,51E-06 | 0,000122773 |
| PIK3IP1 | 3313,24148 | 1,549054891 | 0,352505797 | 4,42026686 | 9,86E-06 | 0,000126646 |
| SMTNL1 | 26,20968107 | 1,206617562 | 0,272854977 | 4,419653143 | 9,89E-06 | 0,000126924 |
| AC010247.2 | 108,6006819 | 1,751829787 | 0,408572456 | 4,416610391 | 1,00E-05 | 0,000128474 |
| FPR2 | 32,26783206 | 2,812957924 | 0,586331833 | 4,414977997 | 1,01E-05 | 0,000129113 |
| TBKBP1 | 190,9601071 | 1,155598435 | 0,269633612 | 4,410544963 | 1,03E-05 | 0,000131278 |
| FBXO32 | 634,638113 | 1,334966847 | 0,300758638 | 4,409183759 | 1,04E-05 | 0,000131915 |
| EZH1 | 2760,488621 | 1,024724935 | 0,233506286 | 4,398972741 | 1,09E-05 | 0,000136894 |
| MST1P2 | 44,49123567 | 1,487308001 | 0,338995118 | 4,391979895 | 1,12E-05 | 0,000140659 |
| CERCAM | 358,6930088 | 2,05338782 | 0,465546825 | 4,369907416 | 1,24E-05 | 0,000152669 |
| ZNNT1 | 14,06611698 | 1,666753235 | 0,384730813 | 4,365328721 | 1,27E-05 | 0,000155134 |
| HHIPL1 | 12,29385891 | 2,440397057 | 0,586643925 | 4,36195048 | 1,29E-05 | 0,000157065 |
| AGBL2 | 172,1027624 | 1,091825321 | 0,251245593 | 4,342605766 | 1,41E-05 | 0,000168555 |
| AJM1 | 493,903758 | 1,166318095 | 0,270096934 | 4,337255959 | 1,44E-05 | 0,000172086 |
| MMP23B | 565,0281546 | 1,426836318 | 0,330894457 | 4,336378185 | 1,45E-05 | 0,000172493 |
| ACRBP | 71,76783533 | 1,5565221 | 0,354867671 | 4,335078283 | 1,46E-05 | 0,000173278 |
| LILRA1 | 34,58448064 | 2,671567669 | 0,590158482 | 4,33373998 | 1,47E-05 | 0,000173918 |
| AC015813.2 | 194,8450986 | 1,409014415 | 0,321690641 | 4,333027371 | 1,47E-05 | 0,000174273 |
| TNNT3 | 49,39452397 | 1,804851787 | 0,41390741 | 4,329582053 | 1,49E-05 | 0,000176365 |
| ARHGAP8 | 41,14702571 | 1,869067694 | 0,391968748 | 4,328443884 | 1,50E-05 | 0,000176883 |
| AC087500.1 | 71,25157823 | 1,36274409 | 0,313124684 | 4,328552407 | 1,50E-05 | 0,000176883 |

| | | | | | | |
|---|---|---|---|---|---|---|
| SIGLEC14 | 32,97944356 | 2,867917613 | 0,586000779 | 4,314342775 | 1,60E-05 | 0,0001858 |
| GIPR | 247,2273223 | 1,265795748 | 0,294605051 | 4,311569226 | 1,62E-05 | 0,000187271 |
| ITPKB-IT1 | 11,65616568 | 1,94870385 | 0,442908753 | 4,304491259 | 1,67E-05 | 0,000192127 |
| AL031432.2 | 48,63230437 | 1,149306495 | 0,267555358 | 4,303230646 | 1,68E-05 | 0,000192889 |
| FCN1 | 240,7170617 | 2,239097406 | 0,572694685 | 4,300606497 | 1,70E-05 | 0,000194625 |
| AC241377.4 | 88,72225019 | 1,163268268 | 0,257997161 | 4,29405576 | 1,75E-05 | 0,000199312 |
| SMARCD3 | 82,5657991 | 1,061315237 | 0,245133676 | 4,290751423 | 1,78E-05 | 0,000201632 |
| CHI3L1 | 105,0187819 | 2,764764411 | 0,561411352 | 4,290005909 | 1,79E-05 | 0,000202171 |
| CXCL9 | 1631,844563 | 3,019144376 | 0,567702029 | 4,286376323 | 1,82E-05 | 0,000205034 |
| CCL4L2 | 2083,779693 | 1,332132681 | 0,317476168 | 4,280361404 | 1,87E-05 | 0,000209541 |
| AL109955.1 | 56,45528254 | 1,22339593 | 0,287083275 | 4,280156044 | 1,87E-05 | 0,000209541 |
| AC012020.1 | 150,8591323 | 1,13377389 | 0,262738046 | 4,279761268 | 1,87E-05 | 0,000209791 |
| AC117503.5 | 55,09409353 | 1,086407585 | 0,258588869 | 4,27956127 | 1,87E-05 | 0,00020986 |
| CLEC10A | 73,08940149 | 2,590400054 | 0,552820393 | 4,271991036 | 1,94E-05 | 0,000215653 |
| DLEC1 | 189,4312435 | 1,269736394 | 0,295617882 | 4,270105333 | 1,95E-05 | 0,000217241 |
| LAX1 | 1875,573381 | 1,124027965 | 0,265052092 | 4,269530701 | 1,96E-05 | 0,000217679 |
| MTMR9LP | 120,9932721 | 1,196541494 | 0,279272188 | 4,250639616 | 2,13E-05 | 0,000233348 |
| AC008555.1 | 65,24832511 | 1,529956752 | 0,355594055 | 4,24861174 | 2,15E-05 | 0,00023508 |
| AL031432.4 | 52,55290791 | 1,406199828 | 0,328327625 | 4,248006873 | 2,16E-05 | 0,000235456 |
| ERBB2 | 2779,98432 | 1,771240615 | 0,422205266 | 4,237211366 | 2,26E-05 | 0,00024371 |
| IFIT3 | 988,5316983 | 1,879000492 | 0,446826736 | 4,225013506 | 2,39E-05 | 0,000253305 |
| ST3GAL5-AS1 | 15,98539508 | 1,361023574 | 0,322771102 | 4,225133523 | 2,39E-05 | 0,000253305 |
| AC015911.7 | 168,4716196 | 1,194825632 | 0,286871494 | 4,221939215 | 2,42E-05 | 0,000255829 |
| NLRC4 | 20,05947594 | 2,21351362 | 0,491287507 | 4,221693233 | 2,42E-05 | 0,000255972 |
| C1QB | 392,594276 | 2,765195687 | 0,57671314 | 4,217893765 | 2,47E-05 | 0,000259494 |
| NSG1 | 213,3118137 | 2,151324676 | 0,465034021 | 4,200775413 | 2,66E-05 | 0,000275964 |
| SBK1 | 2498,439639 | 1,486623003 | 0,352244724 | 4,198557899 | 2,69E-05 | 0,000278098 |
| CETP | 25,78379367 | 1,520827665 | 0,34856314 | 4,197411794 | 2,70E-05 | 0,000279071 |

| | | | | | | |
|---|---|---|---|---|---|---|
| MORN3 | 119,5276288 | 1,181763759 | 0,283665883 | 4,196401452 | 2,71E-05 | 0,000280027 |
| SNX29P2 | 64,91047058 | 1,145671423 | 0,271929173 | 4,192631513 | 2,76E-05 | 0,000283689 |
| AC022382.2 | 13,66233815 | 1,420709546 | 0,337932477 | 4,192213526 | 2,76E-05 | 0,000283842 |
| U73169.1 | 44,52090523 | 1,071515801 | 0,256254032 | 4,191664825 | 2,77E-05 | 0,000284017 |
| OR52N4 | 23,66026102 | 1,561418814 | 0,377858277 | 4,175564778 | 2,97E-05 | 0,000300827 |
| NF1 | 645,8109695 | 1,044720041 | 0,252128524 | 4,164946871 | 3,11E-05 | 0,000311834 |
| ZNF528-AS1 | 80,98846976 | 1,268779464 | 0,305086055 | 4,161081028 | 3,17E-05 | 0,000315726 |
| C9orf163 | 18,5394897 | 1,187612285 | 0,286313822 | 4,159935506 | 3,18E-05 | 0,000316677 |
| TET1 | 95,91593596 | 1,087796479 | 0,269314783 | 4,154274202 | 3,26E-05 | 0,000323323 |
| CEBPA | 273,3535858 | 1,693595605 | 0,383895284 | 4,152059581 | 3,29E-05 | 0,000325817 |
| LGALS2 | 79,70384392 | 3,037782072 | 0,581743211 | 4,146918286 | 3,37E-05 | 0,0003314 |
| SIRPA | 69,69050234 | 2,514324763 | 0,548528424 | 4,145239844 | 3,39E-05 | 0,000333341 |
| PLXDC1 | 148,4655896 | 1,737290221 | 0,37078693 | 4,143083583 | 3,43E-05 | 0,000335165 |
| MAFB | 77,03530969 | 2,44252124 | 0,582044745 | 4,14144348 | 3,45E-05 | 0,000337238 |
| MARCKS | 50,14277391 | 2,558047711 | 0,565758684 | 4,132939356 | 3,58E-05 | 0,000347048 |
| LINC02384 | 974,945981 | 1,646170751 | 0,423241969 | 4,128697116 | 3,65E-05 | 0,000351807 |
| AL160313.2 | 18,36002482 | 1,9082612 | 0,443445282 | 4,127559352 | 3,67E-05 | 0,000353379 |
| TCAF2P1 | 90,61487765 | 1,196699269 | 0,344802197 | 4,118903513 | 3,81E-05 | 0,000364591 |
| PER3 | 406,3538786 | 1,176375143 | 0,286096826 | 4,117082009 | 3,84E-05 | 0,000367152 |
| ARHGEF25 | 12,10795433 | 1,523866138 | 0,369421632 | 4,11067572 | 3,95E-05 | 0,000375858 |
| AC010894.5 | 15,68235509 | 1,429634561 | 0,352352394 | 4,104378682 | 4,05E-05 | 0,000384238 |
| AC138150.2 | 20,99619197 | 1,611050011 | 0,395209006 | 4,096971637 | 4,19E-05 | 0,000394637 |
| AC008033.3 | 24,92576769 | 1,644614108 | 0,439318701 | 4,090677198 | 4,30E-05 | 0,000403584 |
| S1PR1 | 5902,533667 | 1,650275277 | 0,405233926 | 4,089638087 | 4,32E-05 | 0,000404631 |
| AC010332.3 | 220,6792336 | 1,18054779 | 0,287886468 | 4,087329077 | 4,36E-05 | 0,000408292 |
| AL732406.1 | 25,77180853 | 2,525015641 | 0,595819635 | 4,082382507 | 4,46E-05 | 0,000416102 |
| ZEB2-AS1 | 20,92822888 | 2,041743092 | 0,543004141 | 4,078380522 | 4,54E-05 | 0,000421781 |
| A2M | 46,10435988 | 1,855720802 | 0,465311273 | 4,077395203 | 4,55E-05 | 0,000422936 |

| | | | | | | |
|---|---|---|---|---|---|---|
| C1QA | 275,1619046 | 2,666743726 | 0,577682916 | 4,072474985 | 4,65E-05 | 0,000430965 |
| ALDH2 | 138,8625881 | 2,151838446 | 0,478658617 | 4,07101691 | 4,68E-05 | 0,00043319 |
| LINC00469 | 44,735068 | 2,329501796 | 0,595299585 | 4,066003525 | 4,78E-05 | 0,000440019 |
| ABTB1 | 3730,092258 | 1,015826688 | 0,250840392 | 4,060212195 | 4,90E-05 | 0,000450033 |
| TGFBI | 240,632952 | 2,378503177 | 0,564127078 | 4,04868559 | 5,15E-05 | 0,000466731 |
| HEXD-IT1 | 99,19477023 | 1,606417503 | 0,393003783 | 4,047096326 | 5,19E-05 | 0,000468839 |
| SECTM1 | 210,7848666 | 2,539037872 | 0,566820489 | 4,038806098 | 5,37E-05 | 0,00048264 |
| FCGR1B | 32,76756643 | 2,529744718 | 0,598428639 | 4,038521574 | 5,38E-05 | 0,000483007 |
| HCK | 238,8229275 | 2,62929151 | 0,577984795 | 4,036712715 | 5,42E-05 | 0,000485864 |
| AL627422.2 | 7,307006393 | 2,088340758 | 0,579863734 | 4,033784066 | 5,49E-05 | 0,000491075 |
| TMEM132C | 15,95285903 | 2,113240572 | 0,570097812 | 4,028123874 | 5,62E-05 | 0,000499218 |
| INGX | 56,6903233 | 1,013427798 | 0,252357648 | 4,024988721 | 5,70E-05 | 0,000503984 |
| AC011468.5 | 33,83511075 | 1,00624306 | 0,25045443 | 4,02197368 | 5,77E-05 | 0,000508793 |
| CCL5 | 26076,1303 | 1,350996157 | 0,338240824 | 4,018892107 | 5,85E-05 | 0,000513853 |
| DNAH10OS | 40,39125171 | 1,458902435 | 0,369652043 | 4,017319822 | 5,89E-05 | 0,00051619 |
| LITAF | 23187,8389 | 1,162951381 | 0,291672676 | 4,00566205 | 6,18E-05 | 0,000536211 |
| AC006252.1 | 52,37109194 | 1,009691775 | 0,250258005 | 4,004858797 | 6,21E-05 | 0,000537282 |
| CTSL | 74,21962201 | 2,454853756 | 0,540016838 | 4,001871625 | 6,28E-05 | 0,000542219 |
| Z97989.1 | 67,93343703 | 1,135059575 | 0,283732596 | 3,995354494 | 6,46E-05 | 0,00055494 |
| SNPH | 180,8298809 | 1,253589867 | 0,315616339 | 3,994005513 | 6,50E-05 | 0,000557626 |
| LINC00896 | 28,77510339 | 1,411251209 | 0,353359802 | 3,991520666 | 6,57E-05 | 0,000562285 |
| PLBD1 | 20,7191469 | 2,333156757 | 0,597824193 | 3,990912588 | 6,58E-05 | 0,000563 |
| AC009093.8 | 58,26159984 | 2,08156925 | 0,511004232 | 3,989005573 | 6,64E-05 | 0,000566811 |
| SYNGR1 | 567,9251608 | 1,927252588 | 0,483645475 | 3,98663967 | 6,70E-05 | 0,000571506 |
| AC099521.3 | 4,465709563 | 1,915216771 | 0,507221182 | 3,985714789 | 6,73E-05 | 0,00057349 |
| LINC00243 | 64,48078132 | 1,561970772 | 0,389064624 | 3,985374808 | 6,74E-05 | 0,000574066 |
| AC092636.2 | 7,528712191 | 2,192861718 | 0,592925011 | 3,98377413 | 6,78E-05 | 0,000577452 |
| AC002316.1 | 1373,011225 | 1,781229551 | 0,444075125 | 3,979749852 | 6,90E-05 | 0,000585806 |

| | | | | | | |
|---|---|---|---|---|---|---|
| AMY2B | 265,8743418 | 1,086094794 | 0,269209238 | 3,977767361 | 6,96E-05 | 0,000589217 |
| EFNA1 | 14,24715828 | 1,423949984 | 0,353903706 | 3,976513335 | 6,99E-05 | 0,000591301 |
| AL592295.4 | 142,3120406 | 1,031011265 | 0,262967147 | 3,973726899 | 7,08E-05 | 0,000595923 |
| LSMEM1 | 18,07762195 | 1,192849374 | 0,300746055 | 3,970909637 | 7,16E-05 | 0,000601029 |
| CD4 | 240,2007002 | 2,606279363 | 0,593337825 | 3,96790605 | 7,25E-05 | 0,000606599 |
| C1QC | 207,6228252 | 2,627950954 | 0,578340915 | 3,966688121 | 7,29E-05 | 0,000608936 |
| AGPAT4 | 825,0728962 | 1,142423688 | 0,292463785 | 3,951534591 | 7,77E-05 | 0,000641791 |
| NPC1 | 6856,804572 | 1,1169577 | 0,283898687 | 3,951619429 | 7,76E-05 | 0,000641791 |
| SSBP3-AS1 | 83,55414745 | 1,35851243 | 0,348467799 | 3,949429756 | 7,83E-05 | 0,000645846 |
| SPTBN5 | 56,12225854 | 1,790237666 | 0,449479988 | 3,94387235 | 8,02E-05 | 0,000658542 |
| LRRC25 | 99,37734494 | 2,366414987 | 0,554275373 | 3,943665224 | 8,02E-05 | 0,000658838 |
| MGAM2 | 16,39325032 | 2,038434496 | 0,600942011 | 3,936225634 | 8,28E-05 | 0,000675674 |
| LIPC | 100,0814985 | 1,466979253 | 0,372709729 | 3,932402949 | 8,41E-05 | 0,000684541 |
| PRR5L | 5668,248956 | 1,120823026 | 0,286527703 | 3,922323325 | 8,77E-05 | 0,000708018 |
| LYZ | 2603,852795 | 2,274736707 | 0,559041505 | 3,921639446 | 8,79E-05 | 0,000709165 |
| AL445228.3 | 17,73035404 | 1,353675163 | 0,341398838 | 3,910800302 | 9,20E-05 | 0,000734566 |
| CCDC183 | 36,65153314 | 1,398895017 | 0,37414863 | 3,902339346 | 9,53E-05 | 0,000753179 |
| LINC00921 | 253,0803091 | 1,17283326 | 0,298308603 | 3,900763022 | 9,59E-05 | 0,000757195 |
| AC106782.1 | 18,21332579 | 1,688685299 | 0,428966603 | 3,89942032 | 9,64E-05 | 0,000759893 |
| PLIN1 | 28,93255006 | 1,540497579 | 0,401704587 | 3,892711527 | 9,91E-05 | 0,000778133 |
| AC108704.2 | 273,4213432 | 1,200214784 | 0,31201904 | 3,891357084 | 9,97E-05 | 0,000781872 |
| LILRB2 | 146,8986618 | 2,51266224 | 0,5753606 | 3,87920882 | 0,000104797 | 0,000813637 |
| FAM110C | 211,470031 | 1,787405297 | 0,45720726 | 3,879197894 | 0,000104801 | 0,000813637 |
| COL6A4P2 | 11,74889994 | 1,809747089 | 0,570450663 | 3,86995067 | 0,000108857 | 0,000839866 |
| AC092117.1 | 95,12539326 | 1,010526045 | 0,260581465 | 3,861426274 | 0,000112727 | 0,000863341 |
| AC107375.1 | 27,79797232 | 1,208929691 | 0,315845007 | 3,849790212 | 0,000118219 | 0,000896747 |
| H2AC3P | 7,187114823 | 1,946149143 | 0,495613703 | 3,846985969 | 0,00011958 | 0,000904992 |
| FCGR1A | 120,6255376 | 2,35160559 | 0,598788438 | 3,846140739 | 0,000119993 | 0,000907772 |

| | | | | | | |
|---|---|---|---|---|---|---|
| CARNS1 | 481,2979576 | 1,02262037 | 0,26340231 | 3,842597591 | 0,000121739 | 0,00091818 |
| MAP3K6 | 269,7151428 | 1,021393561 | 0,265297573 | 3,837253772 | 0,000124418 | 0,000934477 |
| AC067945.4 | 14,57270036 | 1,306885021 | 0,34569316 | 3,83336978 | 0,0001264 | 0,000946495 |
| ITGA5 | 7566,222291 | 1,017538868 | 0,267460512 | 3,82982707 | 0,000128233 | 0,000956795 |
| HSD11B1-AS1 | 10,3404823 | 1,498716783 | 0,39565234 | 3,829069372 | 0,000128629 | 0,000958948 |
| AC009951.4 | 24,76922044 | 1,812496874 | 0,465994527 | 3,826523032 | 0,000129966 | 0,000965545 |
| ADAMTS6 | 6,127241271 | 2,131827374 | 0,582871238 | 3,824732893 | 0,000130914 | 0,000971133 |
| SERPING1 | 254,6799424 | 2,48752146 | 0,58399494 | 3,820698331 | 0,000133074 | 0,000984581 |

**Supplementary table 4.** List of CYTOKINE_Bright signature genes used in this study, Related to Fig. 3D, 5B

| Name | Gene symbols |
|---|---|
| CYTOKINE_Bright signature | *IFNG, TNFA, CSF2, IL6, XCL1, XCL2* |

**Supplementary table 5.** List of CYTOKINE_Dim signature genes used in this study, Related to Fig. 3D, 5B

| Name | Gene symbols |
|---|---|
| CYTOKINE_Dim signature | *CCL3, CCL4, CCL5, IL8* |

**Supplementary table 6.** List of HOMING_Bright signature genes used in this study, Related to Fig. 3D, 5B

| Name | Gene symbols |
|---|---|
| HOMING_Bright signature | *CCR7, CXCR3, CCR1, CCR5, CCR6, SELL* |

**Supplementary table 7.** List of HOMING_Dim signature genes used in this study, Related to Fig. 3D, 5B

| Name | Gene symbols |
|---|---|
| HOMING_Dim signature | *S1PR5, CX3CR1, CXCR1, CXCR2, CXCR4* |

**Supplementary table 8.** List of NK cell signature genes used in this study (from (41)), Related to Fig. 6B-E, Supplementary Fig. S13A, S13A

| Name | Gene symbols |
|---|---|
| NK cell signature | *NCR3, KLRB1, PRF1, CD160, NCR1* |

**Supplementary table 9.** Research Resource Identifier

| REAGENT OR RESOURCE | SOURCE | IDENTIFIER |
|---|---|---|
| **Flow cytometry antibodies (Human)** | | |
| Anti-CD3-BV711 | BD Bioscience | Clone UCHT1 Mouse IgG1 κ |
| Anti-CD3-PerCP | Miltenyi | Clone BW264/56, Mouse IgG2a κ |
| Anti-CD3-V450 | BD Bioscience | Clone UCHT1 Mouse IgG1 κ |
| Anti-CD4-BB515 | BD Bioscience | Clone RPAT4 Mouse IgG1 κ |
| Anti-CD4-FITC | BD Bioscience | Clone RPAT4 Mouse IgG1 κ |
| Anti-CD7-BV650 | BD Bioscience | Clone M-T701, Mouse IgG1 κ |
| Anti-CD7-BV711 | BD Bioscience | Clone M-T701, Mouse IgG1 κ |
| Anti-CD7-FITC | Beckman Coulter | Clone 8h8.1, Mouse IgG2a κ |
| Anti-CD8-PerCP-Cy5.5 | BD Bioscience | Clone RPA T8, Mouse IgG1 κ |
| Anti-CD14-BV510 | BD Bioscience | Clone MϕP9, Mouse IgG2b κ |
| Anti-CD16-V500 | BD Bioscience | Clone 3G8, Mouse IgG1 κ |
| Anti-CD25-PE | BD Bioscience | Clone M-A251, Mouse IgG1 κ |
| Anti-CD45-AF700 | BD Bioscience | Clone HI30, Mouse IgG2b κ |
| Anti-CD45-BV510 | BD Bioscience | Clone HI30, Mouse IgG2b κ |
| Anti-CD56-APC | BD Bioscience | Clone NCAM16.2, Mouse IgG2b κ |
| Anti-CD56-PEVio770 | Miltenyi | Clone AF12-7H3, Mouse IgG1 κ |
| Anti-CD56-VioBright-FITC | Miltenyi | Clone AF12-7H3, Mouse IgG1 κ |
| Anti-CD57-APC | BD Bioscience | Clone HNK-1, Mouse IgM κ |
| Anti-CD69-APC | BD | Clone FN50, Mouse IgG1 κ |
| Anti-CD127-APC | eBioscience | Clone eBioRDR5, Mouse IgG1 κ |
| Anti-CD158(KIR2DL1/S1/S3/S5)-PE-Cy7 | Biolegend | CloneHP-A4, Mouse IgG2b κ |
| Anti-CD158e1(KIR3DL1)-PE-Cy7 | Biolegend | Clone DX9, Mouse IgG1 κ |
| Anti-IFNg-BV421 | Biolegend | Clone 4SB3, Mouse IgG1 κ |
| Anti-IL-1R1-BV421 | BD Bioscience | Clone 89412, Mouse IgG1 κ |
| Anti-IL-18Ra-PE | BD Bioscience | Clone H44, Mouse IgG1 κ |

| | | |
|---|---|---|
| Anti-p-p38-PE | BD Bioscience | Clone 36/p38, Mouse IgG1 κ |
| Anti-p-p65-PE | BD Bioscience | Clone K10-895.12.50, Mouse IgG2b κ |
| Anti-p-S6 | Cell Signaling Technology | Clone D57.2.2E, Rabbit IgG |
| Anti-p-STAT4-PE | BD Bioscience | Clone 38/p-Stat4, Mouse IgG2b κ |
| Anti-ST2-PE | R&D | Polyclonal goat |
| **Flow cytometry antibodies (Mouse)** | | |
| anti-CD11b-APCeFluor 780 | eBioscience | Clone M1/70, Rat IgG2b κ |
| anti-CD27-PerCPeFluor 710 | eBioscience | Clone LG.7F9, Hamster IgG |
| anti-IFN-γ-PE | Biolegend | Clone XMG1.2, Rat IgG1 κ |
| anti-IL-1R-BV421 | BD Bioscience | Clone 4E2, Rat IgG2a κ |
| anti-IL-18Rα-PE | eBioscience | Clone P3TUNYA, Rat IgG2a κ |
| anti-NK1.1-APC | BD Bioscience | Clone PK136, Mouse IgG2a κ |
| anti ST2-PE | BD Bioscience | Clone U29-93, Rat IgG2a κ |
| **Other antibodies** | | |
| Anti-goat | Agilent | E0466 |
| Anti-IL-18 | MBL | Clone 125-2H, Mouse IgG1 κ |
| Anti-IL-33 | R&D | Clone AF3625, Goat |
| Anti-NK1.1 | BioXCell | Clone PK136 |
| Anti-IFN-γ | BioXCell | Clone XMG1.2, Rat IgG1 κ |
| Anti-rat IgG1 isotype control | BioXCell | Clone HRPN |
| Anti-NKp30 | R&D | Clone 210845 Mouse IgG2a κ |
| Anti-NKp46 | R&D | Clone 195314, Mouse IgG2b κ |
| Anti-PanCK | Agilent | clone AE1-AE3, |
| Anti-p-STAT4 | Cell Signaling Technology | Y693, Rabbit IgG |
| Anti-ST2 (blocking) | R&D | Clone 97203, Mouse IgG1 κ |
| **Reagents** | | |
| Calcein | Invitrogen | C1430 |
| CTV | Invitrogen | C34557 |

*Continued*

| REAGENT OR RESOURCE | SOURCE | IDENTIFIER |
|---|---|---|
| DAPI | Invitrogen | D1306 |
| DNAse I | Sigma-Aldrich | D4263 |
| GolgiPlug | BD bioscience | 555029 |
| TotalSeq™-B0251 anti-human Hashtag 1 Antibody | Biolegend | LNH-94; 2M2; Mouse IgG1 κ |
| TotalSeq™-B0252 anti-human Hashtag 2 Antibody | Biolegend | LNH-94; 2M2; Mouse IgG1 κ |
| Lisofylline | Cayman chemicals | 10010785 |
| Lymphocyte Separation Medium | Eurobio | CMSMSL0101 |
| Pharm LyseTM Buffer | BD Biosciences | 555899 |
| Lyse/fix Buffer Phosflow | BD Biosciences | 558049 |
| Perm Buffer III Phosflow | BD Biosciences | 558050 |
| RPMI 1640 Medium, GlutaMAX | Gibco | 61870036 |
| Sulfinpyrazone | Sigma-Aldrich | S9509 |
| Type IV collagenase | Sigma-Aldrich | C5138 |
| Zombie Dye Aqua | Biolegend | 423101 |
| **Cytokines** | | |
| Recombinant Human IFN-α2b | Schering-Plough | |
| Recombinant Human IL-1α | Peprotech | 200-01A |
| Recombinant Human IL-1β | Peprotech | 200-01B |
| Recombinant Human IL-1RA | Peprotech | 200-01RA |
| Recombinant Human IL-2 | Chiron | |
| Recombinant Human IL-12 | Miltenyi | 130-096-704 |
| Recombinant Human IL-15 | Peprotech | 200-15 |
| Recombinant Human IL-18 | MBL | B003-5 |
| Recombinant Human IL-33 | Miltenyi | 130-109-378 |

***Continued***

| **REAGENT OR RESOURCE** | **SOURCE** | **IDENTIFIER** |
|---|---|---|
| Recombinant Mouse IL-1α | Miltenyi | 130-094-050 |
| Recombinant Mouse IL-1β | Miltenyi | 130-094-053 |
| Recombinant Mouse IL-12 | Miltenyi | 130-096-707 |
| Recombinant Mouse IL-12 (*in vivo*) | R&D | 419-ML |
| Recombinant Mouse IL-18 | MBL | B002-5 |
| Recombinant Mouse IL-33 | Miltenyi | 130-112-958 |
| Recombinant Mouse IL-33 (*in vivo*) | Biolegend | 580502 |
| **Kits** | | |
| Bio-Plex Pro™ Human Cytokine 17-plex | BioRad | M5000031YV |
| ChIP-IT High Sensitivity | Active Motif | 53040 |
| Extra Sensitive IFN gamma Mouse ELISA Kit | ThermoFischer | BMS609 |
| FoxP3/Transcription Factor Staining Set | eBioscience | 00-5523-00 |
| Human IFN-γ DuoSet ELISA kit | R&D | DY285B |
| iScript Reverse Transcription kit | BioRad | 1708840 |
| MycoAlertTM Mycoplasma Detection Kit | Lonza | LT07-118 |
| NK Cell Isolation Kit, human | Miltenyi | 130-092-657 |
| NucleoSpin RNA Kit | Macherey-Nagel | 740955 |
| **Oligonucleotides** | | |
| *GADD45a* TaqMan | ThermoFischer | Hs00169255_m1 |
| Negative Primer Set 1 and 2 | Active Motif | 71001 and 71002 |
| *IL1RL1* TaqMan | ThermoFischer | Hs00249384_m1 |
| *IL1RL1* (promoter) Forward | Agilent | 5'-GTGATCATCGGGTTCAGCTTATC-3' |
| *IL1RL1* (promoter) Reverse | Agilent | 5-'GCTTTACTAAATACAACAGCCAGCCT-3' |
| SimpleChIP Human *PRF1* Primers | Cell Signaling Technology | 9014S |

342

**Supplementary table 10.** Summary of data preprocessing and clustering settings for downstream analyses of the validation scRNA-seq dataset (42)

| Downstream analysis | Cell number after filtered | Number of dimensions used | Resolution of granularity | Number of unsupervised clusters | Number of annotated population | Computational method used | Space embedding selected |
|---|---|---|---|---|---|---|---|
| T/NK cells | 71,623 | 20 | 0.3 | 12 | 12 | Harmony | UMAP |
| Cycling cells | 3,270 | 20 | 0.5 | 12 | 10 | Harmony + S/G2M scores regression | UMAP |
| ILC/NK cells | 4,624 | 15 | 0.15 | 5 | 5 | Harmony | UMAP |
| Total NK cells | 4,605 | 15 | 0.15 | 5 | 5 | Harmony + S/G2M scores regression | t-SNE |
